# Transcription factor activities as a shared representation for drug repurposing hypotheses in breast cancer

**DOI:** 10.64898/2026.09.08.750045

**Authors:** Carlos Fernandez-Lozano, David Ferreiro, Patricia V.-del-Río

## Abstract

Drug repurposing could expand the limited targeted options for triple-negative breast cancer (TNBC). PRECISION trains a heterogeneous graph neural network (GNN) over prior-knowledge edges on the PRISM screen, describing cell lines mostly by transcription factor (TF) activities. On held-out cell lines the GNN fell below a drug-mean reference and a TF-based Random Forest. Integrated Gradients attributions define ten core TFs, shared by the drugs examined and partly stable across seeds. Per-drug Ridge models on the same TF activities carried predictions to unselected breast cancer cohorts, where predicted sensitivity was associated with overall survival for 623 of 1,447 drugs in SCAN-B and 74 in METABRIC, partly through a shared prognostic axis. An explicit rule selects seven kinase or histone deacetylase inhibitors whose associations are prognostic and persist only partly within TNBC. Neither TNBC cell lines nor a triple-negative cohort supported them. The shared TF representation yields repurposing hypotheses, not validated treatments.

## Introduction

Triple-negative breast cancer (TNBC) accounts for approximately 10–20% of all breast cancer diagnoses (Chen et al., 2023) and has a poorer prognosis than other subtypes (Bianchini et al., 2016). Defined by the absence of estrogen receptor (ER), progesterone receptor (PR), and human epidermal growth factor receptor 2 (HER2) expression, TNBC lacks the molecular targets that enable endocrine or anti-HER2 therapies in other subtypes. Standard-of-care treatment relies primarily on cytotoxic chemotherapy. Adding pembrolizumab to neoadjuvant chemotherapy improves pathological complete response (Schmid et al., 2020) and event-free survival (Schmid et al., 2022) in early disease, and options for advanced disease now include poly(ADP-ribose) polymerase (PARP) inhibitors for carriers of germline BRCA1/2 mutations (Robson et al., 2017; Litton et al., 2018) and the antibody-drug conjugate sacituzumab govitecan, which is given without biomarker selection (Bardia et al., 2021). In the advanced setting, however, single-agent checkpoint inhibition benefits only a minority of patients, with objective response rates below 25% (Chen et al., 2023). Drug repurposing, the systematic identification of new indications for existing approved compounds (Pushpakom et al., 2019), offers a path forward: it builds on established safety profiles and existing manufacturing pipelines to compress the timeline from discovery to clinical deployment.

The PRISM (Profiling Relative Inhibition Simultaneously in Mixtures) platform provides viability measurements for thousands of compounds (4,518 at a single dose in the primary screen and 1,448 used here from the eight-dose secondary screen) across hundreds of cancer cell lines, with the secondary-screen responses summarized as the area under the dose-response curve (AUC) (Corsello et al., 2020). The Genomics of Drug Sensitivity in Cancer (GDSC) project offers complementary screening data with partially overlapping compound libraries (Yang et al., 2013). Combined with the rich molecular profiling available through the Cancer Dependency Map (DepMap), including gene expression, CRISPR essentiality, and mutation data (Ghandi et al., 2019; Meyers et al., 2017), these resources enable systematic modeling of drug sensitivity as a function of molecular features.

A fundamental limitation of cell-line-based pharmacogenomics is the translation gap: models trained on cancer cell lines frequently fail to generalize to patient tumors (Mourragui et al., 2021; He et al., 2022). Several methods have attempted to bridge this gap. TRANSACT uses kernel-based domain adaptation to identify consensus features between cell lines and patients (Mourragui et al., 2021). CODE-AE separates shared and private embeddings of cell lines and tumors, and aligns the two domains on the concatenated embeddings, adversarially or by maximum mean discrepancy (He et al., 2022). While each of these methods represents a meaningful advance, they share a common limitation: they operate in the space of gene expression, relying on statistical alignment between domains without incorporating biological prior knowledge about the regulatory mechanisms that connect drug targets to transcriptional programs.

Graph neural networks (GNNs) offer a framework for integrating biological prior knowledge into predictive models (Li et al., 2022). DrugCell applied this idea by embedding the Gene Ontology hierarchy into a neural network to predict drug response, achieving both competitive accuracy and biological interpretability (Kuenzi et al., 2020). However, to our knowledge, existing graph-based approaches for drug response prediction have not integrated transcription factor (TF) regulatory networks as a core component of the knowledge graph. This is a relevant gap, because TF regulatory programs represent the mechanistic link between upstream signaling (perturbed by drugs) and downstream transcriptional output (measured by expression profiling).

The use of TF activities as a molecular representation offers a distinct advantage over raw gene expression for pharmacogenomic applications. Garcia-Alonso et al. (2018) showed that TF activities inferred from regulon expression improve 59% (95 of 160) of the known pharmacogenomic marker associations compared with genomic features alone. The decoupleR framework (Badia-i-Mompel et al., 2022) and its curated regulon resource CollecTRI (Müller-Dott et al., 2023) provide a standardized pipeline for computing TF activities from any expression matrix using univariate linear modeling (ULM). TF activities compress thousands of gene expression measurements into a mechanistically grounded lower-dimensional space and may be more robust to the technical variation that separates cell-line from clinical data, although we do not compare them here with transfer based on expression.

In this work, we present PRECISION (Figure 1), a heterogeneous GNN framework that integrates TF regulatory networks from CollecTRI, protein-protein interactions (PPI) from OmniPath (Türei et al., 2021), drug-target annotations, and pharmacogenomic response data from PRISM into a knowledge graph with 23,498 nodes and approximately 830,000 edges, reserving GDSC for cross-screen evaluation. In a cell-line hold-out evaluation, the GNN falls below a Random Forest and a reference that predicts each drug’s mean response. Integrated Gradients (Sundararajan et al., 2017) on the trained model attributes its predictions to individual TFs. These attributions surface a core set of regulators spanning epithelial-mesenchymal transition (EMT), NF-*κ*B, interferon, homeobox developmental, epithelial and stromal programs. Per-drug Ridge models on the TF activities, which the GNN also takes as input, carry predictions to patients. We then test whether predicted drug sensitivity is associated with overall survival in three breast cancer cohorts on two expression platforms, the two larger of which, the Sweden Cancerome Analysis Network-Breast (SCAN-B) and the Molecular Taxonomy of Breast Cancer International Consortium (METABRIC), have more than 1,100 deaths each, and examine the core TFs in the AURORA-US cohort of matched primary and metastatic tumors. The survival cohorts are not selected for TNBC and none of the candidate drugs is a standard breast cancer treatment, so these associations are prognostic, and we report how much of them persists within triple-negative tumors. Finally, we examine the candidates in the TNBC cell lines of PRISM and in FUSCC-TNBC, a triple-negative cohort from the Fudan University Shanghai Cancer Center.

**Figure 1.**
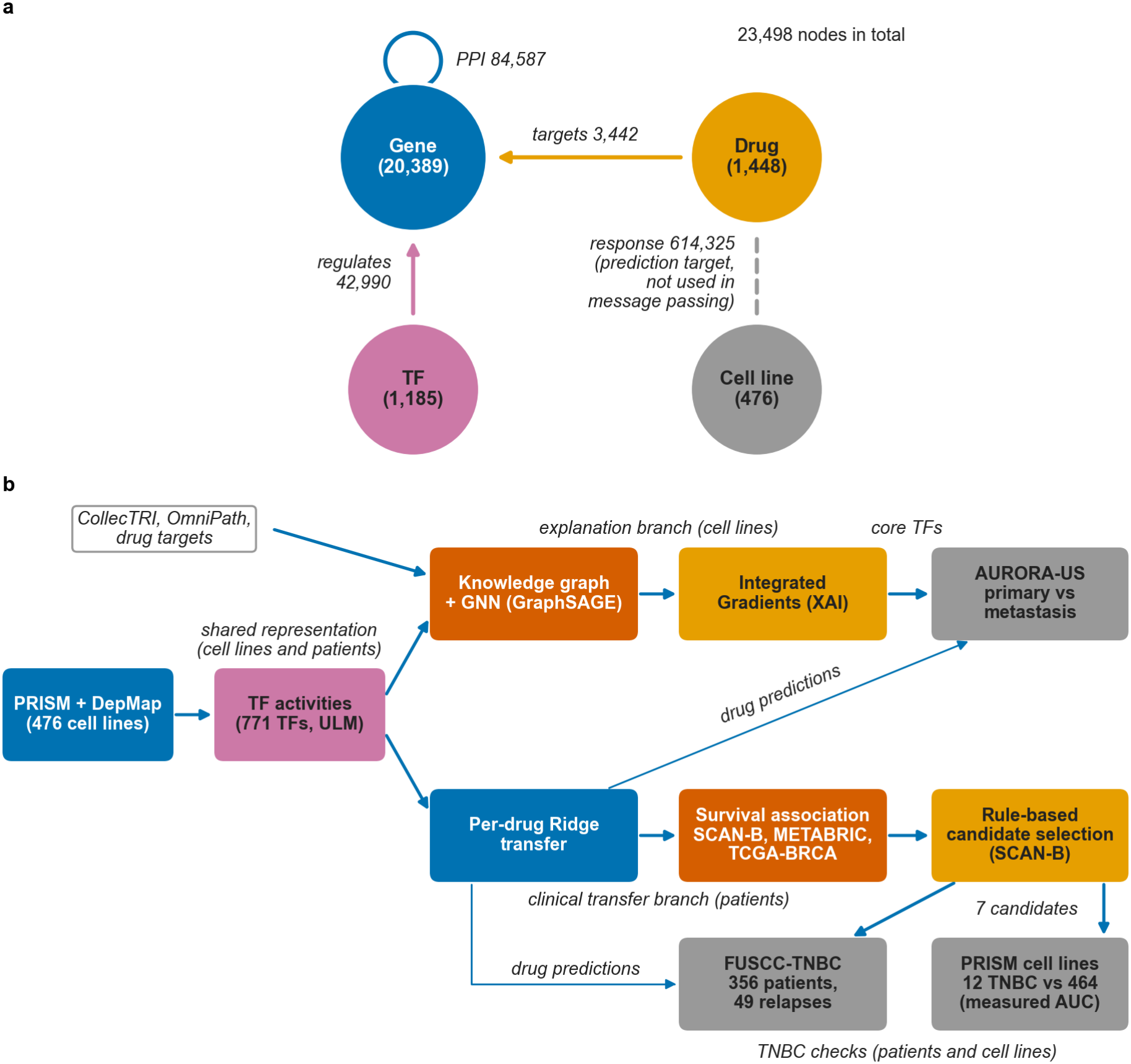
PRECISION framework overview. (**a**) Heterogeneous biomedical knowledge graph: four node types (genes, TFs, drugs, cell lines) connected by four edge types encoding biological prior knowledge (TF regulatory network from CollecTRI, protein-protein interactions from OmniPath, drug-target annotations) and pharmacogenomic observations (drug response AUC). The response edges are the prediction targets and are excluded from message passing. Each of the 84,587 OmniPath interactions is stored in both directions, giving 169,174 directed gene-gene edges over 83,176 distinct gene pairs. (**b**) End-to-end pipeline: pharmacogenomic data (PRISM response, DepMap expression) and prior knowledge are integrated into the knowledge graph, and Integrated Gradients on the trained GNN gives TF-level attributions. In the cell lines, TF activities are computed for the 771 of the 1,185 TFs with at least five measured targets. Per-drug Ridge regression on TF activities transfers predictions to the three cohorts of the survival screen (SCAN-B, METABRIC, TCGA-BRCA), to AURORA-US and to FUSCC-TNBC, and the core TFs from Integrated Gradients are examined in the AURORA-US cohort of matched primary and metastatic tumors. Candidates are selected from the PAM50-adjusted survival associations in SCAN-B by an explicit rule. They are then examined in TNBC: the 12 TNBC cell lines of PRISM are compared with the other 464 on measured AUC, and the candidates are tested in FUSCC-TNBC (356 patients, 49 relapses), where the relapse-free survival associations of all 1,447 drugs are also compared with the overall survival associations of the triple-negative tumors of SCAN-B.

## Results

### Model comparison in the cell-line hold-out

We compared a heterogeneous GNN (GraphSAGE, with a Heterogeneous Graph Transformer, HGT, as a variant) with Ridge regression on expression, a Random Forest on TF activities and a hybrid Ridge model on expression plus GNN cell-line embeddings, all in the same hold-out of 381 training and 95 test cell lines over 1,447 drugs (Figure 2, Table 1). Between-drug differences in mean response explained 59% of the test AUC variance and dominated the global Pearson correlation, so predicting each drug’s mean training AUC, without cell-line information, reached 0.764. GraphSAGE (0.743 *±* 0.003 over five seeds, training curves in Supplementary Figure S1) and Ridge on expression (0.726) fell below this value, and the Random Forest (0.769 over five seeds) barely exceeded it. The median per-drug correlation, which measures how well a model ranks cell lines for a given drug, was low for every model and lowest for the GNNs (Table 1, seed-42 distribution in Supplementary Figure S2). The cell-line embeddings learned by the GNN partly clustered by lineage, for example skin and bowel, and within breast cancer separated TNBC from most ER+ and HER2+ lines (Supplementary Figure S3).

**Table 1.** Model comparison in the cell-line hold-out (381 train and 95 test cell lines), over the same 1,447 drugs for every model. GNN, hybrid and Random Forest rows give the mean *±* standard deviation over five seeds on a single split. Ridge on expression is a single fit. The two reference rows are a drug-mean predictor that ignores the cell line and the per-drug Ridge models on TF activities used for clinical transfer. The hyperparameters of Ridge on expression and the Random Forest were selected by cross-validation within the training lines, whereas those of GraphSAGE, whose optimizer settings HGT also used, were selected on a split that included these test lines (Methods), so the GNN and hybrid rows are optimistic. RF, Random Forest. expr, expression. emb, embeddings.

| Model | Global<br>Pearson | Median<br>per-drug | Drugs<br>>0.3 |
|---|---|---|---|
| RF (TF activities) | 0.769 ± <0.001 | 0.126 ± 0.003 | 14.5% |
| Ridge (expression) | 0.726 | 0.114 | 10.6% |
| GraphSAGE | 0.743 ± 0.003 | 0.032 ± 0.009 | 0.8% |
| HGT | 0.472 ± 0.027 | 0.048 ± 0.012 | 1.2% |
| Ridge (expr + GNN emb) | 0.723 ± <0.001 | 0.112 ± 0.001 | 10.4% |
| *References* |  |  |  |
| Drug mean | 0.764 | – | – |
| Ridge (TF, transfer) | 0.700 | 0.086 | 6.2% |

**Figure 2.**
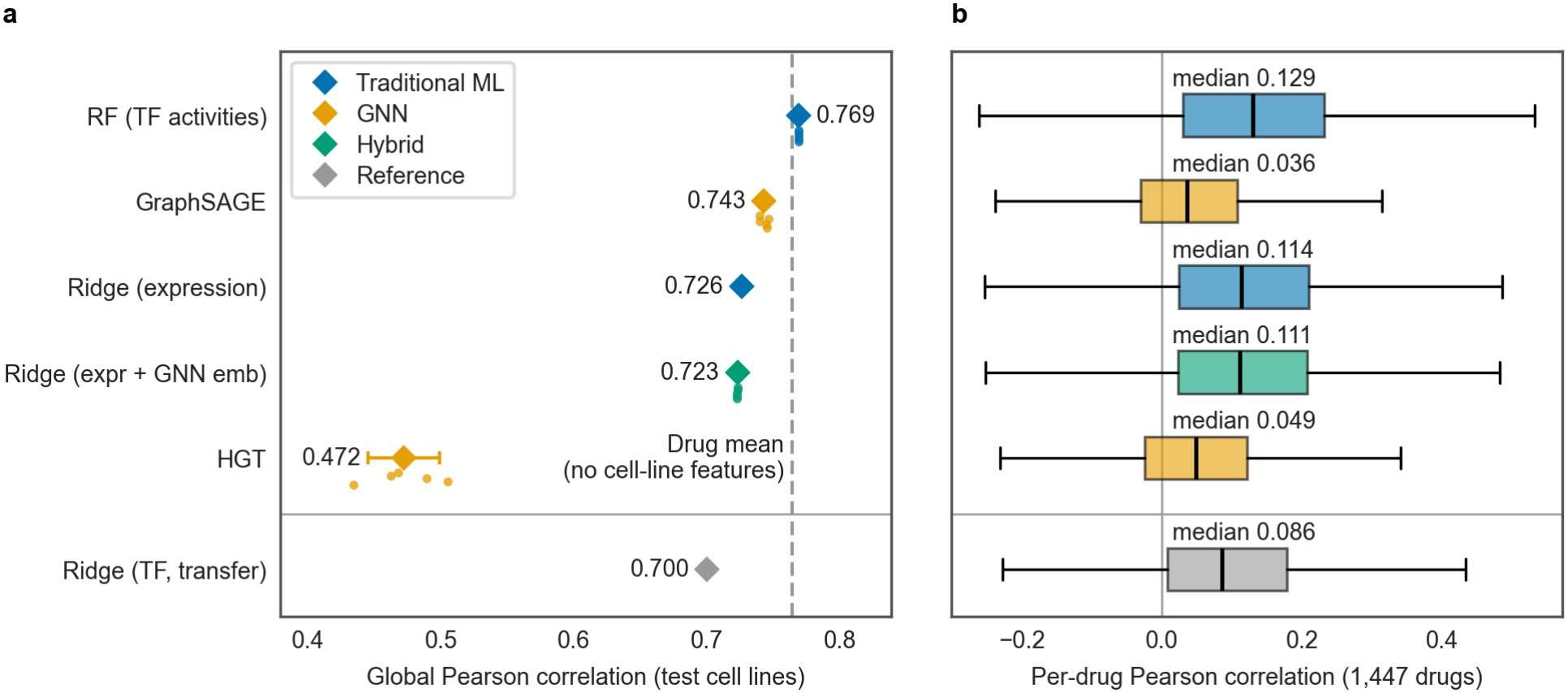
Model comparison in the cell-line hold-out. Blue marks traditional machine-learning models, orange the GNNs, green the hybrid and gray the references. (**a**) Global Pearson correlation over all test pairs: one point per seed for the GNNs, the hybrid and the Random Forest, with mean and standard deviation (diamond and bar, the bar hidden when narrower than the diamond), single fits for the other models, and the drug-mean reference as a dashed line. (**b**) Per-drug Pearson correlation over the 1,447 drugs, with the median above each box. GNN, hybrid and Random Forest boxes show the seed-42 models, whose medians differ slightly from the five-seed means of Table 1. Boxes span the interquartile range with the median as a black line, whiskers extend to 1.5 times the interquartile range, and values beyond them are not drawn.

### TF attributions of the trained GNN

We applied Integrated Gradients (IG) to the trained GNN for the 12 TNBC cell lines paired with 11 drugs, the seven repurposing candidates and four of the five positive controls (paclitaxel, docetaxel, epirubicin and olaparib), to identify the TFs that most influence the model’s predictions (Figure 3).

**Figure 3.**
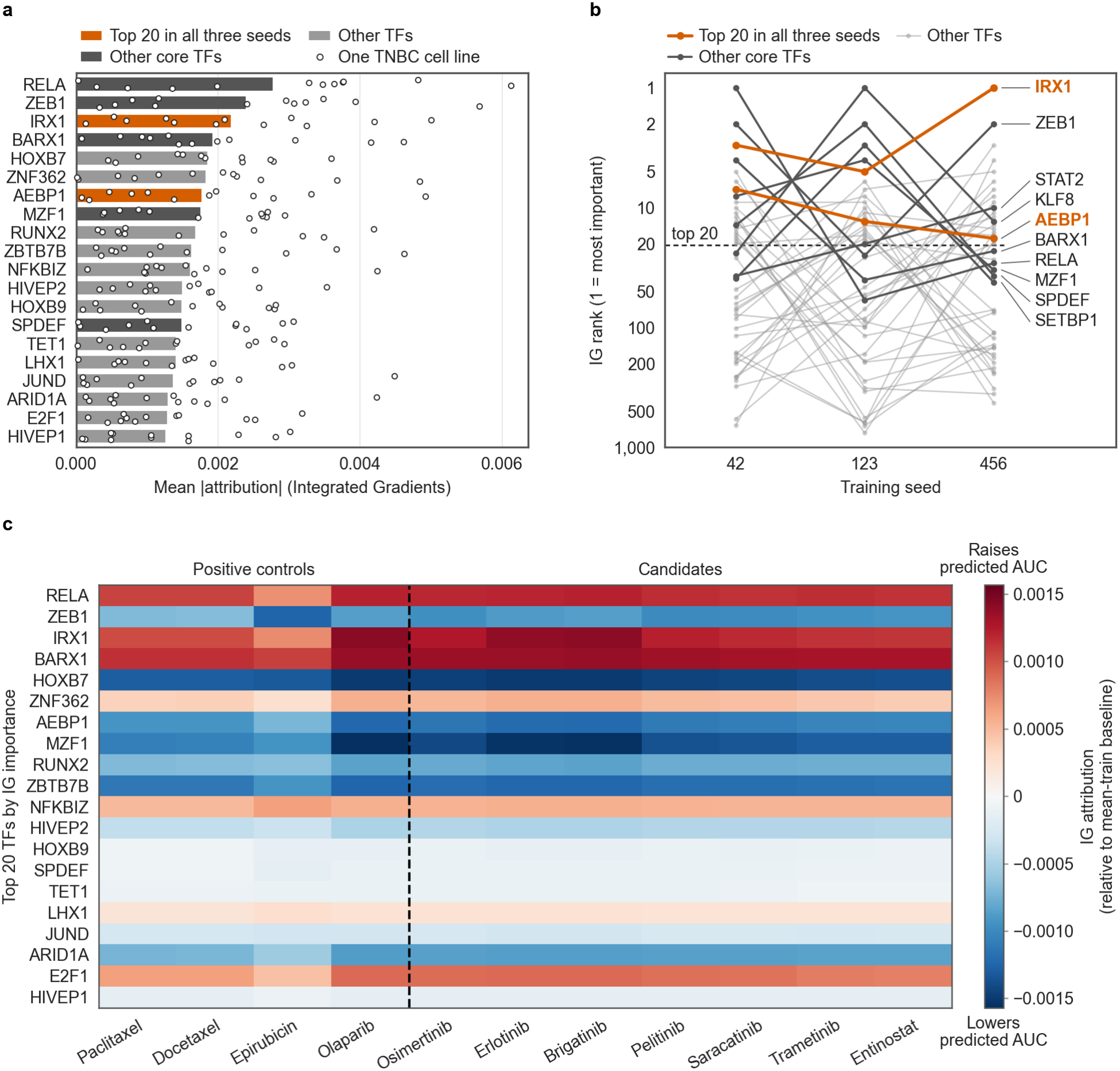
Integrated Gradients attribution on the trained GNN. (**a**) Top 20 TFs by global IG importance (seed-42 model, mean-train baseline, 50 integration steps), the mean absolute attribution over the 12 TNBC cell lines *×* 11 drugs (seven candidate drugs plus four of the five positive controls: paclitaxel, docetaxel, epirubicin, olaparib). Each point is one cell line, with its absolute attribution averaged over the 11 drugs, and each bar is the mean of its 12 points. Colors follow panel b: the two TFs in the top 20 of all three seeds (IRX1, AEBP1) are in vermillion and the other core TFs in dark gray. Seven of the ten core TFs are in this top 20. The other three, KLF8, STAT2 and SETBP1, rank 24, 37 and 39 in the seed-42 model and do not appear in panels a and c. (**b**) Multi-seed stability: the rank obtained with IG on three GNNs trained with seeds 42, 123 and 456, each on its own cell-line split, for the 44 TFs in the top 20 of at least one seed (logarithmic rank axis). The ten core TFs, with the lowest mean rank across seeds, are labeled, and the TFs appearing in the top 20 of all three seeds (IRX1, AEBP1) are highlighted. (**c**) Signed IG attributions, averaged over the 12 cell lines, for the top 20 TFs *×* 11 drugs, relative to the mean-train baseline: red raises and blue lowers the mean predicted AUC of these cell lines. For 7 of the 20 TFs higher activity moves the prediction opposite to the sign shown (main text). The dashed line separates the four positive controls (left) from the seven candidates (right). Panels a and c and the baseline test use the same seed-42 model.

The IG ranking of the model (Figure 3a) placed RELA, ZEB1, IRX1, BARX1, HOXB7, ZNF362, AEBP1, MZF1, RUNX2 and ZBTB7B in the top 10. The top 20 included EMT (ZEB1), NF-*κ*B (RELA, NFKBIZ) and home-obox developmental factors (IRX1, BARX1, HOXB7, HOXB9). To assess single-seed artifacts, we recomputed IG on GNNs trained with three different seeds, each on its own cell-line split (Figure 3b). Two TFs (IRX1 and AEBP1) appeared in the top 20 across all three seeds. The top-20 overlap between seeds was 4 to 7 of 20, well above the 0.5 expected by chance, and Spearman correlations across all 771 TFs were 0.30 to 0.40, so the ranking replicated only partly. We therefore defined the core set as the ten TFs with the lowest mean IG rank across the training seeds: IRX1, ZEB1, KLF8, AEBP1, MZF1, SPDEF, STAT2, BARX1, SETBP1 and

RELA. Random Gaussian baselines gave a correlated ranking (Spearman *ρ* = 0.927, 10 of the top 20 shared), whereas a zero baseline yielded a distinct ranking (2 of the top 20 shared), led by E2F1, ZBTB4 and ESR1, reflecting a different attribution question (importance relative to zero activity rather than to the typical training cell).

The signed attributions (Figure 3c) were shared by the 11 drugs. Each of the top 20 TFs had the same sign in all 11 (220 of 220 TF and drug combinations), and the attribution vectors of the 11 drugs over the 771 TFs had a mean pairwise correlation of 0.97. The shared cell-line encoder does not force this by itself, because the prediction head combines the cell-line and drug embeddings, so the near-identical profiles suggest that the model adds a largely drug-independent cell-line component to the mean response of each drug. HOXB7, MZF1, ZBTB7B, AEBP1 and ZEB1 had the most negative attributions and BARX1, IRX1, RELA and E2F1 the most positive. Because IG attributions are relative to the baseline, their sign does not by itself give the direction in which higher activity of a TF moves the prediction. For 7 of the top 20 TFs the path-averaged gradient had the opposite sign in all 11 drugs (77 of the 220 combinations), because their mean activity in these TNBC lines lay on the other side of the training mean. Higher activity of ZEB1, RUNX2, ARID1A, HOXB9 and HIVEP1 raised the predicted AUC, and higher activity of ZNF362 and LHX1 lowered it. Supplementary Figure S4 shows, for comparison, per-drug Ridge regression coefficients of the ten core TFs for positive controls and candidate drugs. Unlike the GNN attributions, these models are fitted drug by drug.

### Predicted drug sensitivity is associated with overall survival

Per-drug Ridge transfer from PRISM, based on TF activities, yielded 1,447 drug models in each of three breast cancer cohorts that are not selected for TNBC (Table 2). Survival was modeled with penalized Cox models adjusted for the intrinsic subtype of the PAM50 classifier. In these models, the predicted AUC was associated with overall survival at a false discovery rate (FDR) below 0.05 for 623 of the 1,447 drugs in SCAN-B (7,215 patients, 1,164 events) and for 74 in METABRIC (1,979 patients, 1,143 events). The Cancer Genome Atlas breast cancer cohort (TCGA-BRCA) yielded no significant results after FDR correction, consistent with its limited statistical power (median follow-up 2.7 years against 7.0 years in SCAN-B, and 149 events in the 1,060 patients of the Cox models). QQ-plots of survival p-values showed marked inflation in SCAN-B (genomic-control inflation factor *λ*_GC_ = 7.98) and METABRIC (2.72), while TCGA-BRCA showed no inflation (0.84, Supplementary Figure S5). Per-patient predictions were correlated across drugs (mean inter-drug correlation r = 0.073 in SCAN-B, mean absolute correlation 0.16, with 34% of the drug pairs negatively correlated). These counts therefore measure how widespread the association is rather than the number of independent discoveries, and Benjamini-Hochberg control is not guaranteed for this family. The Benjamini-Yekutieli procedure, which is valid under any dependence, retained 391 drugs in SCAN-B and 15 in METABRIC.

**Table 2.**
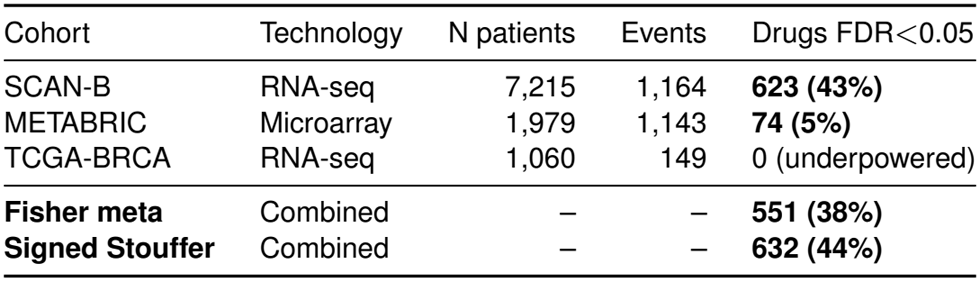
Survival associations of the predicted drug sensitivity in three breast cancer cohorts. Penalized Cox model per drug (penalizer *λ* = 0.1, PAM50-adjusted, two-sided Wald test), Benjamini-Hochberg FDR *<* 0.05 within each cohort over 1,447 drugs. Fisher meta-analysis of the per-cohort p-values with Benjamini-Hochberg correction over the 1,447 combined p-values, and a signed Stouffer combination that down-weights discordant directions (Methods).

At the individual-drug level, stratifying SCAN-B patients at the median predicted AUC produced a significant survival separation whose direction differed between drugs (Figure 4a). For the cytotoxic chemotherapy positive controls (paclitaxel, epirubicin), patients predicted sensitive (below-median predicted AUC) had significantly worse overall survival than predicted-resistant patients. This pattern is consistent with a proliferation paradox, in which tumors predicted sensitive to antimitotic and DNA-damaging agents tend to be more proliferative and therefore carry a worse baseline prognosis in this mixed-treatment cohort: the half predicted sensitive to paclitaxel contained 15% basal-like tumors, against 4% in the other half. For erlotinib and entinostat, two of the candidates selected later with a hazard ratio (HR) above 1, the direction reversed, with predicted-sensitive patients surviving longer. The separations persisted, although weaker, when the comparison was stratified by PAM50 subtype (p = 7.0 *×*10^*−*3^, 1.9 *×*10^*−*4^, 2.1*×* 10^*−*2^ and 4.9 *×*10^*−*7^). This difference between drugs was graded at the quintile level, where paclitaxel (a cytotoxic control) and entinostat (a candidate) showed opposite trends (log-rank test for trend over the quintile index, p = 8.1 *×*10^*−*8^ and 4.9*×* 10^*−*12^, Figure 4b).

**Figure 4.**
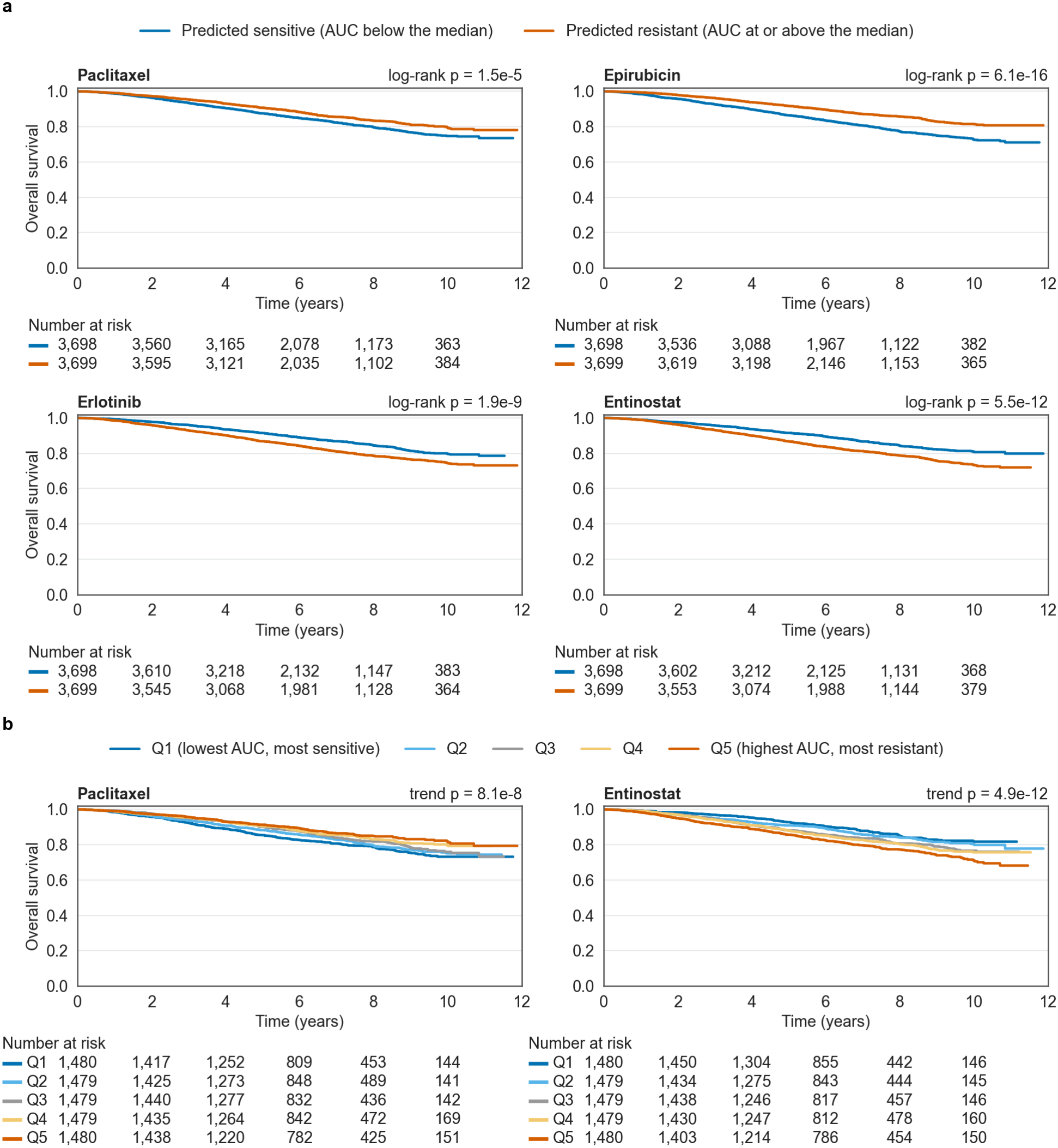
SCAN-B survival stratified by predicted drug sensitivity. (**a**) Kaplan-Meier curves for patients below (predicted sensitive, blue) and at or above (predicted resistant, vermillion) the median predicted AUC, for two cytotoxic positive controls (paclitaxel, epirubicin) and two candidates (erlotinib, an epidermal growth factor receptor (EGFR) inhibitor, and entinostat, a histone deacetylase (HDAC) inhibitor). The direction of separation differs between these drugs: for the cytotoxic controls, predicted-sensitive patients show significantly worse survival, whereas for the targeted candidates predicted-sensitive patients survive longer (the direction imposed by the HR *>* 1 filter of the selection rule). (**b**) Quintiles of predicted AUC for paclitaxel (cytotoxic control) and entinostat (candidate): paclitaxel shows a graded association (most-sensitive Q1, dark blue, worst), while entinostat shows the opposite direction (most-sensitive Q1 best). The titles give the p-value of the log-rank test for trend over the quintile index. The curves use all 7,397 SCAN-B patients with overall survival, including those without a PAM50 subtype (Methods). The log-rank tests are two-sided and not adjusted for PAM50 subtype, which differs between the groups, and the absolute differences in overall survival at ten years are 4.8 to 8.8 percentage points. Numbers at risk are given below each panel at two-year intervals. Time axis in years.

The direction of the associations largely agreed across platforms. Of the 54 drugs significant in both SCAN-B (RNA-seq) and METABRIC (microarray), 53 (98%) had hazard ratios in the same direction (Figure 5), a concordance partly expected from conditioning on significance in both cohorts and from the shared prognostic axis described below. Across all 1,447 drugs the direction agreed for 70%, against 52% expected under independence, and the p-values of the two cohorts correlated weakly (Spearman *ρ* = 0.20, Supplementary Figure S6). Fisher meta-analysis combining p-values across three cohorts identified 551 drugs (38% of 1,447 tested) at FDR *<* 0.05 (Figure 6), but it ignores the direction of the hazard ratios, and 75 of the 551 had opposite directions in SCAN-B and METABRIC. A signed Stouffer combination identified 632 drugs, 502 of them also Fisher-significant and 604 with concordant SCAN-B and METABRIC hazard ratios.

**Figure 5.**
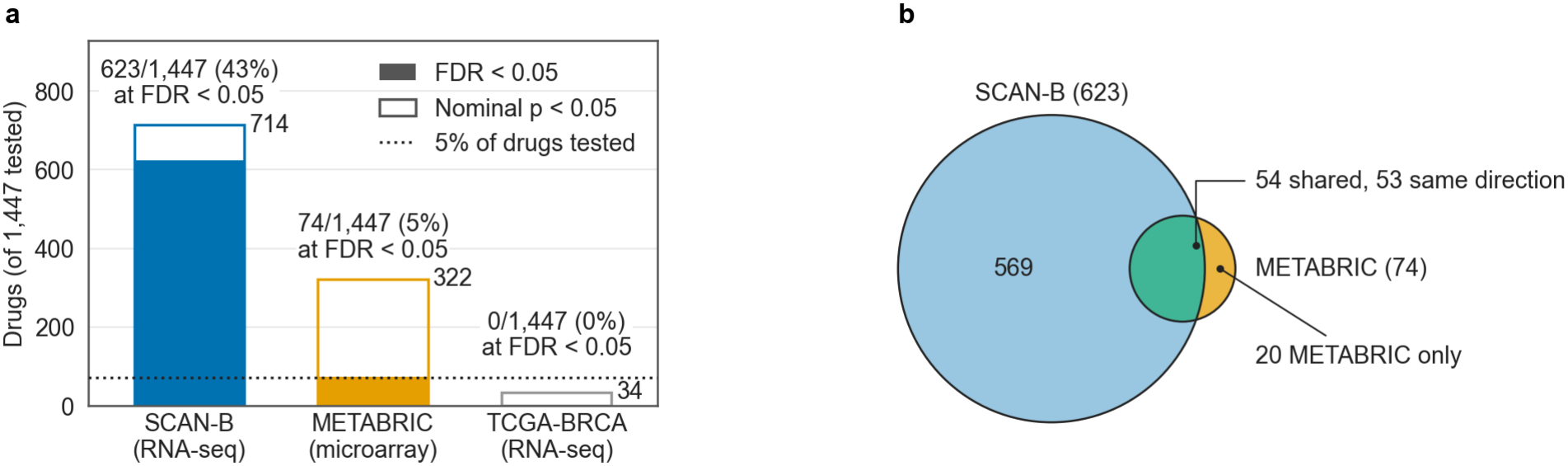
Multi-cohort survival associations of TF-based drug sensitivity predictions. (**a**) Number of drugs with a significant survival association (FDR *<* 0.05, filled bars) in each cohort: 623 in SCAN-B (RNA-seq), 74 in METABRIC (microarray) and none in the underpowered TCGA-BRCA. Outline bars give the drugs at nominal p *<* 0.05 (714, 322 and 34), and the dotted line marks 5% of the 1,447 drugs, the number expected at that threshold under a calibrated null. Because the survival p-values are inflated (Supplementary Figure S5), the line is a reference and not a test of excess. (**b**) Overlap between SCAN-B and METABRIC: 54 drugs are significant in both cohorts, of which 53 (98%) show a consistent hazard-ratio direction.

**Figure 6.**
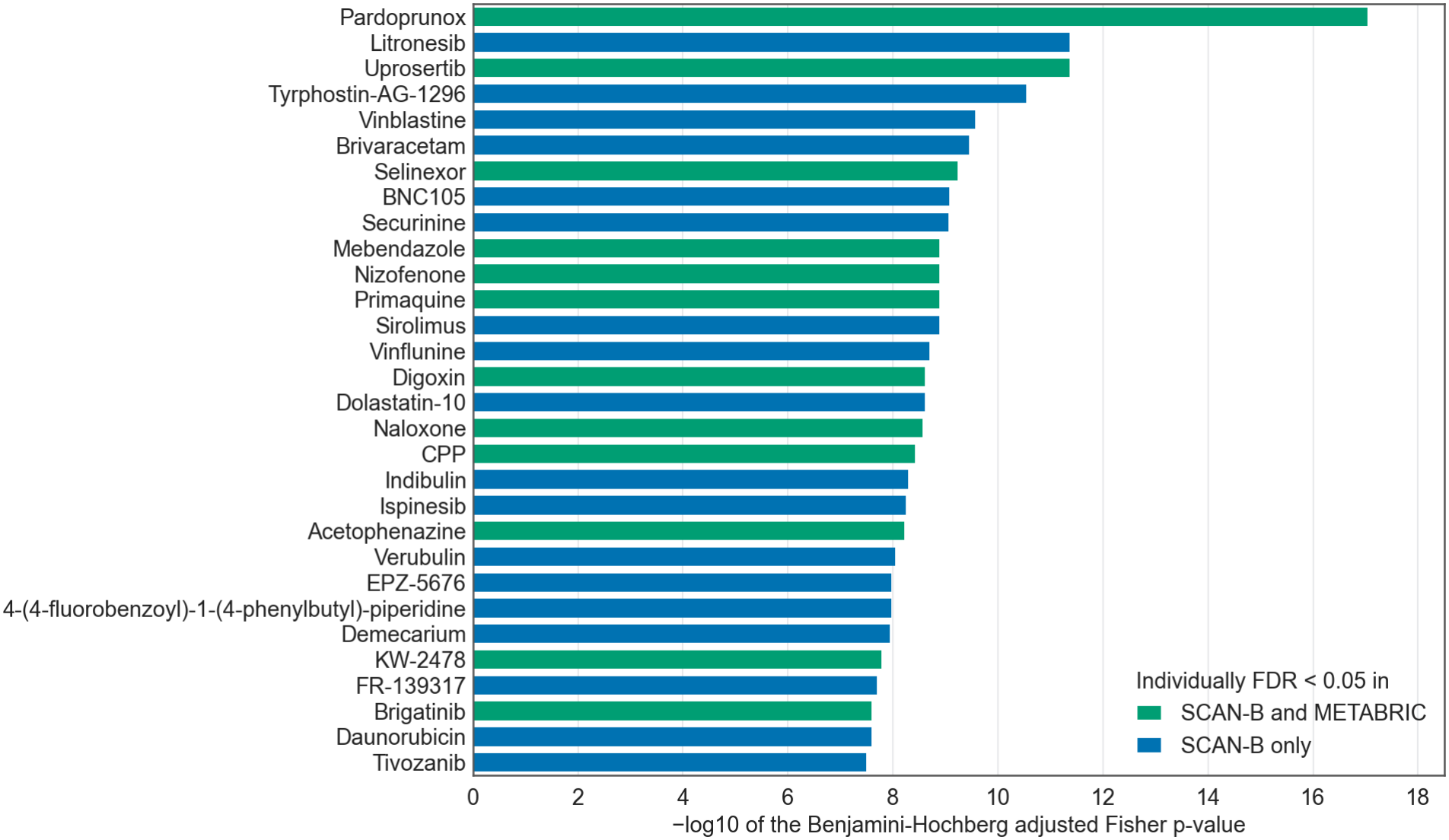
The 30 most significant drugs in the Fisher meta-analysis across three cohorts. Bar length is *−* log_10_ of the Benjamini-Hochberg adjusted Fisher p-value. Bar color indicates the cohort(s) in which the drug is individually FDR *<* 0.05: SCAN-B only or both SCAN-B and METABRIC (none of the 30 is significant in METABRIC alone). These are the strongest survival associations regardless of direction or mechanism. The repurposing candidates are selected separately by the rule described in Methods.

Five clinically established breast cancer drugs served as positive controls: the taxanes paclitaxel and docetaxel, the anthracycline epirubicin and the PARP inhibitors olaparib and talazoparib (Table 3, Supplementary Figure S7). The platinum salts used in TNBC were not among them, and carboplatin and cisplatin had no survival association in SCAN-B (*p*_adj_ = 0.86 and 0.65). All five controls were individually FDR-significant in SCAN-B, all with HR *<* 1 (higher predicted resistance associated with better survival), and four reached significance in the Fisher meta-analysis: epirubicin, talazoparib, paclitaxel and docetaxel. Only epirubicin was also significant in METABRIC, and paclitaxel even shifted above HR = 1 there, so the other three reached Fisher significance through SCAN-B. With 38% of all drugs significant in the Fisher meta-analysis, four of five controls is only marginally more than expected by chance (binomial p = 0.073), and the controls ranked between 82nd and 682nd of the 1,447 drugs. The direction of their associations, HR *<* 1, shows that predicted sensitivity to these drugs tracks a poor-prognosis phenotype, not that the transfer isolates drug-specific signal. Olaparib and talazoparib were approved on trials in carriers of germline BRCA1/2 mutations (Robson et al., 2017; Litton et al., 2018), a status not available in these cohorts, so their associations in unselected patients, like those of the cytotoxic drugs, are more plausibly driven by proliferation than by homologous-recombination deficiency.

**Table 3.** Positive control drugs across cohorts. Hazard ratios per unit of predicted AUC from the PAM50-adjusted penalized Cox models, with approximate 95% confidence intervals of the penalized fit in brackets and Benjamini-Hochberg *p*_adj_ over the 1,447 drugs in parentheses. Bold marks Fisher or Stouffer *p*_adj_ *<* 0.05.

| Drug | SCAN-B HR [95% CI] (*p*<sub>adj</sub>) | METABRIC HR [95% CI] (*p*<sub>adj</sub>) | Fisher *p*<sub>adj</sub> | Stouffer *p*<sub>adj</sub> |
|---|---|---|---|---|
| Paclitaxel | 0.47 [0.32–0.70] (1.1e-3) | 1.25 [0.77–2.03] (0.61) | **8.6e-3** | 7.0e-2 |
| Docetaxel | 0.50 [0.34–0.74] (2.4e-3) | 0.98 [0.64–1.50] (0.97) | **3.1e-2** | **2.3e-2** |
| Epirubicin | 0.15 [0.07–0.34] (5.7e-5) | 0.12 [0.04–0.42] (2.5e-2) | **3.2e-6** | **2.6e-7** |
| Talazoparib | 0.40 [0.29–0.57] (4.9e-6) | 0.93 [0.57–1.51] (0.89) | **1.4e-4** | **5.1e-4** |
| Olaparib | 0.60 [0.41–0.87] (1.9e-2) | 0.85 [0.55–1.33] (0.70) | 1.2e-1 | 5.7e-2 |

The METABRIC attenuation is not explained by the number of deaths (1,143, against 1,164 in SCAN-B). It more likely reflects the microarray platform and its restricted gene set, an older treatment era and competing mortality, since only 646 of these deaths were from breast cancer (43% from other causes). With disease-specific survival as the outcome, 184 drugs reached FDR *<* 0.05 in METABRIC instead of 74, only 30 of them significant with both endpoints, and paclitaxel’s hazard ratio fell below 1 (0.82, non-significant). Mechanism-of-action (MOA) and target enrichment of the Fisher-significant drugs identified a single enriched class, tubulin polymerization inhibitors (25 of 27 significant, FDR = 3.2 *×*10^*−*7^), and its tubulin targets (Supplementary Figures S8 and S9). Because drugs within a class have correlated predictions, this reflects the proliferation-linked axis of antimitotic agents rather than independent convergence of many drugs.

Principal component analysis of the per-drug prediction matrix of SCAN-B identified a shared prognostic axis common to the TF-based predictions: the first component (PC1) explained 12% of the prediction variance (the second 11%) and was itself associated with over-all survival (PAM50-adjusted penalized Cox, p = 1.6*×* 10^*−*9^). Oriented so that higher scores mean higher predicted AUC for paclitaxel, PC1 correlated negatively with the activity of E2F1 (r = *−*0.55) and MYC (r = *−*0.37) and positively with that of TP53 (r = 0.45) and ESR1 (r = 0.37). This pattern supports its reading as a proliferation-linked prognostic axis. Residualizing PC1 reduced the number of drugs with a nominal survival association from 90 to 54 among the 200 most variable drugs, so the SCAN-B hits reflect a mixture of this shared prognostic axis and residual signal that may come from other shared components or be drug-specific. Most associations also ran in the direction of this axis: 464 of the 623 SCAN-B associations had HR *<* 1, the direction of the cytotoxic controls. As a check of the clinical TF activities, univariate Cox regression in SCAN-B showed that high MYC activity (HR = 1.13 per unit of activity, p = 3.0 *×*10^*−*5^) and high E2F1 activity (HR = 1.12, p = 4.1 *×*10^*−*5^) both associated with worse overall survival, whereas high TP53 activity associated with better survival (HR = 0.90, p = 1.5 *×*10^*−*4^). These three TFs were not among the most significant (ranks 262, 267 and 295 of 769), which included luminal regulators such as TOX3 and GATA3.

SCAN-B records whether each patient received chemotherapy (2,802 yes, 4,041 no and 372 unknown among the 7,215 patients) and endocrine therapy, but not the agents. If predicted sensitivity to a cytotoxic drug identified patients who benefit from it, its association should differ between treated and untreated patients. For the taxanes it did not. In Cox models with PAM50 as strata, the interaction between the standardized predicted AUC and chemotherapy, expressed as the ratio of the hazard ratios in treated and untreated patients, was not significant for paclitaxel (1.02, p = 0.80) or docetaxel (1.05, p = 0.45), and their association was present in patients who did not receive chemotherapy (Supplementary Table S1). For epirubicin (1.37, p = 5.8*×* 10^*−*6^) and talazoparib (1.30, p = 1.6 *×*10^*−*4^), the adverse association of predicted sensitivity seen in untreated patients (HR = 0.76 and 0.74 per standard deviation) was absent in treated patients. Talazoparib is not a chemotherapy agent, so for it this pattern could at most reflect how chemotherapy changes the prognosis of the tumors predicted sensitive. Neither inter-action survived adjustment for the clinical factors that guide the use of chemotherapy: with age, tumor stage, grade, nodal status, ER and PR status and endocrine therapy as main effects (6,173 patients, 946 events), the interaction was not significant for epirubicin (p = 0.18) or talazoparib (p = 0.50), and none of the 12 interactions remained significant after correction. In METABRIC, where 412 of the 1,979 patients received chemotherapy, no interaction reached p *<* 0.05.

### Core TFs in matched primary and metastatic tumors

To ask whether the core TFs vary in patient tumors, we analyzed the AURORA-US cohort of 129 samples from 53 patients, restricted to the 39 patients with both primary and matched metastatic samples (Garcia-Recio et al., 2023). Five of the ten TFs (KLF8, STAT2, AEBP1, SPDEF and SETBP1) survived Bonferroni correction for multiple testing (ten tests, p *<* 0.005, Table 4, Figure 7a). At the same threshold, 231 of the 770 TFs (30%) changed (Figure 7b), so five of ten was close to the three expected for a random set of ten TFs (hypergeometric p = 0.15).

**Table 4.** Core TF activity in primary vs. metastatic breast tumors (AURORA-US, 39 paired patients). Primary and Metastasis give the mean TF activity (ULM score) across patients, with metastatic samples averaged per patient. Δ is metastasis minus primary, computed before rounding. p-values are from two-sided paired Wilcoxon signed-rank tests, unadjusted. In the Bonferroni column, sig marks p *<* 0.005, the Bonferroni threshold over the ten core TFs, and n.s. marks TFs above it.

| TF | Primary | Metastasis | Δ | p-value | Bonferroni |
|---|---|---|---|---|---|
| KLF8 | +3.48 | +3.22 | −0.26 | 3.8e-5 | **sig** |
| STAT2 | +6.91 | +6.38 | −0.54 | 3.9e-4 | **sig** |
| AEBP1 | +1.87 | +2.03 | +0.16 | 8.4e-4 | **sig** |
| SPDEF | +1.77 | +2.11 | +0.34 | 2.2e-3 | **sig** |
| SETBP1 | +0.21 | −0.09 | −0.31 | 2.4e-3 | **sig** |
| ZEB1 | −3.39 | −3.56 | −0.17 | 0.060 | n.s. |
| MZF1 | +3.15 | +3.23 | +0.08 | 0.10 | n.s. |
| RELA | +19.02 | +18.47 | −0.55 | 0.15 | n.s. |
| IRX1 | +0.52 | +0.47 | −0.05 | 0.34 | n.s. |
| BARX1 | +3.66 | +3.63 | −0.03 | 0.81 | n.s. |

**Figure 7.**
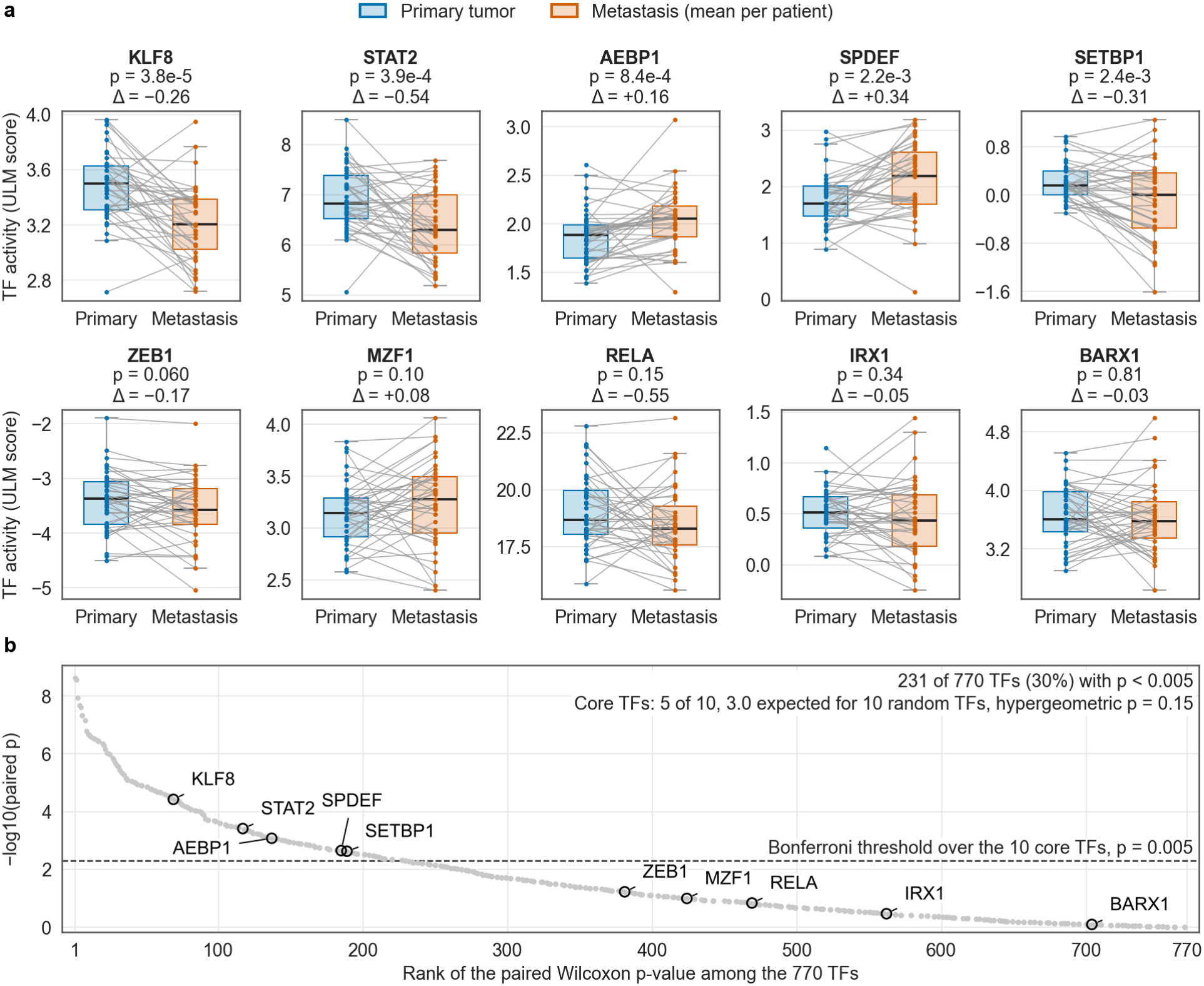
AURORA-US: core TF activity in primary vs. matched metastatic breast tumors, against the background of all TFs. (**a**) Each panel shows one of the ten core TFs, the same ten reported in Table 4, for the 39 patients with both primary and metastatic samples: one line per patient joins the primary value to the mean of that patient’s metastatic samples. Titles give the paired Wilcoxon p-value (unadjusted) and the mean paired difference Δ, metastasis minus primary. Boxes summarize the per-patient values (median line, whiskers to 1.5 times the interquartile range). Blue for primary, vermillion for metastasis. (**b**) The same paired test applied to all 770 TFs of the AURORA-US activity matrix, ranked by p-value (gray points), with the ten core TFs circled and labeled. The dashed line marks p = 0.005, the Bonferroni threshold over the ten core TFs used in Table 4. Five of the ten core TFs pass it, as do 231 of the 770 TFs (30%).

Bulk TF activity also reflects the stromal and immune content of each biopsy, and metastases differ from primaries in host organ, tumor purity, prior treatment and, for 66 of the 129 samples, collection at autopsy, so these differences may reflect tissue composition as much as tumor-intrinsic reprogramming. Restricting the primary-versus-metastasis comparison to paired patients grouped by biopsy tissue site (liver, brain, lymph node, lung, chest, skin and soft tissue) showed that KLF8 was direction-concordant in all seven sites, whereas STAT2, AEBP1 and SETBP1 retained the pooled direction in five of the seven sites and SPDEF in four. Among the 231 TFs that passed the same threshold, 47% kept the direction in all seven sites and 96% in at least five, so the site pattern of the core TFs was not unusual. Drug sensitivity predictions for AURORA-US samples, paired by patient as in the TF analysis, revealed that 711 of the 1,447 drugs showed significantly different predicted sensitivity between primary and metastatic tumors (FDR *<* 0.05, Supplementary Figure S10). Of these, 597 were predicted more active (lower AUC) in metastases and 114 less active. Entinostat showed increased predicted resistance in metastatic tumors (paired p = 6.2*×* 10^*−*3^), while paclitaxel and panobinostat did not show significant changes (p = 0.33 and p = 0.88, respectively).

### Rule-based repurposing candidates

Applying the selection rule described in Methods to the SCAN-B PAM50-adjusted penalized Cox results yielded seven candidates (Table 5). The statistical filter retained 143 drugs with *p*_adj_ *<* 0.05 and a hazard ratio per unit of predicted AUC in [1.5, 10] at the canonical Cox penalizer (*λ* = 0.1). Of these, 52 remained significant at *λ* = 0.5, 20 of the 52 were annotated as kinase or HDAC inhibitors and 16 of the 20 had reached at least phase 2. Seven of the 16 fell among the 30 most significant of the 143 drugs (Supplementary Table S2), at ranks 14 to 26, and 85 to 161 among all 1,447 drugs. The selection rule depends only on objective metrics and involves no manual curation. Its thresholds, however, were fixed after preliminary exploration of the data rather than pre-specified. To assess whether this choice shaped the results, we repeated the analysis with a looser rank cutoff and without the mechanism-class filter. Relaxing the rank cutoff to 50 added seven kinase inhibitors: BMS-690514, BVD-523 (ulixertinib), indirubin, dasa-tinib, lestaurtinib, rociletinib and AZD8931. Repeating the Integrated Gradients analysis on all 16 kinase and HDAC inhibitors that pass the development-stage filter, together with the four positive controls other than tala-zoparib (20 drugs instead of 11), left the attribution results essentially unchanged: 19 of the 20 most important TFs were the same, all ten core TFs were shared, and the same five core TFs were Bonferroni-significant in AURORA-US (Supplementary Note 1). Without the mechanism-class filter, the most significant drugs were dominated by central nervous system and other non-oncology agents (Supplementary Data 1).

**Table 5.** Repurposing candidates selected by the rule of Methods (statistical, stability, mechanism-class and development-stage filters, rank cutoff of 30) from the SCAN-B PAM50-adjusted penalized Cox analysis. HR_0.1_ is the hazard ratio per unit of predicted AUC at the canonical penalizer *λ* = 0.1, with the approximate 95% confidence interval of the penalized fit in brackets. HR/SD is the same estimate per standard deviation of the predicted AUC across patients. The *λ* = 0.5 columns report the stability filter, which every candidate passes by construction. *r*_test_ is the Pearson correlation of the transfer model on the held-out cell lines with a measured response to the drug (83 to 95 of the 95). Mechanism of action (MOA) and phase as annotated in PRISM (release 19Q4), curated by the Broad Drug Repurposing Hub (Corsello et al., 2017). inh., inhibitor.

| Drug | MOA | Phase | HR<sub>0.1</sub> [95% CI] | *p*<sub>adj,0.1</sub> | HR/SD | HR<sub>0.5</sub> | *p*<sub>adj,0.5</sub> | *r*<sub>test</sub> |
|---|---|---|---|---|---|---|---|---|
| Osimertinib | EGFR inh. | Launched | 6.38 [3.02–13.48] | 1.3e-5 | 1.12 | 2.09 | 1.5e-2 | 0.29 |
| Saracatinib | Src inh. | Phase 2/3 | 5.50 [2.74–11.02] | 1.7e-5 | 1.12 | 2.01 | 1.2e-2 | 0.36 |
| Erlotinib | EGFR inh. | Launched | 4.28 [2.51–7.30] | 1.5e-6 | 1.14 | 1.94 | 1.7e-3 | 0.06 |
| Brigatinib | ALK / EGFR inh. | Launched | 3.63 [2.24–5.88] | 2.6e-6 | 1.13 | 1.74 | 3.4e-3 | 0.10 |
| Pelitinib | EGFR inh. | Phase 2 | 3.42 [2.03–5.77] | 4.0e-5 | 1.11 | 1.56 | 3.9e-2 | 0.47 |
| Entinostat | HDAC inh. | Phase 3 | 3.28 [1.94–5.54] | 8.4e-5 | 1.11 | 1.71 | 1.3e-2 | 0.04 |
| Trametinib | MEK inh. | Launched | 2.00 [1.48–2.69] | 4.7e-5 | 1.11 | 1.34 | 1.4e-2 | 0.50 |

The HR *>* 1 filter is a heuristic. For the cytotoxic positive controls, the tumors predicted most sensitive have the worst survival, an HR *<* 1 that reflects the proliferation paradox (Figure 4a). Requiring HR *>* 1 selects the opposite pattern, drugs for which predicted sensitivity is instead associated with better survival. This association can arise from a less proliferative, more luminal phenotype of the tumors predicted to be sensitive. For five of the seven candidates basal-like tumors were predicted less sensitive than luminal A tumors, and for all candidates except pelitinib luminal A tumors were over-represented in the half of the cohort predicted sensitive (up to 67%, against 49.5% of the whole cohort). Hazard ratios are expressed per unit of predicted AUC, a range much wider than the spread of predictions across patients, and on the standard-deviation scale the candidates have hazard ratios of only 1.11 to 1.14 (Table 5).

In the penalizer sweep (Supplementary Figure S11) the seven remained significant at *λ* = 0.01, but only erlotinib did at *λ* = 1.0. The HER2-enriched and luminal A terms of the SCAN-B Cox models violated proportional hazards in all 12 models of the candidates and controls, and so did the prediction terms of six of them (Supplementary Table S3). With PAM50 as a stratification factor instead of covariates, no prediction term violated the assumption in the penalized or the unpenalized fit (smallest p = 0.11). In the unpenalized stratified model all seven candidates remained significant, with hazard ratios of 1.14 to 1.26 per standard deviation. In METABRIC all seven had HR *>* 1 for overall survival, and brigatinib and pelitinib reached FDR significance. With disease-specific survival five had HR *>* 1, trametinib (0.85) and entinostat (0.88) fell below 1, and only brigatinib remained significant (*p*_adj_ = 0.034, Supplementary Table S1).

We then adjusted for age, tumor stage, grade, nodal status, ER and PR status and the receipt of chemotherapy and endocrine therapy, with PAM50 as strata (6,231 SCAN-B patients, 959 events). The hazard ratios of the candidates shrank to 1.01 to 1.17 per standard deviation, and with Benjamini-Hochberg correction over the 12 drugs only brigatinib and saracatinib remained significant, together with the control talazoparib (Supplementary Table S3). In METABRIC, brigatinib and pelitinib survived the analogous adjustment (Supplementary Table S1). Except for osimertinib and pelitinib, the candidates’ associations were stronger in patients who did not receive chemotherapy, most clearly for entinostat (HR = 1.24 per standard deviation without and 0.93 with chemotherapy, interaction p = 9.5*×* 10^*−*6^). As for the controls, none of these interactions remained significant after adjustment for the clinical covariates and endocrine therapy, so they are more consistent with differences in the prognosis of treated and untreated patients than with any effect of the candidate drugs.

In the 676 SCAN-B patients with ER-, PR- and HER2-negative tumors (183 events), all seven candidates had HR *>* 1 per standard deviation, but only pelitinib (HR = 1.43, *p*_adj_ = 1.9 *×*10^*−*6^) and osimertinib (HR = 1.24, *p*_adj_ = 0.011) remained significant after correction for the seven tests, and none of the five positive controls was associated with survival. Among the 686 PAM50 basal-like patients, pelitinib and trametinib were significant (Supplementary Table S3). This subgroup, like the adjusted and treatment analyses above, reuses the SCAN-B patients in which the candidates were selected, so these analyses test the stability of the associations rather than replicate them.

Four of the seven candidates (osimertinib, erlotinib, brigatinib and pelitinib) are annotated as EGFR inhibitors, brigatinib being primarily an anaplastic lymphoma kinase (ALK) inhibitor, and saracatinib (Src) and trametinib (MEK1/2) act on kinases shared by many receptor pathways. This grouping does not indicate convergence on EGFR: the mechanism-class filter admits only kinase and HDAC inhibitors, and EGFR was not among the targets enriched among the drugs with a significant Fisher association. Entinostat completes the candidate list as the lone HDAC inhibitor and has already been tested in breast cancer. In advanced TNBC, adding it to atezolizumab did not pro-long progression-free survival in the randomized phase 2 ENCORE 602 trial (O’Shaughnessy et al., 2020), and in hormone receptor-positive advanced breast cancer adding it to exemestane improved neither progression-free nor overall survival in the phase 3 E2112 trial (Connolly et al., 2021).

### Sensitivity analyses

The number of significant drugs and the range of hazard ratios depended on regularization (Supplementary Figure S11). At the canonical *λ* = 0.1, 623 drugs reached FDR *<* 0.05, with hazard ratios from 0.009 to 54.4. Increasing the penalizer to *λ* = 0.5 compressed the HR range among significant drugs to [0.13, 5.53] while retaining 306 drugs at FDR *<* 0.05, and at *λ* = 1.0 only 87 remained. Decreasing it to a near-unpenalized regime (*λ* = 0.01) gave 678 significant drugs and inflated the hazard ratios up to 245.5. Expressed per standard deviation of the predicted AUC, the significant hazard ratios at *λ* = 0.1 spanned only 0.81 to 1.17. Replacing over-all survival with relapse-free survival as the outcome in the same PAM50-adjusted penalized Cox model yielded direction-concordant hazard ratios for all 12 candidate and positive-control drugs. Eight remained significant at FDR *<* 0.05 over these 12 tests (erlotinib, brigatinib, entinostat, osimertinib and saracatinib among the candidates, and docetaxel, epirubicin and talazoparib among the positive controls). We also swapped the QuantileTransformer, which maps each TF activity to a normal distribution fitted on the training cell lines, for ComBat empirical-Bayes adjustment on the joint PRISM and SCAN-B TF-activity matrix. This swap yielded 634 drugs at FDR *<* 0.05 (against 623 with the Quantile-Transformer), but only 367 were significant under both preprocessing strategies, with 98.6% direction concordance among them. All seven candidates remained significant under ComBat, whereas paclitaxel, docetaxel and talazoparib did not, so the individual drug lists depended on the normalization.

Graph component ablation with five seeds per configuration (Supplementary Figure S12) revealed that removing the PPI or the CollecTRI edges alone had minimal impact on global prediction performance (0.742 in both cases, against 0.743 for the full graph). Removing the drug-target edges, the only relation that passes messages to the drugs, lowered it to 0.726, the level of the other configurations without drug-target edges (0.722 with no prior-knowledge edges and 0.731 with the CollecTRI edges only). Rewiring the drug-target edges at random, with each drug keeping its number of targets and each gene the number of drugs that target it, kept 86% of the gain (0.741 *±*0.001), so most of the gain does not depend on which genes are annotated as targets. The gain was confined to global correlation: the median per-drug correlation was not higher with prior-knowledge edges (0.032 with the full graph against 0.037 to 0.052 for the reduced graphs).

Within-subtype analysis in SCAN-B showed that the survival associations of predicted drug sensitivity were also present within the PAM50 basal-like subtype and the pooled basal-like and HER2-enriched subtypes. Basal-like tumors alone (n = 686, 167 events) yielded 117 drugs at FDR *<* 0.05, and pooling basal-like and HER2-enriched tumors (n = 1,361, 314 events) yielded 221 (Supplementary Figure S13), consistent with the larger number of events. These within-subtype screens used a median-split log-rank test, so their counts are not directly comparable with those of Table 2.

### The candidates in TNBC cell lines and in a triple-negative cohort

Two analyses asked whether the candidates relate to TNBC itself rather than to unselected breast cancer. In PRISM we compared a composite of the seven candidates between the 12 TNBC cell lines and the remaining 464. At 80% power the design detects a Cliff’s delta of 0.473, which only 6 of the 1,409 drugs reach, so it can exclude only a large difference. Twelve lines can carry a large effect of this kind: random draws of twelve skin lines recovered the BRAF inhibitor dabrafenib at a delta of *−*0.71 (Figure 8a). The composite gave a Cliff’s delta of 0.095 (95% confidence interval *−*0.236 to 0.426, two-sided permutation p = 0.6, Figure 8a,b), none of the 1,409 evaluable drugs reached FDR *<* 0.05, and the candidates sat at a mean percentile of 52.1 among them. Against matched random sets of kinase or HDAC inhibitors, the candidates did not stand out (two-sided p = 0.6). Of the 17 checks of established effects with an expected direction, only two met their criterion, both for paclitaxel: against the other breast lines in PRISM and against all other lines in GDSC, a contrast that the breast lineage dominates.

**Figure 8.**
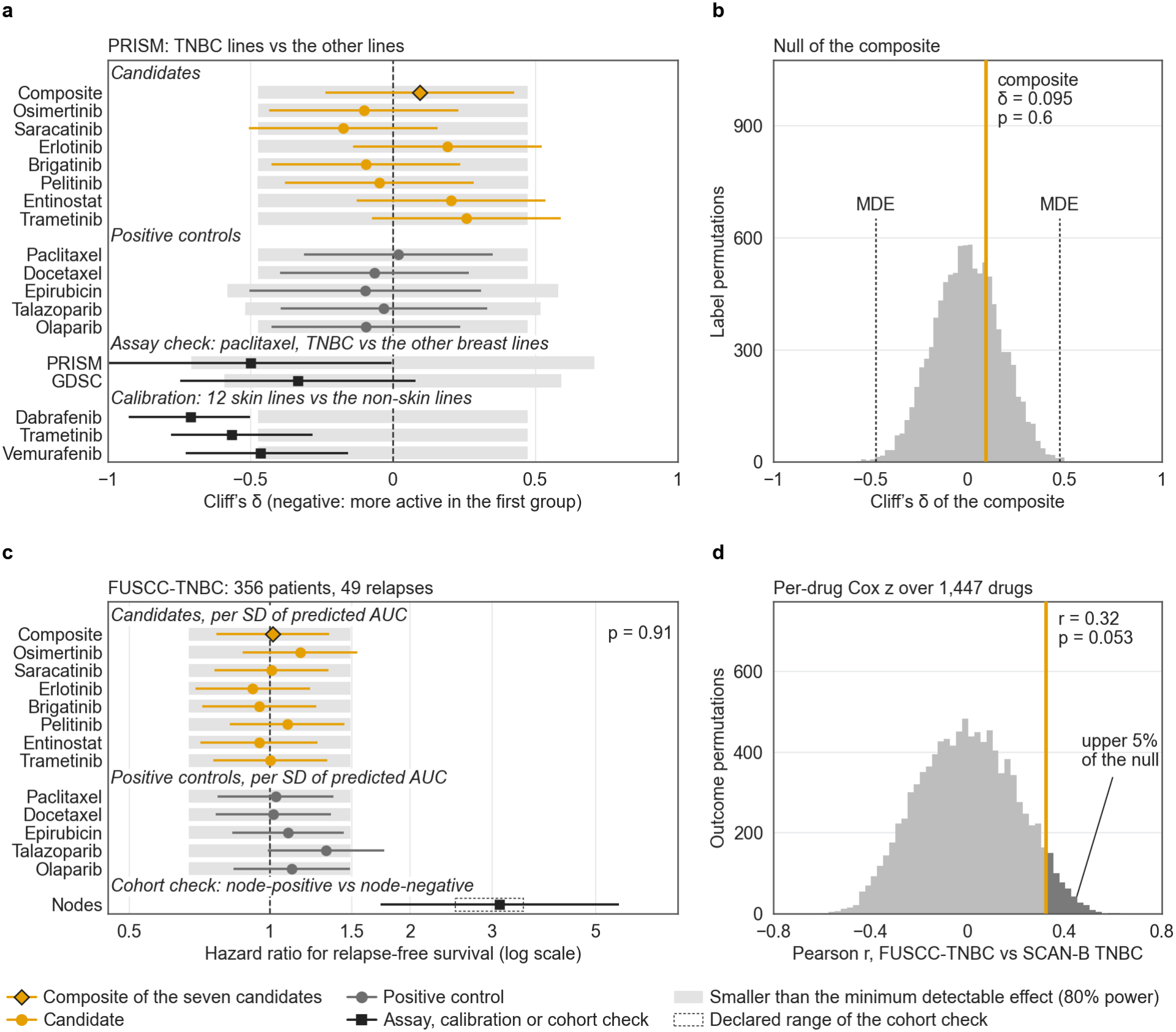
The candidates in TNBC cell lines and in a triple-negative cohort. (**a**) Cliff’s *δ* between the TNBC lines of PRISM and the other lines for the composite of the seven candidates (12 and 464 lines), each candidate and each positive control (the lines measured for each drug), with 95% confidence intervals from the null standard error. Gray bars span the effects smaller than the minimum detectable effect (MDE) of each comparison at 80% power, 0.473 for the composite. The assay-check rows compare paclitaxel between the TNBC lines and the other breast lines in PRISM and GDSC, and the calibration rows compare 12 randomly drawn skin lines with the non-skin lines (median and central 95% of 200 draws). Of the 17 assay checks with an expected direction (Methods), 2 met their criterion, the PRISM row shown here and paclitaxel against all other lines in GDSC, and the lowest calibration detection rate, 0.79, fell short of the required 0.8. (**b**) The composite (orange line) against its null over 10,000 permutations of the cell-line labels (two-sided p = 0.6). Dotted lines mark the MDE. Of the 1,409 evaluable drugs, 6 reach it and none reaches FDR *<* 0.05. (**c**) Hazard ratios for relapse-free survival in FUSCC-TNBC (356 patients, 49 relapses) per standard deviation of the predicted AUC or of the composite, from univariate, unpenalized Cox models, with 95% confidence intervals. The composite had a hazard ratio of 1.02 (permutation p = 0.91). Gray bars span hazard ratios from 1/1.49 to 1.49, the range below the MDE at 80% power. The cohort check, the hazard ratio of node-positive against node-negative disease (3.11), had to fall in the declared range, 2.5 to 3.5 (dashed box). (**d**) Pearson correlation over 1,447 drugs between the per-drug Cox score statistics of FUSCC-TNBC and those of the triple-negative subgroup of SCAN-B (676 patients, 183 deaths, overall survival), against its null over 10,000 permutations of the FUSCC-TNBC outcomes. The observed r = 0.32 lies just below the 95th percentile of the null, 0.33 (darker bars, one-sided p = 0.053). No drug reached FDR *<* 0.05 in FUSCC-TNBC.

The second analysis used FUSCC-TNBC (Jiang et al., 2019), 356 patients and 49 relapses from a single center in Shanghai. Its endpoint was relapse-free survival, whereas the reference, the triple-negative subgroup of SCAN-B, was analyzed for overall survival. With 49 events the design detects only hazard ratios above 1.49 per standard deviation (Figure 8c), so it cannot tell the candidates apart from the rest of the screen. The cohort is not inert at that scale: positive lymph nodes carried a hazard ratio of 3.11 in it (3.08 in the complete cases). The sequenced patients did not differ in relapse risk from the rest of the cohort (hazard ratio 0.92, p = 0.76). Across the 1,447 drugs, the per-drug Cox score statistics of FUSCC-TNBC correlated with those of the reference at Pearson r = 0.32, just below the 95th percentile of its permutation null (0.33, one-sided permutation p = 0.053, Figure 8d). The composite of the seven candidates was flat, at a hazard ratio of 1.02 per standard deviation (permutation p = 0.91, 0.99 to 1.09 in the five sensitivity analyses of Methods), and no drug reached FDR *<* 0.05. Neither the composite nor the node term departed from proportional hazards (p = 0.099 and 0.084). Of the 136 terms tested (Methods), 22 did, among them tumor size in the 13 clinically adjusted models and in its univariate fit (p = 0.035) and the prediction terms of brigatinib and trametinib in their univariate fits (p = 0.037 and 0.045).

## Discussion

We present PRECISION, a framework combining a heterogeneous GNN on a prior-knowledge graph with per-drug clinical transfer of TF-based predictions for drug repurposing in breast cancer. Three findings shape how the results should be read.

First, in the cell-line hold-out the GNN did not outper-form the Random Forest and had the lowest per-drug correlation of the models compared (Figure 2), and its global correlation fell below that of a reference that ignores the cell line. Because cell lines receive no messages in the graph, the prior knowledge encoded in the graph edges reaches the prediction only through the drug representations, whereas CollecTRI also enters the cell-line features through the TF activities. The ablation showed a small gain in global correlation when the drug-target relation was kept, most of which remained when its edges were rewired at random, and no gain in per-drug correlation.

Second, Integrated Gradients on the trained model gave TF-level attributions that were partly reproducible across training seeds and shared by the drugs examined. The attributions are taken with respect to the cell-line TF inputs, which receive no messages from the graph. Whether prior knowledge changes which TFs are identified was therefore not tested. In the pairs where the attribution of the full cell-line feature vector was measured, the TF inputs carried about a quarter of it, and the rest fell on the expression features. The value of the model in this study is that it provides one trained predictor on which attribution can be computed, not pre-dictive accuracy or graph-derived regulatory structure. What connects the two stages of the pipeline is the shared TF-activity representation: the TF activities that make up most of the cell-line features of the graph (771 of 1,271) are also the features on which the per-drug Ridge models carry predictions to patients.

Third, TF-based predicted sensitivity was associated with overall survival for many drugs in SCAN-B and METABRIC, two cohorts on different platforms with more than 1,100 deaths each, and the directions largely agreed, which argues against a platform artifact. Much of this signal is prognostic and shared: a proliferation-linked axis accounts for part of it, and most associations point in the direction of the cytotoxic controls. The controls are weak evidence, because more than a third of all drugs reach significance and the taxane associations were as strong in patients without chemotherapy. Because every cohort uses the same TF representation, training cell lines and Ridge specification, with per-cohort refits on the shared TFs, the concordance supports the consistency of the transfer more than the drug specificity of each association.

The rule-based candidates should be read in this light. For five of the seven candidates basal-like tumors were predicted less sensitive than luminal A tumors, so in an unselected cohort the HR *>* 1 filter, which selects drugs whose predicted sensitivity is associated with better survival, tends to favor luminal phenotypes over the TNBC phenotype. The associations also shrank after clinical adjustment, and for three candidates the transfer model barely ranks cell lines by response. Pelitinib and osimertinib, the two candidates for which basal-like tumors were not predicted less sensitive than luminal A tumors, were the only ones that kept a significant association within the triple-negative tumors of SCAN-B. Four of the candidates are annotated as EGFR inhibitors, and four are approved drugs: osimertinib (Soria et al., 2018) and erlotinib (Shepherd et al., 2005) for non-small cell lung cancer (NSCLC), brigatinib for ALK-positive NSCLC (Camidge et al., 2018) and trametinib for BRAF-mutant melanoma (Flaherty et al., 2012). EGFR is over-expressed in roughly half of basal-like tumors (Nielsen et al., 2004; Carey et al., 2012), but EGFR-directed antibodies combined with platinum gave modest responses in unselected TNBC, and the randomized trial did not meet its primary endpoint (Carey et al., 2012; Baselga et al., 2013). Single-agent Src inhibition was similarly inactive (Finn et al., 2011; Gucalp et al., 2011). Osimertinib was designed to inhibit mutant EGFR while sparing the wild-type receptor (Cross et al., 2014). The candidates are therefore hypotheses rather than treatments supported by these data. Before any clinical testing they would need evidence that TNBC models respond to them and that a TF-based score identifies the responders.

Several limitations should be noted. The survival screen and the candidate rule used breast cancer not selected for TNBC, in which basal-like tumors are 9.5% of the SCAN-B Cox cohort, and none of the candidate drugs is a standard breast cancer treatment. The screen of 1,447 drugs adjusts only for PAM50 subtype, with age, stage and treatment adjusted only for the 12 candidates and controls, and its endpoint is overall survival, so its counts can include associations with deaths from other causes. No clinical expert reviewed the candidates before selection, so the list carries no judgment of clinical feasibility beyond the development-stage filter. PRISM and GDSC recovered few established effects in the TNBC contrasts (two of 17 checks), so the null comparison in the TNBC lines carries little weight on its own.

The per-drug Ridge models used for clinical transfer have modest accuracy on held-out cell lines (median per-drug r = 0.086), so a survival association with a drug’s predicted AUC is not an association with sensitivity to that drug. In the canonical model the PAM50 terms are penalized and violate proportional hazards, and the stratified sensitivity analysis covers only the candidates and controls.

In summary, the kinase and HDAC inhibitors selected from these associations, four of them annotated as EGFR inhibitors, rest on prognostic associations in unselected breast cancer. Neither the TNBC cell lines nor FUSCC-TNBC supported them, although both analyses could detect only large effects, and their relevance to TNBC remains to be tested experimentally.

## Methods

### Data sources and preprocessing

We obtained drug sensitivity data from PRISM (476 cell lines *×*1,448 drugs, AUC values, Corsello et al., 2020) and GDSC release 8.5 (696 cell lines *×* 397 drugs, Yang et al., 2013). Cell-line gene expression (log_2_(TPM+1), 19,193 protein-coding genes) was obtained from DepMap release 24Q4 (Ghandi et al., 2019; Broad DepMap, 2024). Cell-line annotations (lineage and receptor subtype, used to select the breast and TNBC lines) were taken from the DepMap 25Q3 model table. TF-target interactions were obtained from Collec-TRI (Müller-Dott et al., 2023). The March 2026 snapshot, used for the knowledge graph, the cell-line TF activities, AURORA-US and FUSCC-TNBC, comprises 42,990 signed regulatory edges connecting 1,185 TFs to 6,675 target genes. The SCAN-B, METABRIC and TCGA-BRCA TF-activity matrices were computed in April 2026 with a CollecTRI network retrieved separately through decoupleR, and Supplementary Note 2 quantifies the effect of recomputing them with the snapshot. Curated protein-protein interaction (PPI) data were obtained from OmniPath (Türei et al., 2021), yielding 84,587 interactions. Mechanism of action (MOA) categories and molecular targets were extracted from the PRISM treatment metadata (1,079 drugs with MOA annotations across 155 categories after filtering out MOAs with fewer than three drugs). All cell lines were checked by Cellosaurus accession against the International Cell Line Authentication Committee (ICLAC) register (version 14). None of the 12 TNBC lines used for Integrated Gradients is listed, and the 13 listed lines among the 476 PRISM cell lines (Supplementary Table S4) were retained because the models receive no tissue or lineage label as input. No cells were cultured in this study. Cell-line data were used as distributed in the DepMap, PRISM and GDSC releases, and the lines were not authenticated or tested for mycoplasma by the authors. We used five patient cohorts: three for the survival screen (SCAN-B, METABRIC and TCGA-BRCA), AURORA-US for the comparison of primary and metastatic tumors, and FUSCC-TNBC, a triple-negative cohort, to examine the screen and the candidates within TNBC. SCAN-B (Saal et al., 2015; Staaf et al., 2022) provided FPKM RNA-seq adjusted for library preparation protocol to a TruSeq-like baseline (Mendeley Data, Vallon-Christersson, 2023) for 8,438 samples from Swedish breast cancer patients (7,397 unique patients with overall survival after deduplication, of whom 7,215 with a PAM50 subtype entered the PAM50-adjusted Cox models). METABRIC (Curtis et al., 2012), obtained from cBioPortal (Cerami et al., 2012), provided Illumina HT-12 microarray expression (log_2_ intensities) for 1,980 patients, 1,979 of them with overall-survival follow-up. The query was restricted at download to the CollecTRI target genes, so the matrix holds 6,549 genes, covering 6,519 of the 6,675 targets of the snapshot. TCGA-BRCA (The Cancer Genome At-las Network, 2012) provided RNA-seq TPM expression from the Genomic Data Commons (GDC) SummarizedExperiment built with TCGAbiolinks (Colaprico et al., 2016) for 1,231 samples (1,072 unique patients with primary tumors and overall survival after filtering and deduplication, of whom 1,060 with a PAM50 subtype entered the PAM50-adjusted Cox models, with 149 events). The RNA-seq matrices were transformed via log_2_(*x* + 1) before TF activity inference, to match the DepMap log_2_(TPM+1) scale used for cell-line training. The METABRIC matrix already arrives as log_2_ intensities and was used as distributed. The AURORA-US study (Garcia-Recio et al., 2023) (Gene Expression Omnibus, GEO: GSE209998) provided upper-quartile normalized RNA-seq expression for 129 samples (44 primary, 79 metastatic and 6 normal) from 53 patients in the GEO deposit, compared with 55 patients in the original publication. Of these 53 patients, 39 had both primary and metastatic samples available for paired analysis. FUSCC-TNBC (Jiang et al., 2019) provided FPKM RNA-seq expression (hg38) for 360 patients, taken from the figshare deposit of its data descriptor (Chen et al., 2022a), with clinical annotations from Table S1 of the original article, and its FPKM matrix was transformed via log_2_(*x* + 1) like the other RNA-seq matrices. Its RNA-seq libraries, unlike those of SCAN-B, were prepared from total RNA with ribosomal depletion. Two patients with a relapse-free survival time of zero were excluded, as were two who did not meet the definition of invasive triple-negative disease, one with microinvasive ductal carcinoma in situ and one with equivocal HER2 immunohistochemistry (2+) and no fluorescence in situ hybridization result. The remaining 356 patients had 49 relapses, defined in the source study as a first recurrence or a contralateral breast cancer, and a median follow-up of 47.6 months. Sex was not used in any analysis. All METABRIC and FUSCC-TNBC patients and all AURORA-US samples are recorded as female, the 12 male patients of TCGA-BRCA had no PAM50 subtype and therefore did not enter the Cox models, and the SCAN-B clinical table has no sex variable, so the findings apply to female breast cancer. The ancestry composition of the clinical cohorts was not assessed.

### Heterogeneous biomedical knowledge graph

We constructed a heterogeneous graph *G* = (*V, E*) with four node types and four edge types (Figure 1a). The graph contains genes (n = 20,389), TFs (n = 1,185), drugs (n = 1,448) and cell lines (n = 476) as nodes. Edges encode TF-gene regulatory interactions (42,990, signed), gene-gene edges from PPI between the encoded proteins (169,174, bidirectional, over 83,176 distinct gene pairs), drug-gene target interactions (3,442) and cell line-drug response edges (614,325, weighted by AUC).

Cell lines were represented by their TF activity profiles (771 dimensions, computed via ULM from decoupler in Python with at least five targets per TF, Badiai-Mompel et al., 2022) concatenated with expression of the top 500 most variable genes (1,271 total features). Drug nodes used multi-hot MOA encoding (155 dimensions) combined with learned per-drug embeddings (128 dimensions) to ensure unique drug representations. Gene and TF nodes used their mean expression and activity values over all 476 cell lines, and the 500 most variable genes were also selected over all 476, so these unsupervised features include the expression of the held-out cell lines but none of their responses.

### GNN architecture and training

We implemented a heterogeneous GNN using Graph-SAGE convolutions (Hamilton et al., 2017) with three layers, 128 hidden dimensions, per-node-type input projections, reverse edges for bidirectional message passing and learned drug embeddings, using PyTorch Geometric (Fey and Lenssen, 2019). Cell-line and drug embeddings were concatenated and passed through a three-layer multilayer perceptron to predict AUC. A Heterogeneous Graph Transformer (HGT, Hu et al., 2020) variant with two layers and four attention heads was also evaluated. Models were trained on the full batch of training response edges with Huber loss, Adam optimizer (Kingma and Ba, 2015) (weight decay 10^*−*4^) with cosine annealing learning-rate schedule (Loshchilov and Hutter, 2017) (800 epochs, learning rate 1.3 *×*10^*−*3^, Supplementary Figure S1), dropout of 0.33 for Graph-SAGE and 0.20 for HGT, and response edges excluded from message passing to prevent label leakage. Because the response edges are the only edges of the cell lines, the cell lines receive no messages, and their representation is a learned projection of their own features. The weights of the final epoch were kept, so the test cell lines were never used to select a checkpoint.

The GNN rows of Table 1 give the mean and standard deviation over five initialization seeds (42, 123, 456, 789 and 1011) on the same split. Each GNN of Table 1, the graph ablation and the attribution analyses was trained once, and the saved models were reused. These models were trained on a single NVIDIA GeForce RTX 5060 Laptop GPU. The PRISM-to-GDSC cross-screen evaluation used a separate HGT without learned drug embeddings, trained on a random 80/20 split of the PRISM response edges (seed 42).

To measure the contribution of each edge type, we retrained GraphSAGE with the same split, protocol and five initialization seeds after removing the CollecTRI, PPI and drug-target edges individually, the PPI and drug-target edges together, and all three together. As a control for the drug-target relation, we kept it and rewired its edges by degree-preserving swaps (Maslov and Sneppen, 2002), so that each drug kept its number of targets and each gene the number of drugs that target it, with a different rewiring for each seed, in which 1.6 to 1.9% of the edges matched an original drug-target pair. For each configuration we report the mean and standard deviation over the five seeds of the global and median per-drug test Pearson correlation (Supplementary Figure S12).

### Baseline and reference models

All models were evaluated in a single cell-line hold-out split (80/20, 381 training and 95 test cell lines) drawn at random with a fixed seed (42). Per-drug and global test metrics of every model were computed over the same 1,447 drugs with at least 30 training and 5 test cell lines. Test cell lines were never seen during training, unlike edge-level splits, in which a cell line can appear in training and test with different drugs. Three baselines were compared against the GNN, all implemented in scikit-learn (Pedregosa et al., 2011). The first was Ridge regression on full gene expression (19,193 standardized features, *α* = 506), and the second a Random Forest on TF activities (771 features, 150 trees, maximum depth 8, at least 13 samples per leaf), fitted with the five seeds of the GNNs. The third was a hybrid Ridge regression (*α* = 100) on the concatenation of gene expression features and the cell-line embeddings of each of the five GraphSAGE models.

The hyperparameters of Ridge on expression and the Random Forest were selected within the 381 training cell lines. Ten configurations per model were drawn at random from the search space (Optuna, Akiba et al., 2019, seed 42): *α* log-uniform between 10^*−*3^ and 10^3^ for Ridge and, for the Random Forest, 50 to 300 trees in steps of 50, a maximum depth of 3 to 20 and a minimum of 2 to 20 samples per leaf. Each configuration was scored by five-fold cross-validation, with the lines of each donor in the same fold and the folds stratified by lineage, as the mean over folds of the median per-drug correlation over the 200 drugs with most training measurements. The best *α*, 506.2, was rounded to 506 (cross-validated median correlation 0.1622*±* 0.0155, mean*±* standard deviation over folds), and the best Random Forest had 150 trees, a maximum depth of 8 and a minimum of 13 samples per leaf (0.1794 *±*0.0287). The search was run on the supercomputing infrastructure of the Galician Supercomputing Center (CESGA). Supplementary Table S5 lists every configuration, and the values of version 1 of the bioRxiv preprint of this work (https://doi.org/10.64898/2026.09.08.750045), which were chosen on the 95 test lines. During development, the GraphSAGE hidden size, depth, dropout, learning rate and number of epochs came from the best trial of a search with a fixed cosine schedule that maximized global correlation on a random edge-level split, in which every cell line, including the 95 now held out, contributed training and evaluation edges. The HGT used the default architecture of an earlier version of the code and the optimizer settings selected for GraphSAGE. The *α* of the hybrid and transfer Ridge models was fixed at 100, not searched. Two reference predictors were added: the drug mean, which predicts each test pair with the drug’s mean training AUC without cell-line information, and the per-drug Ridge models on TF activities used for clinical transfer (below), in their SCAN-B version restricted to the 769 TFs shared with that cohort. The between-drug share of the variance over the 122,784 test pairs was computed as the sum of squared deviations of each drug’s test mean from the grand test mean, divided by the total sum of squares.

### TF importance analysis

TF importance was computed via Integrated Gradients (IG) (Sundararajan et al., 2017) as implemented in Captum 0.9 (Kokhlikyan et al., 2020), applied to the trained heterogeneous GNN. For a scalar prediction *f* (*x*_*c*_, *x*_*d*_) at cell line *c* with feature vector *x*_*c*_ and drug *d*, IG assigns to each feature *t* the attribution 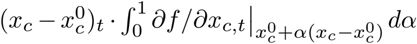, where 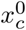 is a reference baseline. IG was applied to the full 1,271-dimensional cell-line feature vector with 50 integration steps and a baseline equal to its mean over the training cell lines of the model’s split, and the attributions of the 771 TF dimensions are reported. In the 24 cell line and drug pairs recomputed for two drugs (erlotinib and epirubicin), these carried on average 26% of the absolute attribution of the full vector, and the rest fell on the 500 expression features. Attributions were computed for the 12 TNBC cell lines paired with the seven repurposing candidates and four positive-control drugs (132 pairs). Talazoparib was added to the positive controls after the attribution analyses had been run. Eight of the 12 TNBC lines belong to the training set of the seed-42 split and four to its test set. Signed attributions are relative to the baseline: a positive attribution for TF *t* and drug *d* means that the activity of *t* in that cell line, relative to the mean training cell line, raises the predicted AUC. Whether higher activity of *t* raises or lowers the prediction is given by the sign of the path-averaged gradient, which we recovered by dividing each attribution by 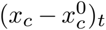 and checked against direct gradients for two drugs. The similarity of the attributions across drugs was measured as the mean pairwise Pearson correlation, over the 11 drugs, of the 771-TF vectors of mean signed attribution across the 12 cell lines.

We evaluated the stability of the ranking along two axes. First, we trained GraphSAGE with three seeds (42, 123 and 456), each seed also defining its own random 80/20 cell-line split, and recomputed IG on each model with the mean-train baseline of its own split. These seed-123 and seed-456 models therefore differ from the seed-123 and seed-456 models of Table 1, which share the seed-42 split. We report the TFs in the top 20 of all three seeds, and we define the core set as the ten TFs with the lowest mean IG rank across the three training seeds. The seed-42 model is the same model in every attribution analysis. Second, with that model, we compared the mean-train baseline against a zero baseline and against the importances averaged over three runs with random Gaussian baselines drawn with the per-feature mean and standard deviation of the training cells. Ranking stability was quantified by Spearman’s *ρ* over the 771 TFs and by the top-20 overlap, which for two random lists is 20 *×* 20 / 771 = 0.52 TFs on average.

### Clinical transfer via Ridge regression

To transfer predictions from cell lines to patients, we adopted a per-drug Ridge regression approach on TF activity profiles. For each clinical cohort except AURORA-US, TF activities were computed from expression data using decoupleR ULM (R) with CollecTRI. For each cohort and each drug with sufficient training data (n *≥*30 training cell lines, 1,447 of the 1,448 drugs), a Ridge regression model (*α* = 100) was trained on PRISM train cell-line TF activities and applied to patient TF activities. Each model used only the TFs shared between PRISM and that cohort (769 in SCAN-B, 767 in METABRIC, 771 in TCGA-BRCA and 769 in FUSCC-TNBC). The FUSCC-TNBC activities were computed with the CollecTRI snapshot over the genes the cohort shares with the PRISM expression matrix (Supplementary Note 2). Before fitting, a QuantileTransformer mapping each TF activity to a normal distribution and a standardization step were fitted on the training cell lines and applied unchanged to the cell lines and to each cohort. This places the patients on the cell-line scale but does not align their distributions with those of the cell lines, and patient values beyond the cell-line range are set to the bounds of the transform. Each drug therefore has one model per cohort, and the GNN does not take part in the transfer.

### Survival analyses

Survival associations used penalized Cox proportional hazards regression (Cox, 1972) (penalizer *λ* = 0.1) implemented in the lifelines Python library (Davidson- Pilon, 2019), with predicted AUC as a continuous covariate, adjusting for PAM50 molecular subtype. In METABRIC the screen was also repeated with disease-specific survival as the outcome, censoring deaths from other causes. Hazard ratios are per unit of predicted AUC, and their confidence intervals and Wald p-values come from the penalized fits and are approximate. The lifelines library applies the penalty to every coefficient on the standardized scale, including the PAM50 terms. P-values were corrected for multiple testing using the Benjamini-Hochberg procedure (Benjamini and Hochberg, 1995) within each cohort over the 1,447 drugs, and the Benjamini-Yekutieli procedure (Benjamini and Yekutieli, 2001) is reported alongside. SCAN-B was deduplicated by patient ID before analysis (from 8,269 clinical records, one per sequencing library, to 7,662 unique patients), and TCGA-BRCA was filtered to primary tumors only and deduplicated by submitter ID (from 1,231 samples to 1,095 patients). For SCAN-B and TCGA-BRCA, PAM50 subtypes were available from the original clinical annotations. For METABRIC, PAM50 subtypes were computed from the expression matrix using the PAM50 classifier (Parker et al., 2009) of the genefu R package (Gendoo et al., 2016) on the 39 of the 50 PAM50 genes present in the target-restricted matrix, achieving 73% concordance with the original PAM50 and claudin-low classification over 1,756 patients (excluding claudin-low and unclassified samples, which have no PAM50 equivalent, Prat et al., 2010). The PAM50 adjustment in METABRIC is therefore less complete than in SCAN-B. To express effect sizes on a common scale, we converted each SCAN-B hazard ratio to the scale of one standard deviation of the drug’s predicted AUC across the 7,215 patients of the Cox cohort, which is exact for the penalized fit because lifelines penalizes the standardized coefficient.

Proportional hazards were assessed with the score test on scaled Schoenfeld residuals (Grambsch and Therneau, 1994) of the survival R package (version 3, rank-transformed time pooled across strata), reimplemented in Python and checked against R. The test was applied to every term of the canonical models of the candidates and controls and of their unpenalized refits in SCAN-B, METABRIC and TCGA-BRCA, with the METABRIC results in Supplementary Table S1 and the TCGA-BRCA results in Supplementary Note 2. Because PAM50 terms violated the assumption in SCAN-B and METABRIC, we refitted the models of the candidates and controls of both cohorts with PAM50 as a stratification factor, and tested these fits in the same way, penalized as above and unpenalized with the predicted AUC standardized.

Cross-cohort meta-analysis used Fisher’s method to combine p-values across the three cohorts for each drug, using the upper tail of the *χ*^2^ distribution with 2*k* degrees of freedom for *k* cohorts, followed by Benjamini-Hochberg correction. As a direction-aware sensitivity analysis we combined signed z-scores, *z* = sign(log HR) Φ*−*1(1 *−p/*2), with a Stouffer test weighted by the square root of the number of events of each cohort. Directional agreement between SCAN-B and METABRIC was compared with the agreement expected if the directions were independent, given the proportion of hazard ratios above 1 in each cohort. For the positive controls, the binomial probability that at least as many of the five controls reach Fisher FDR *<* 0.05 as observed was computed with the fraction of all 1,447 drugs reaching it as the success probability, and each control was ranked among all drugs by its Fisher p-value. All statistical tests were two-sided (Wald tests for Cox coefficients, log-rank and Cox score tests and paired Wilcoxon signed-rank tests), except the Fisher’s exact tests of the enrichment analysis, the hypergeometric test of the AURORA-US core TFs, the binomial test of the positive controls and the permutation test of the FUSCC-TNBC correlation, which were one-sided upper-tail tests. Genomic-control inflation factors (*λ*_GC_) were computed per cohort as the median of the *χ*^2^ statistics implied by the per-drug p-values divided by the median of the *χ*^2^ distribution with one degree of freedom (Supplementary Figure S5). Median follow-up was estimated with the reverse Kaplan-Meier method.

The Kaplan-Meier panels split patients at the median predicted AUC and compared the halves with a two-sided log-rank test, and with a PAM50-stratified score test among patients with a subtype. The trend across quintiles of predicted AUC was tested with the score test of a Cox model with the quintile index (1 to 5) as a numeric covariate, equivalent to a log-rank test for trend. The numbers at risk count, at each two-year mark, the patients whose follow-up reached that time.

To assess whether the survival associations reflect a shared prognostic axis rather than drug-specific effects, we performed principal component analysis on the standardized matrix of per-drug predictions from SCAN-B. The first component (PC1) score per patient, oriented so that it correlates positively with the predicted AUC of paclitaxel, was tested against overall survival with the same PAM50-adjusted penalized Cox model and correlated with the SCAN-B TF activities. PC1 was residualized (by ordinary least squares) from each drug’s prediction vector before recounting nominal (p *<* 0.05) per-drug survival associations, on the 200 most variable drugs, which do not include the seven candidates. The empirical inter-drug prediction correlation was computed as the mean and the mean absolute off-diagonal Pearson correlation of the prediction matrix. As a check of the clinical TF activities, each of the 769 SCAN-B TF activities was tested against overall survival with an unpenalized univariate Cox model, one sample per patient and without subtype adjustment or correction for multiple testing.

Treatment and subgroups were analyzed for the seven candidates and five positive controls in SCAN-B with unpenalized Cox models and the predicted AUC standardized over the Cox cohort, with PAM50 as strata in the treatment models. The clinically adjusted model added age, tumor stage, grade, nodal status, ER and PR status, chemotherapy and endocrine therapy (6,231 patients), with missing receptor or treatment status coded as its own category and complete cases for the other covariates. In METABRIC the same analyses used age, positive lymph nodes, the Nottingham Prognostic Index, ER status, HER2 copy-number gain, chemotherapy and hormone therapy (1,903 patients), and the chemotherapy interaction was tested in the 1,979 patients with known chemotherapy status. Effect modification was tested with the product of the standardized predicted AUC and chemotherapy, in patients with known chemotherapy status (6,843 in SCAN-B), first with PAM50 as strata and then with age, tumor stage, grade, nodal status, ER and PR status and endocrine therapy added as main effects (6,173 patients). The triple-negative subgroup comprised the patients of the 7,215-patient Cox cohort with ER-, PR- and HER2-negative tumors, all three known (676 patients, 183 events), and the basal-like subgroup the PAM50 basal-like patients (686 patients, 167 events).

We tested whether each PRISM MOA class annotated for at least five of the 1,447 tested drugs (93 classes) and each molecular target shared by at least three drugs (330 targets) was over-represented among the 551 drugs with a significant Fisher meta-analysis association, with a one-sided Fisher’s exact test and Benjamini-Hochberg correction within each family (Supplementary Figures S8 and S9). Within-subtype screens were run in the PAM50 basal-like subgroup and in the pooled basal-like and HER2-enriched subgroups of SCAN-B, with a median-split log-rank test (Supplementary Figure S13).

### AURORA-US analysis

The AURORA-US cohort was used to assess whether the core TFs are altered during metastatic progression. TF activities were computed using decoupler ULM (Python) with the CollecTRI snapshot. To respect the paired nature of the data (43 of 53 patients contribute multiple samples, with up to eight metastatic biopsies per patient), metastatic samples from the same patient were averaged and a paired Wilcoxon signed-rank test was applied across the 39 patients with both primary and metastatic samples, with Bonferroni correction for the ten core TFs tested. The same test was applied to all 770 TFs to estimate the background rate, and the probability of observing at least as many core TFs below the threshold was computed with the hypergeometric distribution. The site analysis repeated the paired comparison within the patients of each metastatic site with at least three pairs (3 to 12 patients per site) and counted the sites in which each TF kept the direction of the pooled comparison. Drug sensitivity predictions were generated for each AURORA-US sample using the per-drug Ridge models and compared between primary and metastatic samples with the same patient pairing (paired Wilcoxon test, Benjamini-Hochberg correction over 1,447 drugs).

### Rule-based candidate selection

Candidate repurposing drugs were selected by a fixed rule applied to the SCAN-B PAM50-adjusted penalized Cox results, with no expert scoring. The rule has four filters and one cutoff. We report the number of drugs left by each filter and the effect of the rank cutoff (Supplementary Table S2), and we repeat the attribution analysis without the cutoff (Supplementary Note 1). (i) Statistical filter: *p*_adj_ *<* 0.05 in SCAN-B at the canonical penalizer (*λ* = 0.1) with a hazard ratio per unit of predicted AUC in [1.5, 10] (143 drugs). Because the predicted AUC varies little across patients, the drugs that pass these bounds have hazard ratios of only 1.06 to 1.17 per standard deviation. The lower bound therefore removes weak associations and the upper bound removes numerically extreme fits on the scale of the covariate, and neither defines a clinically meaningful effect. (ii) Stability filter: *p*_adj_ *<* 0.05 also at *λ* = 0.5 (52 drugs). (iii) Mechanism class: agents annotated in the PRISM secondary-screen metadata (release 19Q4) as protein kinase or HDAC inhibitors (20 drugs), two of the targeted classes with clinical precedent in breast cancer. PARP inhibitors, another such class, served as positive controls. The filter is applied to each term of the annotation, so that inhibitors of other enzymes whose name contains ‘kinase’ (for example urokinase) do not pass. (iv) Development stage: at least phase 2 in the same annotation, with phase 1/2 agents counted as phase 2 (16 drugs). (v) Rank cutoff: among these, the drugs that fall within the 30 most significant of the 143 drugs of filter (i), yielding the seven candidates of Table 5. MOA, molecular target and clinical phase are taken from the PRISM metadata as distributed and were not updated. The oncology programs of saracatinib and pelitinib have since been discontinued.

### Sensitivity analyses

To assess the robustness of the survival associations and the candidate ranking, we performed three further sensitivity analyses in SCAN-B. First, we replaced overall survival with relapse-free survival as the outcome, for the seven candidates and five controls, with Benjamini-Hochberg correction over these 12 drugs. Second, we swapped the QuantileTransformer for Com-Bat empirical-Bayes adjustment (Johnson et al., 2007) of the joint TF-activity matrix of the PRISM training cell lines and SCAN-B, restricted to their shared TFs. Third, we swept the Cox penalizer over {0.01, 0.1, 0.5, 1.0} to measure how the hazard-ratio ranges and the number of significant drugs depend on regularization. All used the same PAM50-adjusted penalized Cox framework described above, and the ComBat and penalizer analyses covered all 1,447 drugs, with Benjamini-Hochberg correction within each analysis or penalizer. An earlier specification of the clinical adjustment, penalized with PAM50 as covariates and missing receptor status coded as negative, is provided with the code. Its penalty shrinks the clinical covariates together with the prediction, and it retained six of the seven candidates, against two in the unpenalized, PAM50-stratified model reported here.

### TNBC cell-line and FUSCC-TNBC analyses

In PRISM, each measured AUC was converted to its percentile within its own cell line, computed against a fixed reference panel of the 402 drugs measured in at least 95% of the lines. Ties counted at half weight, which applied to the 2.48% of values at exactly 1, the no-effect value of the assay. The percentiles of each drug were standardized across lines, and the composite of a line was the mean of the standardized percentiles of the seven candidates. The composites of the 12 TNBC lines were compared with those of the other 464 lines by Cliff’s delta (Cliff, 1993), the probability that a TNBC line has a higher composite than a comparison line minus the probability of the reverse. Because a lower AUC means a more active drug, a negative delta means greater activity in the TNBC lines. Its 95% confidence interval used the standard error of the delta under the null hypothesis, and its two-sided p came from 10,000 permutations of the cell-line labels, which keep the correlation between drugs intact. The minimum detectable delta at 80% power and a two-sided alpha of 0.05 was derived from that standard error. The models from which the candidates were selected were trained on the same PRISM matrix, so this comparison is not independent of the selection.

For descriptive comparison, the same statistic was computed for each of the 1,409 PRISM drugs measured in at least 8 of the 12 TNBC lines and in at least 30 comparison lines (the evaluable drugs), with its own label-permutation p-value and Benjamini-Hochberg correction, and each drug’s delta was ranked as a percentile of these deltas, so that low percentiles mean greater activity in TNBC. The mean of the per-drug deltas of the seven candidates was also compared, with a two-sided test, against 10,000 random sets of seven other kinase or HDAC inhibitors among the evaluable drugs (348 including the candidates), each matched to a candidate on the number of TNBC lines measured and on the tercile of its share of AUC values at 1 in those lines. To check that the screens recover established effects, the three cytotoxic controls (paclitaxel, docetaxel and epirubicin) were compared in PRISM and in GDSC in three contrasts (TNBC against the other breast lines, all breast lines against the other lineages and TNBC against all other lines), and, in PRISM only, olaparib and talazoparib in the two BRCA1-mutant TNBC lines and erlotinib, osimertinib and pelitinib in the two EGFR-driven TNBC lines, each against the other TNBC lines. The contrast of all breast lines against the other lineages has no expected direction and is given as context in the result tables of the code. Every other check passed when its interval excluded zero in the expected direction, which in the checks of two lines requires both to be more sensitive than every other TNBC line measured. As a calibration of what twelve lines can detect, the BRAF inhibitors dabrafenib and vemurafenib and the MEK inhibitor trametinib were compared between all skin lines and all non-skin lines and between 200 random draws of twelve skin lines and all non-skin lines, with the detection rate defined as the share of draws whose interval excluded zero and a required rate of 0.8. In FUSCC-TNBC, predictions came from the per-drug Ridge models of the clinical transfer. The association of each drug with relapse-free survival was summarized by the standardized score statistic of a univariate Cox model evaluated at a coefficient of zero. This statistic does not change when the predicted AUC is shifted or multiplied by a positive constant, and for a binary covariate without tied event times it equals the log-rank statistic, which we used to check the implementation against lifelines. The primary statistic was the Pearson correlation, over the 1,447 drugs, between these statistics and those computed in the same way for overall survival in the triple-negative subgroup of SCAN-B (676 patients, 183 deaths). It was tested one-sided against 10,000 permutations of the FUSCC-TNBC (time, event) pairs across patients, which keep the correlation between drugs. For a descriptive count of significant drugs, the two-sided score p-values of the 1,447 drugs were corrected with the Benjamini-Hochberg procedure. The secondary statistic was the hazard ratio per standard deviation of the composite of the seven candidates (the mean of their standardized predicted AUC) in an unpenalized Cox model, with a two-sided permutation p of its score statistic. The seven candidates and five controls were also fitted one at a time in univariate, unpenalized Cox models, and the five controls as a composite. To check that the cohort carries an established prognostic signal, we fitted a univariate, unpenalized Cox model of node-positive against node-negative disease, and the check passed if its hazard ratio fell between 2.5 and 3.5. The two patients without a node count were coded as node-negative, with a complete-case refit as a sensitivity analysis, and tumor size and the immunomodulatory subtype were examined in the same way without a criterion. The minimum detectable hazard ratio per standard deviation at 80% power and a two-sided alpha of 0.05 was computed from the number of relapses. Five sensitivity analyses of the twelve drugs and the composite of the candidates were run: stratified by the mRNA subtypes of the cohort, stratified by the intrinsic subtypes, adjusted for age, tumor size and positive nodes in complete cases, without the luminal androgen receptor subtype, and adjusted for the year of surgery. Proportional hazards were assessed with the same score test as in SCAN-B for every term of every FUSCC-TNBC Cox fit (84 fits, 136 terms), one of them a Cox model of relapse-free survival comparing the sequenced patients with the rest of the cohort.

### Statistics and reproducibility

No statistical method was used to predetermine sample sizes, which were set by the public datasets (476 PRISM cell lines and the cohort sizes given above). Supplementary Table S6 summarizes the patients of each survival analysis. Exclusions are described with each dataset and analysis. The main cell-line hold-out (80/20, 381 training and 95 test lines) was drawn once with seed 42, and the PRISM-to-GDSC evaluation used a separate random 80/20 split of the response edges. The GNNs, the hybrid and the Random Forest of Table 1 and the GNNs of the graph ablation were fitted on the main split with five seeds and are reported as mean and standard deviation. The stability analysis of the attributions drew one split per training seed, and the permutations and random draws of the TNBC cell-line and FUSCC-TNBC analyses used seed 42. The folds and the sampler of the hyperparameter search of Ridge on expression and the Random Forest also used seed 42. The study analyzed existing data, so there was no allocation to interventions, and the analyses were not blinded. The tests, their sidedness and the multiple-testing corrections are given with each analysis.

### Ethics

No ethics approval was sought for this study, which is a secondary computational analysis of de-identified, publicly available data and involved no recruitment of, interaction with or new data collection from human participants. The source cohorts were collected under the approval of their own ethics committees, with written informed consent or, for some AURORA-US participants, a waiver of written informed consent, as reported in the original publications. SCAN-B (Clinical-Trials.gov NCT02306096) was approved by the Regional Ethical Review Board in Lund, Sweden (approval numbers 2009/658, 2010/383, 2012/58, 2013/459 and 2015/277). METABRIC samples were obtained with patient consent and approval from ethics committees (REC ref 07/H0308/161). TCGA-BRCA specimens were obtained with consent under the institutional review boards of the contributing sites. AURORA-US was reviewed and approved by the institutional review boards of the participating institutions. FUSCC-TNBC was approved by the Ethics Committee of Fudan University Shanghai Cancer Center, and its patients gave written informed consent (Jiang et al., 2019).

### Use of generative AI

The authors used Claude (Anthropic) while preparing this work, to assist with writing and refactoring the analysis code, with an internal audit of the numerical consistency of the manuscript, with the verification of reference metadata against CrossRef and DataCite, and with drafting and copy-editing parts of the text. The study was conceived, designed and directed by the authors, who made every methodological and editorial decision and interpreted all results, and every number, table and figure reported here is produced by the scripts of the accompanying repository and checked against the manuscript by automated consistency tests. The authors reviewed and edited all content and take full responsibility for it.

## Data availability

All input data are public. Cell-line expression was downloaded from the figshare deposit of DepMap release 24Q4 under CC BY 4.0 (Broad DepMap, 2024), and cell-line annotations from release 25Q3 (Broad DepMap, 2025; Arafeh et al., 2025) (https://depmap.org/portal, accessed September 2026).

Releases up to 24Q4 are deposited on figshare under CC BY 4.0, whereas 25Q3 is distributed through the portal and is governed by its Terms and Conditions. Drug response data come from the PRISM Repurposing secondary screen, release 19Q4, obtained from its figshare deposit under CC BY 4.0 (Broad DepMap et al., 2019), whose drug annotations are curated by the Broad Drug Repurposing Hub (Corsello et al., 2017), and from GDSC release 8.5, whose datasets are now distributed through the Cell Model Passports (https://cellmodelpassports.sanger.ac.uk/downloads). Prior knowledge comes from CollecTRI and OmniPath (https://omnipathdb.org). The CollecTRI snapshot of March 2026, distributed with the code, was used for the knowledge graph, the cell-line TF activities, AURORA-US and FUSCC-TNBC, whereas the SCAN-B, METABRIC and TCGA-BRCA TF activities used here were computed in April 2026 with CollecTRI retrieved through decoupleR (Supplementary Note 2). Clinical cohorts: SCAN-B (Mendeley Data, https://doi.org/10.17632/yzxtxn4nmd.3), METABRIC (cBioPortal study brca_metabric), TCGA-BRCA (GDC Data Portal, project TCGA-BRCA, open-access STAR counts), AURORA-US (GEO accession GSE209998) and FUSCC-TNBC (FPKM matrix from the figshare deposit of its data descriptor, Chen et al., 2022b, under CC BY 4.0, and clinical annotations from Table S1 of the original article, Jiang et al., 2019). The sequencing reads of FUSCC-TNBC are deposited in the Sequence Read Archive (SRP157974), and its omics data and metadata in the National Omics Data Encyclopedia (OEP000155). The per-drug and per-TF result tables generated in this study are provided with the code (release v1.2.0 of the Zenodo archive, https://doi.org/10.5281/zenodo.24203300), and Supplementary Data 1 lists the 143 drugs of the candidate selection.

## Code availability

The code, the per-drug and per-TF tables behind every figure and table, the trained GNN check-points and the statistics file from which every number in the text is drawn (paper_statistics.json, regenerated with sanity checks by pipeline.py) are available at https://github.com/MALL-Lab/PRECISION_paper and archived at Zenodo (release v1.2.0, https://doi.org/10.5281/zenodo.24203300). The code is released under the GPL-3.0 license, and the third-party data files bundled with it retain the licenses of their original sources, documented in the repository. The heterogeneous graph and the OmniPath interaction cache are not redistributed because of the mixed licenses of the OmniPath source databases. The repository provides the scripts that rebuild them (01_prepare_data.py and 02_build_graph.py) and the reference counts of the graph used here (51_graph_summary.py). The clinical TF-activity matrices of SCAN-B, METABRIC, TCGA-BRCA and FUSCC-TNBC are derived from patient-level data and are not redistributed, whereas the per-sample TF activities of AURORA-US, computed from its public GEO deposit, are included. Rebuilding the SCAN-B, METABRIC and TCGA-BRCA matrices from the source cohorts with the distributed CollecTRI snapshot reproduces the results of Supplementary Note 2, while the published results come from the April 2026 matrices and are reproduced by the code from them. The FUSCC-TNBC clinical table is not redistributed, because its publisher’s license does not permit it. The repository provides the scripts that rebuild the cohort and its TF activities (00c_prepare_fuscc.py and 68_-fuscc_tf_activities.R) once the figshare deposit and Table S1 of Jiang et al. (2019) have been downloaded from their sources, and it includes the per-drug FUSCC-TNBC results, which contain no patient-level data.

## ACKNOWLEDGEMENTS

Grant PID2024-162441OA-I00 funded by MICIU/AEI/10.13039/501100011033 and by “ERDF/EU”. RePo-SUDOE, with project reference S1/1.1/P0033, a project co-financed by the Interreg Sudoe Programme through the European Regional Development Fund (ERDF).

This research project was made possible through the access granted by the Galician Supercomputing Center (CESGA) to its supercomputing infrastructure. We acknowledge the Broad Institute for the DepMap portal, the PRISM Repurposing screen and the Drug Repurposing Hub, whose data made this work possible.

The funders played no role in study design, data collection, analysis and interpretation of data, or the writing of this manuscript.

## AUTHOR CONTRIBUTIONS

CFL: conceptualization, methodology, formal analysis, software, funding acquisition and writing of the manuscript. DF: validation, internal review of the analyses and the code, and manuscript revision. PVdR: clinical interpretation of the prioritized candidates and of the survival results, and manuscript revision. All authors read and approved the final manuscript.

## Supplementary Figures

The Supplementary Information comprises 14 supplementary figures, six supplementary tables and two supplementary notes. Supplementary Figures S1 and S2 cover model training (loss and test correlation over 800 epochs) and per-drug prediction quality. Supplementary Figures S3 and S4 show UMAP projections of the GNN cell-line embeddings and the coefficients of the per-drug Ridge transfer models on the core TFs. Supplementary Figures S5 to S9 present the survival analyses: QQ-plots of the survival p-values, the SCAN-B versus METABRIC p-value comparison, the forest plot of the positive controls in SCAN-B and METABRIC, and target and mechanism-of-action enrichment among survival-associated drugs. Supplementary Figure S10 shows the changes in predicted drug sensitivity between primary and metastatic tumors in AURORA-US. Supplementary Figures S11 to S14 report the sensitivity analyses: the Cox penalizer sweep, the graph component ablation, the within-subtype survival screen in the basal-like and HER2-enriched pool and the PRISM-to-GDSC cross-screen evaluation. Supplementary Table S1 reports the replication of the candidates and positive controls in METABRIC, the signed meta-analysis and the interaction with chemotherapy, Supplementary Table S2 summarizes the candidate selection rule and its sensitivity to the rank cutoff (the 143 drugs passing the statistical filter are listed in Supplementary Data 1), Supplementary Table S3 presents sensitivity analyses of the survival associations of the candidates and positive controls in SCAN-B, Supplementary Table S4 lists the 13 cell lines of the knowledge graph found in the ICLAC register, Supplementary Table S5 reports the hyperparameter search of Ridge on expression and the Random Forest of Table 1, and Supplementary Table S6 describes the patients of each survival analysis. Supplementary Note 1 repeats the attribution analysis on the extended candidate set, and Supplementary Note 2 quantifies two dependencies of the clinical results on the TF-activity step, the version of the CollecTRI network and the gene universe each cohort is quantified over.

**Supplementary Figure S1.**
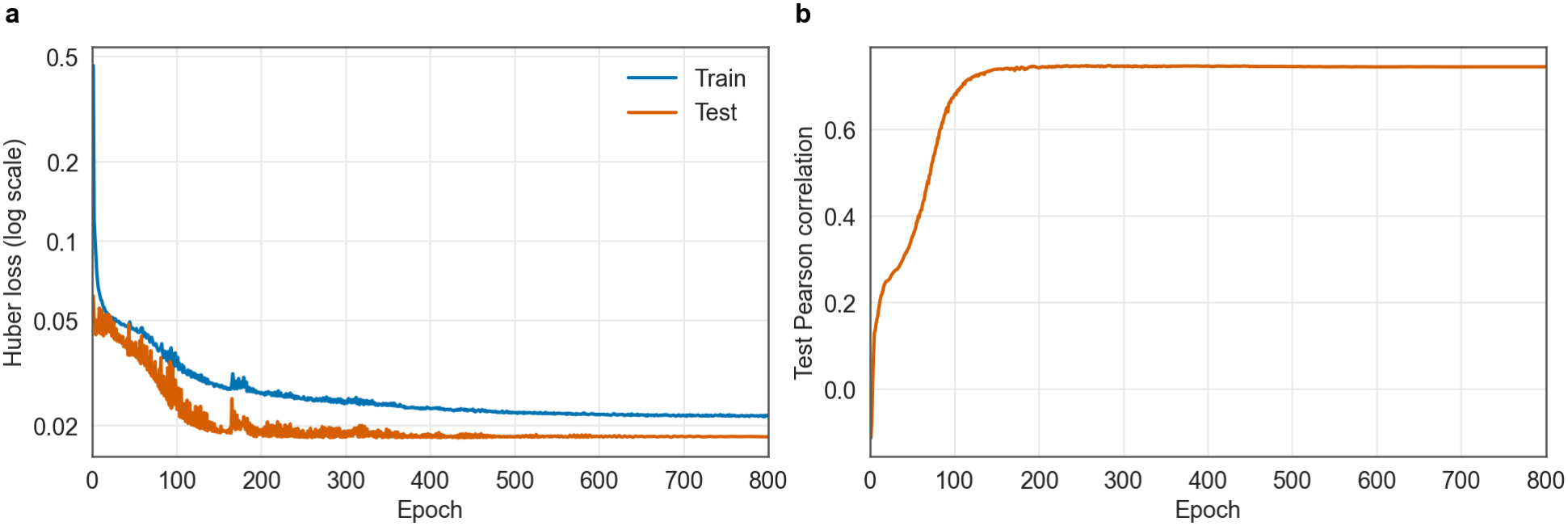
GNN training dynamics of the seed-42 GraphSAGE model, the model used for Integrated Gradients (cell-line hold-out split, 1,447 evaluated drugs). (**a**) Training and test Huber loss (log scale) over 800 epochs with cosine annealing of the learning rate. (**b**) Test Pearson correlation, 0.746 at epoch 800, whose weights are kept (maximum 0.748 at epoch 281, not used to select the model), consistent with the hold-out performance reported in Table 1.

**Supplementary Figure S2.**
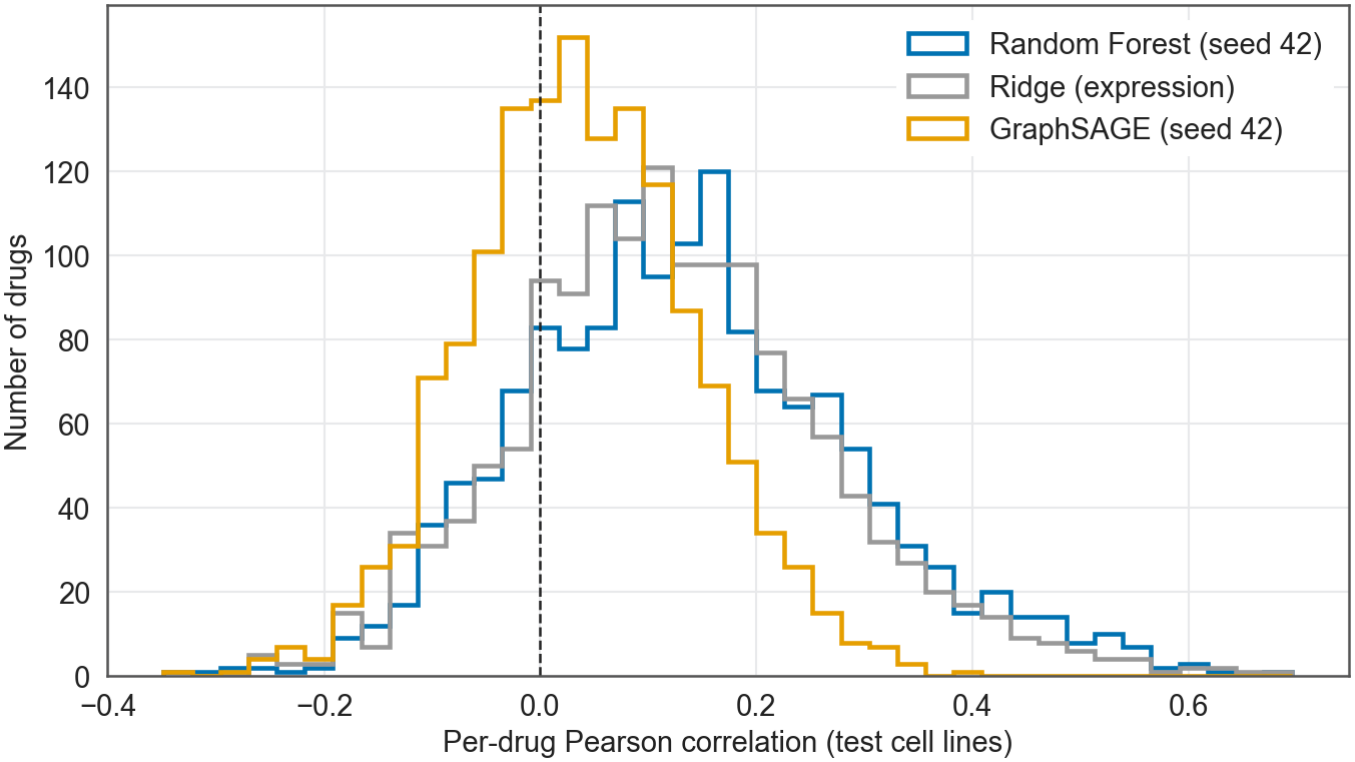
Per-drug Pearson distribution for the seed-42 Random Forest (blue), Ridge (gray) and the seed-42 GraphSAGE GNN (orange) over the same 1,447 drugs, with common bins. Random Forest and Ridge have broad distributions shifted toward positive values, whereas the GNN distribution is narrower and centered close to zero, in line with its lower median per-drug correlation (Table 1).

**Supplementary Figure S3.**
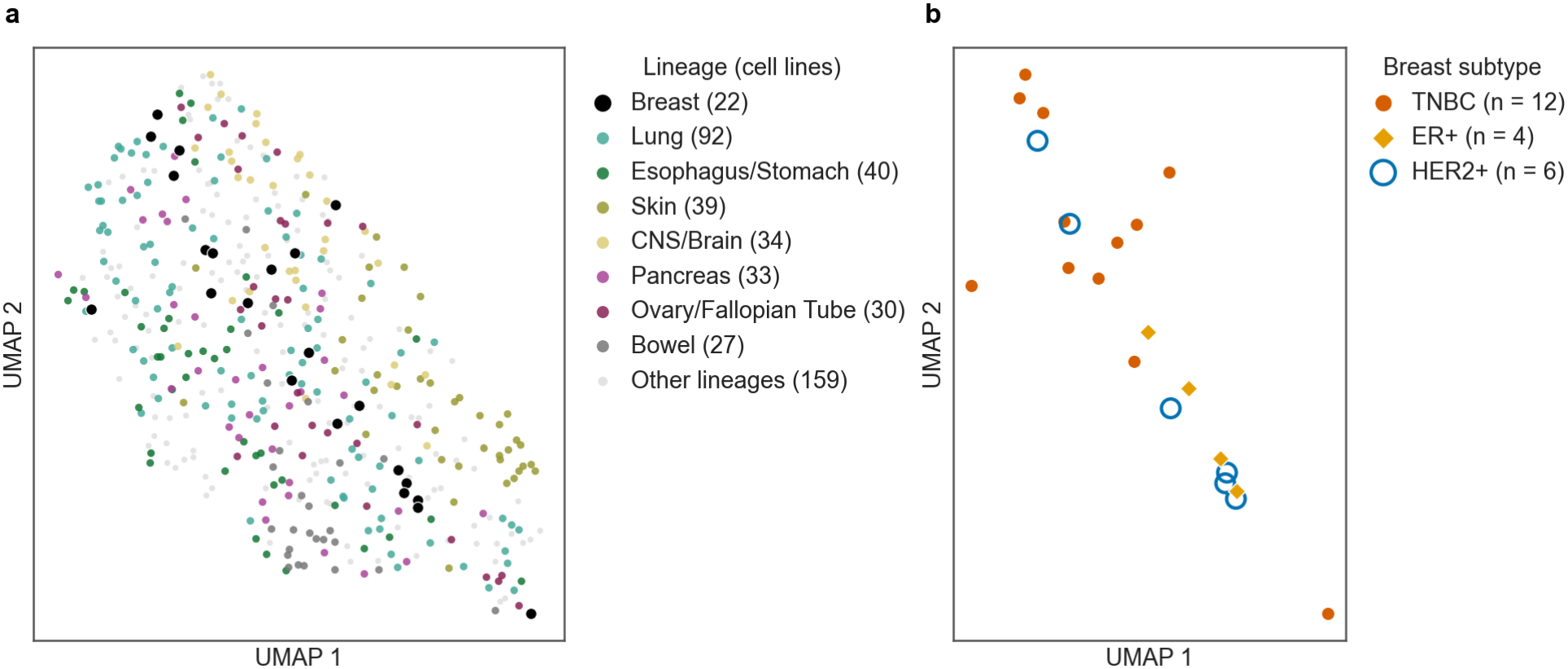
Uniform Manifold Approximation and Projection (UMAP, McInnes et al., 2018) of the cell-line embeddings of the seed-42 GNN. Cell lines receive no messages in the graph, so each embedding is a learned projection of the cell line’s own TF activities and expression. (**a**) All cell lines colored by the seven most frequent cancer lineages other than breast, with breast cell lines in black and the remaining lineages in light gray. Some lineages, such as skin and bowel, form partial clusters. (**b**) Breast cancer cell lines colored by receptor subtype as annotated by DepMap: TNBC (vermillion circles, n = 12), ER+ (orange diamonds, n = 4), HER2+ (blue rings, n = 6). The TNBC lines lie apart from most ER+ and HER2+ lines.

**Supplementary Figure S4.**
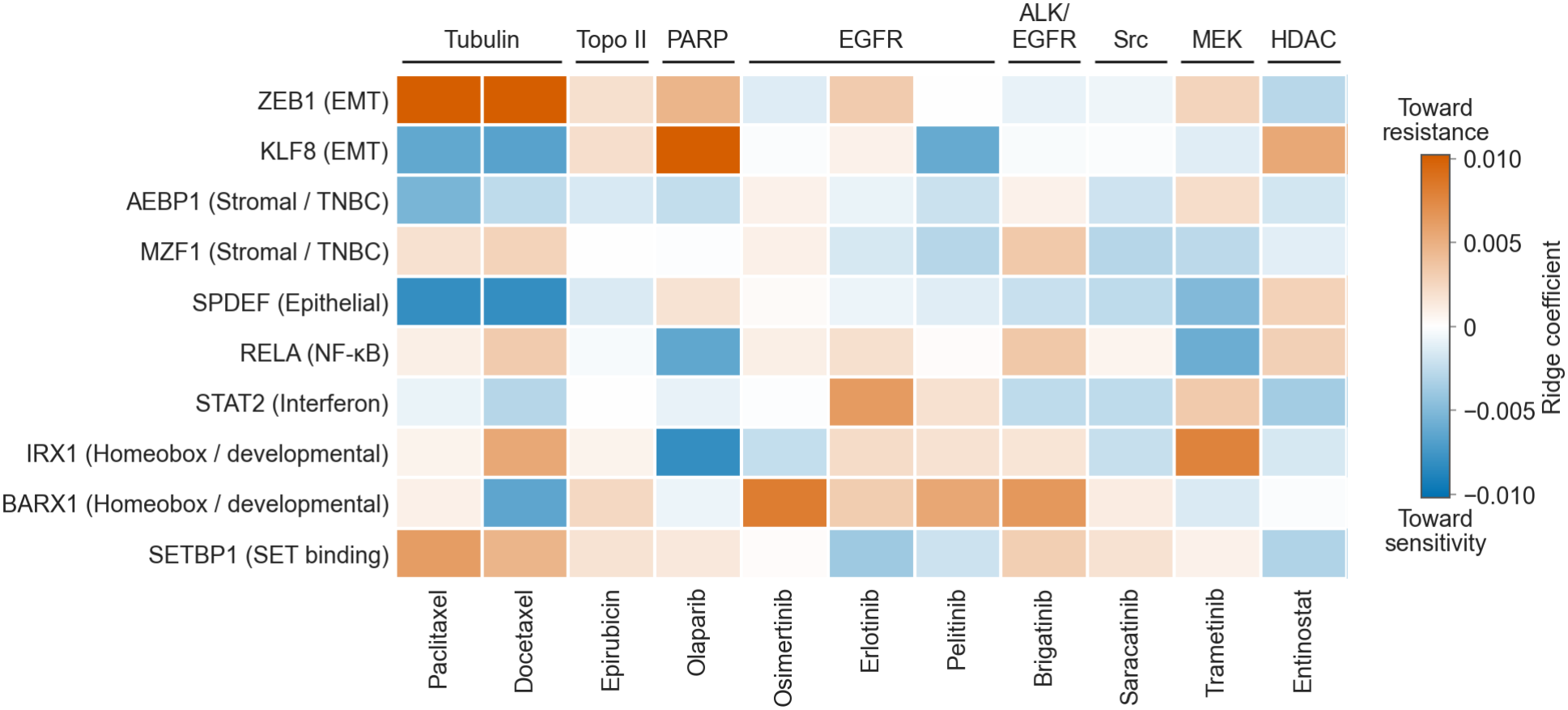
Per-drug Ridge regression coefficients on the TF activities of the ten core TFs, taken from the models that transfer predictions to SCAN-B (*α* = 100, fitted on the 769 TFs shared by PRISM and SCAN-B in the 381 training cell lines after quantile normalization and standardization, Methods) and winsorized at the 2nd and 98th percentiles for display. Drugs are grouped by mechanism of action (MOA): four of the five positive controls (paclitaxel, docetaxel, epirubicin, olaparib) and the seven rule-selected candidates. The core TFs (lowest mean IG rank across the three training seeds, as in Table 4 and Figure 7) are annotated by biological program. Blue marks a negative coefficient (higher activity lowers the predicted AUC, toward sensitivity) and vermillion a positive one (toward resistance). Unlike the GNN attributions of Figure 3c, these coefficients come from separate per-drug models, so the two heatmaps are not expected to agree.

**Supplementary Figure S5.**
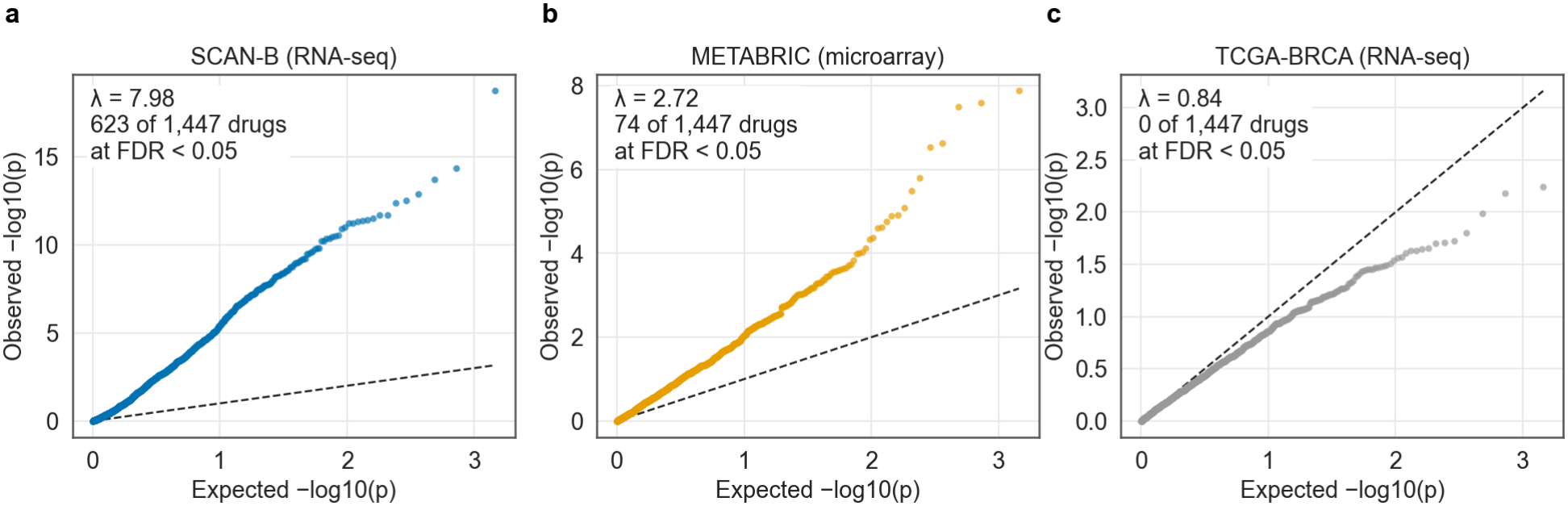
QQ-plots of the per-drug survival p-values in (**a**) SCAN-B, (**b**) METABRIC and (**c**) TCGA-BRCA, with genomic-control inflation factors of 7.98, 2.72 and 0.84. The inflation in SCAN-B and METABRIC reflects widespread, correlated associations, consistent with the shared prognostic axis of the principal component analysis, rather than many independent drug-specific signals. TCGA-BRCA is underpowered.

**Supplementary Figure S6.**
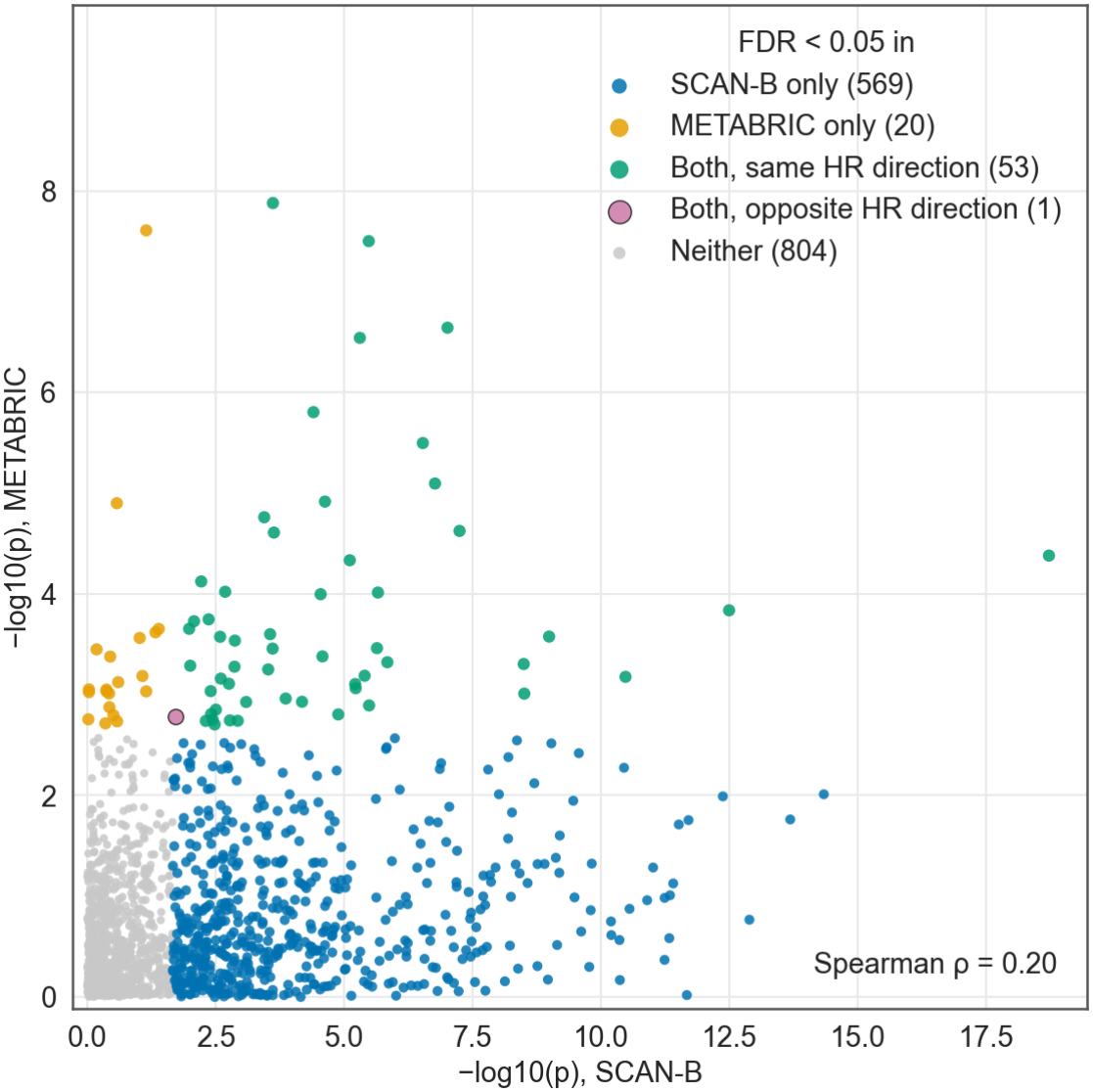
Per-drug survival p-values in SCAN-B versus METABRIC for the 1,447 drugs. The p-values of the two cohorts correlate weakly (Spearman *ρ* = 0.20). Of the 54 drugs significant (FDR *<* 0.05) in both cohorts, 53 (98%) share the direction of the hazard ratio. Colors mark drugs at FDR *<* 0.05 in SCAN-B only (blue), in METABRIC only (orange), in both cohorts with the same hazard-ratio direction (green) or with opposite directions (purple), and in neither (gray).

**Supplementary Figure S7.**
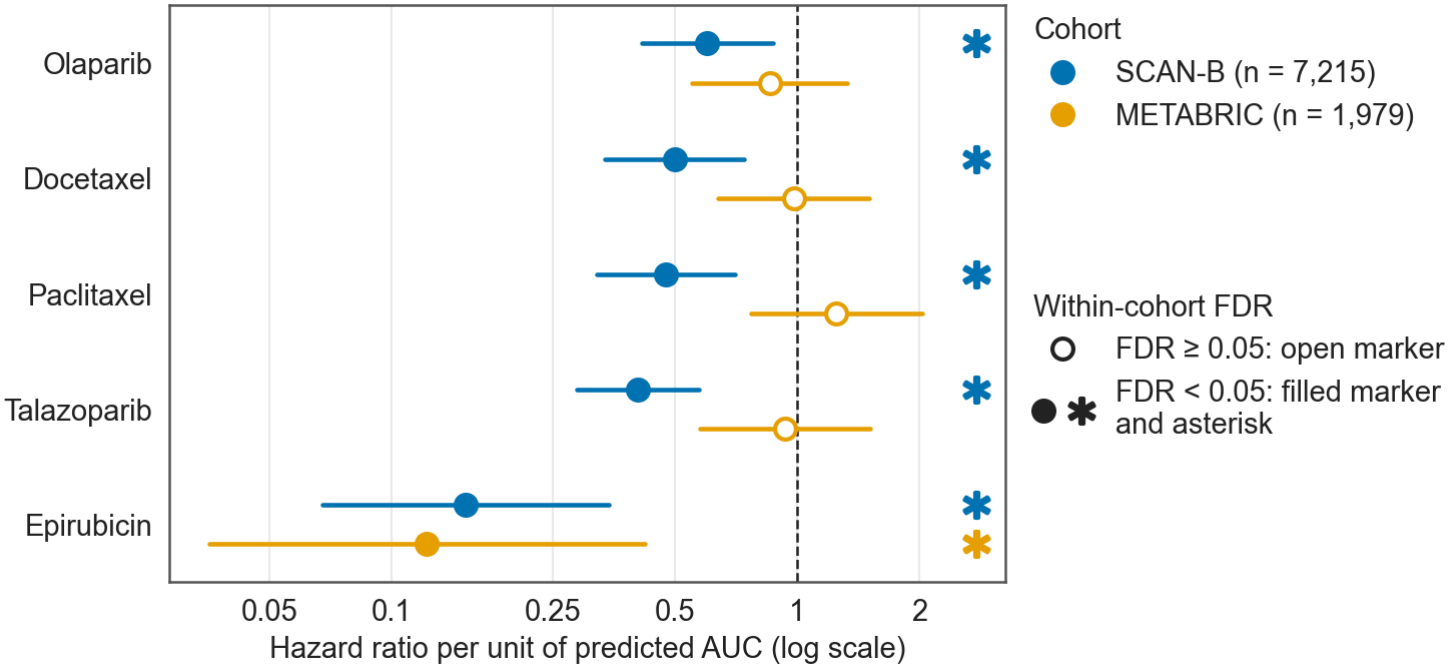
Forest plot of hazard ratios (per unit of predicted AUC, PAM50-adjusted penalized Cox) for the five clinically established breast-cancer drugs across the two adequately powered cohorts. Markers show the point estimate with the 95% confidence interval of the penalized fit, which is approximate. Filled markers with an asterisk indicate FDR *<* 0.05 within the cohort, and open markers FDR *≥* 0.05. All five drugs are individually significant in SCAN-B (n = 7,215), whereas only epirubicin reaches significance in METABRIC (n = 1,979), a cohort with almost as many events (1,143 against 1,164 in SCAN-B). Paclitaxel even shifts above HR = 1 there. TCGA-BRCA is omitted as underpowered (149 events among 1,060 patients, no drug at FDR *<* 0.05, *λ*_GC_ = 0.84, Supplementary Figure S5).

**Supplementary Figure S8.**
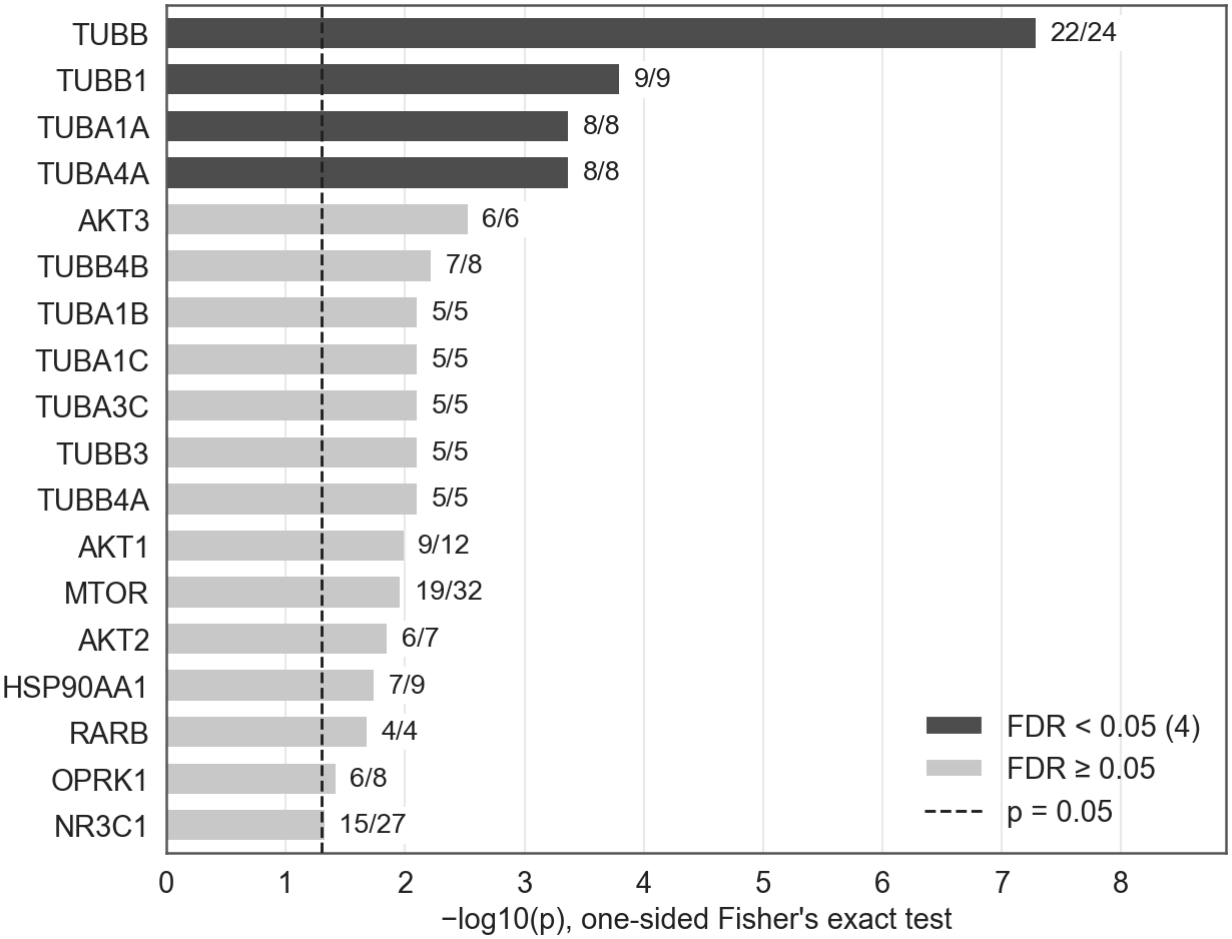
Enrichment of molecular targets among drugs with a significant survival association (Fisher meta-analysis, FDR *<* 0.05), one-sided Fisher’s exact test with Benjamini-Hochberg correction over the 330 targets shared by at least three drugs. The 18 targets with the lowest p-value are shown. The list stops there because the targets that follow are tied in p-value. Four targets are significantly enriched, all of them tubulin subunits (TUBB, TUBB1, TUBA1A and TUBA4A). The drugs annotated to TUBB1, TUBA1A and TUBA4A are a subset of those annotated to TUBB, so the four targets reflect a single tubulin signal, the same one found by the mechanism-of-action analysis (Supplementary Figure S9).

**Supplementary Figure S9.**
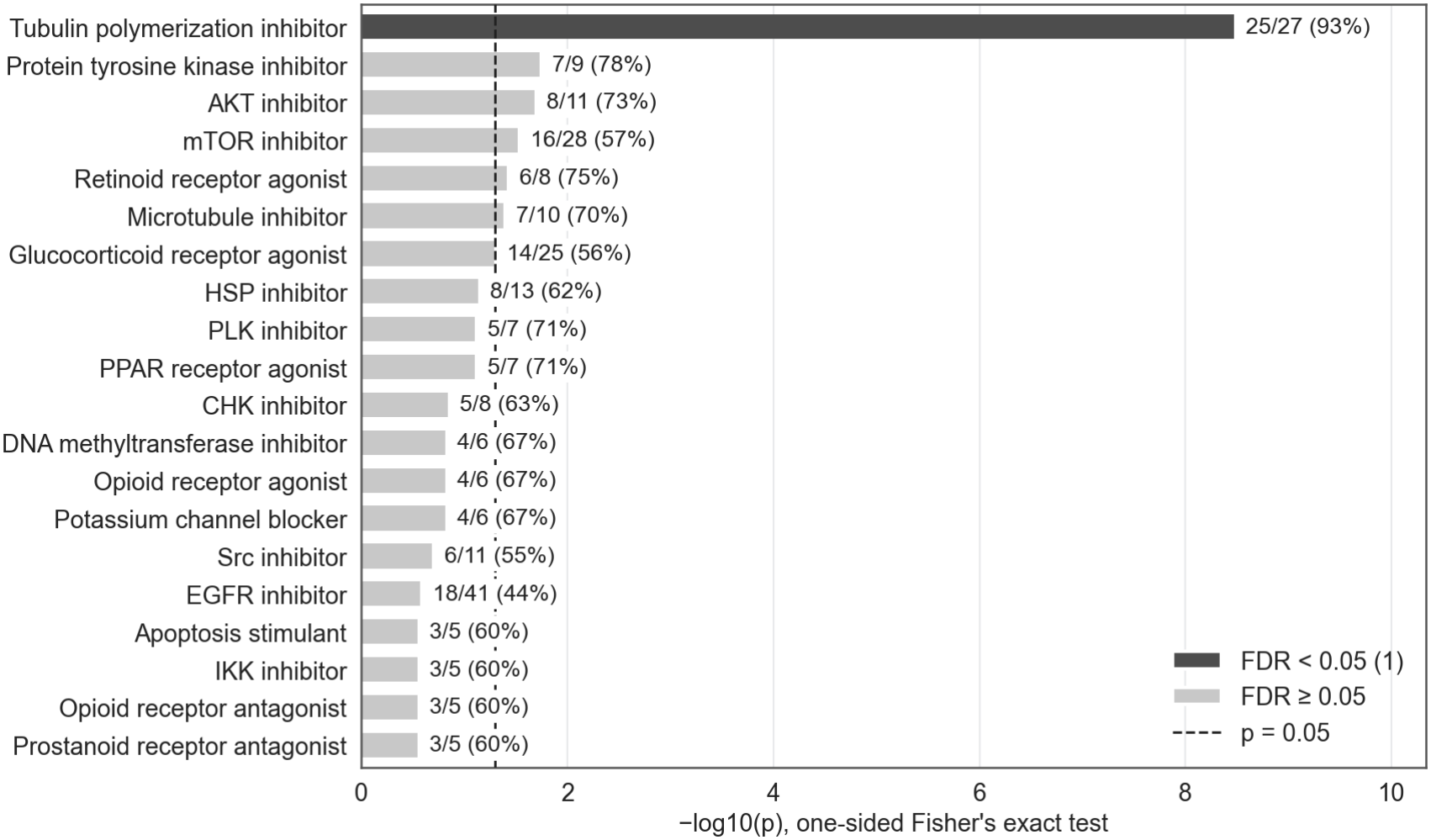
Mechanism-of-action enrichment among drugs with a significant survival association (Fisher meta-analysis, FDR *<* 0.05), one-sided Fisher’s exact test with Benjamini-Hochberg correction over the 93 classes annotated for at least five drugs. The 20 classes with the lowest p-value are shown. Tubulin polymerization inhibitors are over-represented (25 of 27, 93%, FDR = 3.2 *×* 10^*−*7^) and no other class reaches FDR *<* 0.05. The test treats drugs as independent, whereas drugs of one class have correlated predictions, so this single enrichment reflects the proliferation-linked axis shared by antimitotic agents rather than independent convergence of many drugs.

**Supplementary Figure S10.**
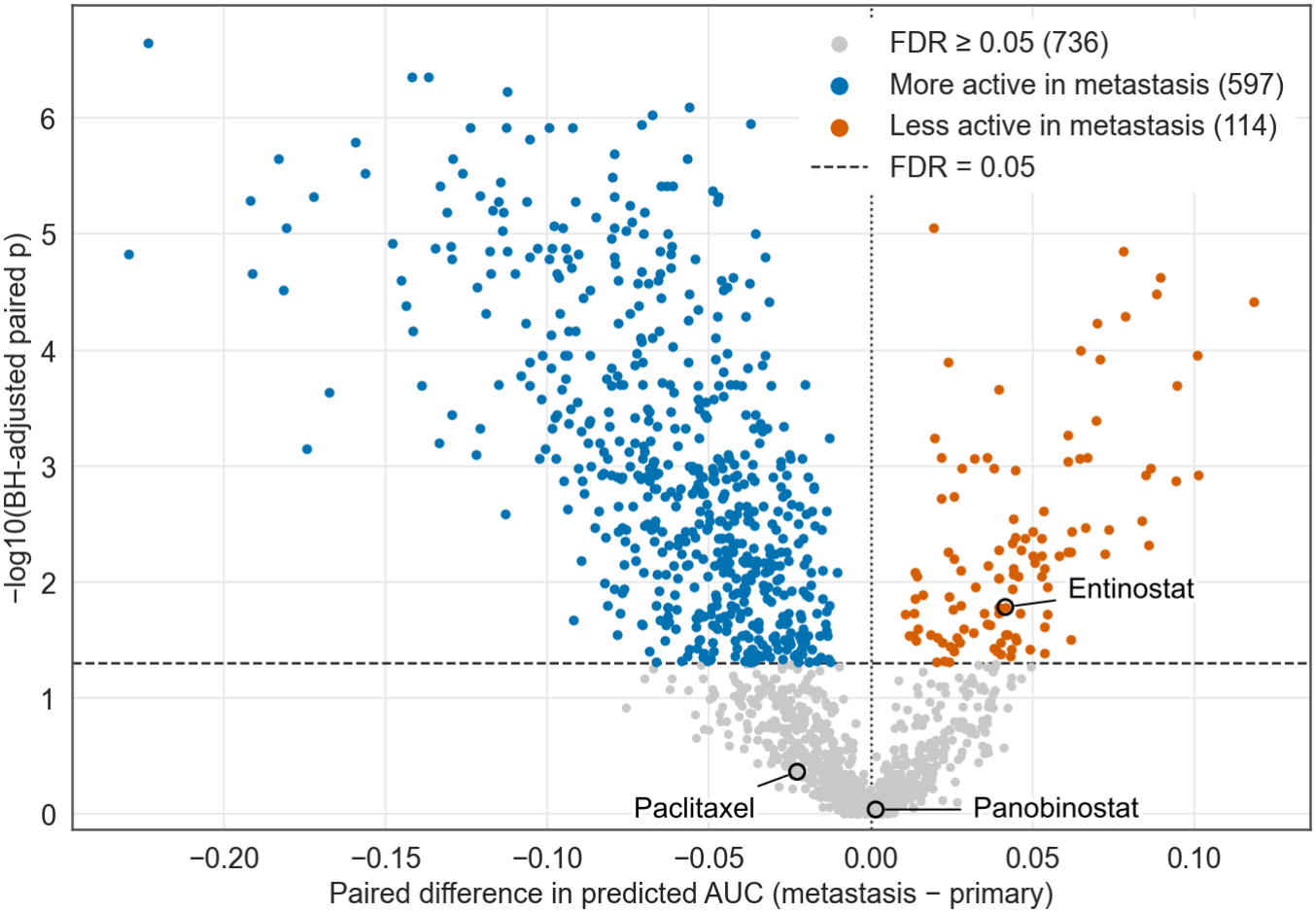
Changes in predicted drug sensitivity between primary and metastatic tumors in AURORA-US, with the patient pairing of the TF analysis (39 patients, metastatic samples averaged per patient, paired Wilcoxon test). The x axis gives the paired difference in predicted AUC (metastasis minus primary) and the y axis the *−* log_10_ Benjamini-Hochberg adjusted paired p-value, with a dashed line at FDR = 0.05. Of the 1,447 drugs, 711 differ at FDR *<* 0.05, 597 of them predicted more active (lower AUC) in metastases (blue) and 114 less active (vermillion), and the rest are gray. Entinostat is predicted less active in metastases (paired p = 6.2 *×* 10^*−*3^), whereas the predictions for paclitaxel and panobinostat do not change significantly (p = 0.33 and 0.88).

**Supplementary Figure S11.**
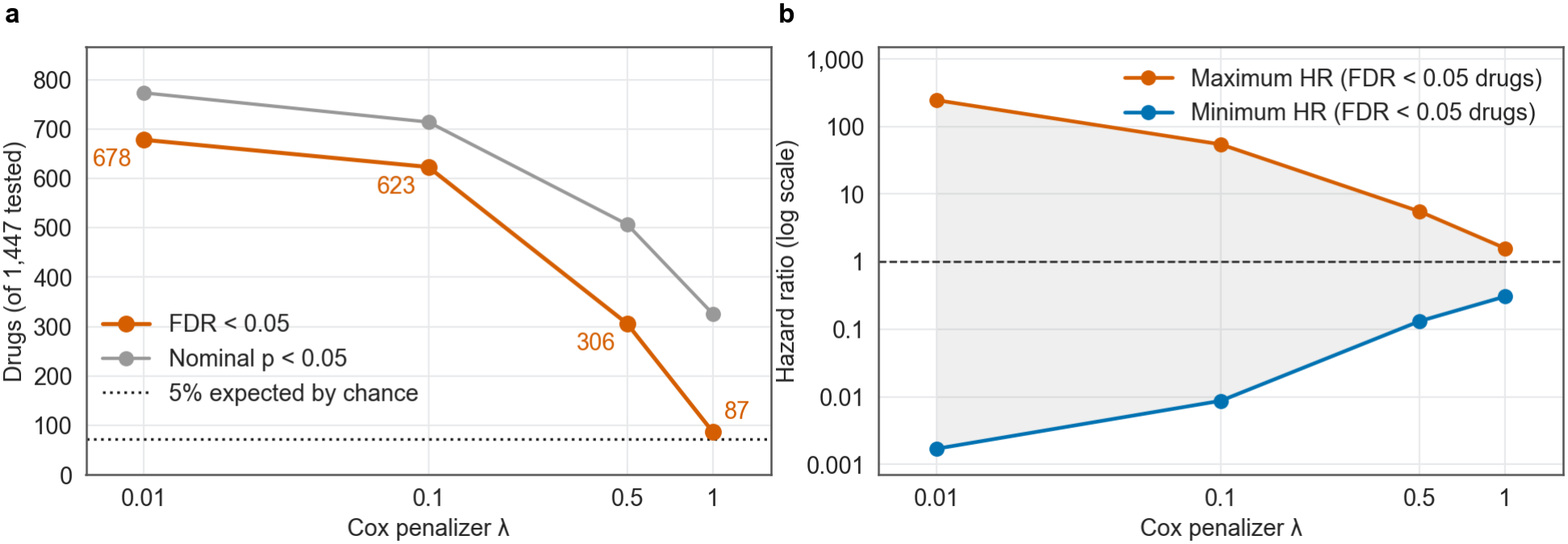
Penalizer sensitivity analysis on all 1,447 drugs in SCAN-B (PAM50-adjusted Cox, one sample per patient). (**a**) Number of drugs with FDR *<* 0.05 and with nominal *p <* 0.05 as a function of the Cox penalizer *λ*, with a dotted line at the 5% of drugs expected at nominal *p <* 0.05 by chance: 678 drugs at FDR *<* 0.05 at *λ* = 0.01, 623 at 0.1, 306 at 0.5 and 87 at 1.0. (**b**) Range of the hazard ratio per unit of predicted AUC among the FDR-significant drugs: [0.002, 245.5] at *λ* = 0.01, [0.009, 54.4] at 0.1, [0.13, 5.53] at 0.5 and [0.31, 1.58] at 1.0. The extreme values come from drugs whose predicted AUC varies little across patients, and per standard deviation of the predicted AUC the significant hazard ratios at *λ* = 0.1 span 0.81 to 1.17. The seven candidates are FDR-significant at *λ* = 0.01, 0.1 and 0.5, and only erlotinib at 1.0.

**Supplementary Figure S12.**
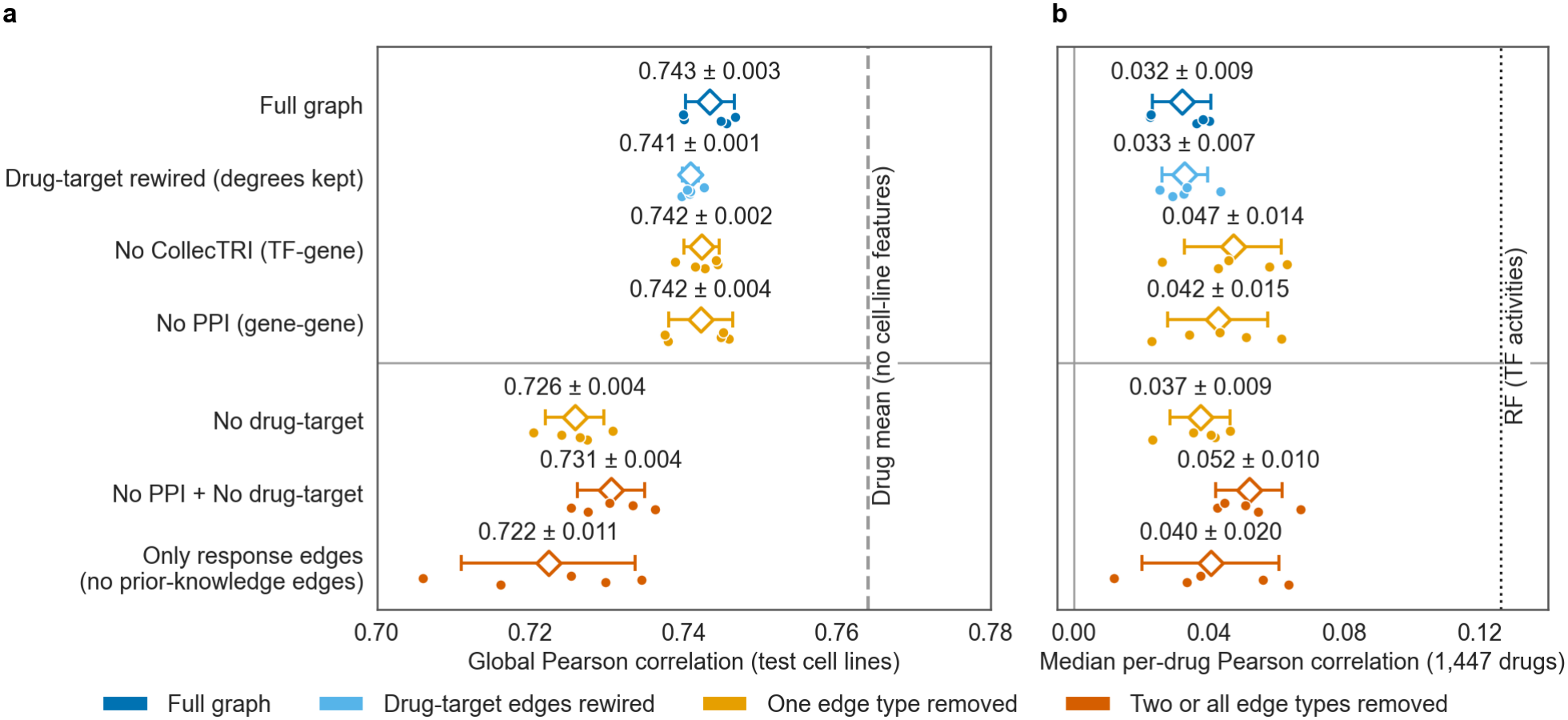
Graph component ablation study (GraphSAGE, seed-42 cell-line split, 800 epochs, final-epoch weights, the protocol of Table 1). (**a**) Global test Pearson correlation. (**b**) Median per-drug test Pearson correlation over the 1,447 drugs. Points show each of the five initialization seeds, spread vertically (seeds with nearly equal values overlap), and open diamonds and bars the mean and standard deviation. The configurations above the horizontal line keep the drug-target relation and those below it do not. The dashed line in a is the drug-mean reference of Table 1 (0.764), which uses no cell-line features and predicts one value per drug, so it has no per-drug correlation. The dotted line in b is the median per-drug correlation of the Random Forest on TF activities (0.126). Removing the PPI or the CollecTRI edges alone barely changes the global correlation (0.742 against 0.743 for the full graph), whereas removing the drug-target edges, the only message-passing relation of the drug nodes, lowers it to 0.726, the level of the two other configurations without drug-target edges (0.731 without PPI and drug-target edges, 0.722 with response edges only). Without drug-target edges no gene or TF node can reach the drug or cell-line nodes, so these three configurations are the same model up to initialization. Every seed of every configuration stays below the drug-mean reference, and keeping the drug-target relation gives a small gain in global correlation, which this comparison cannot attribute to the target annotations because the relation also adds the GraphSAGE layers to every drug node. A control that keeps the relation and rewires its edges at random, with the degree of every drug and gene kept, retains most of that gain (0.741 against 0.743), so most of it does not depend on which genes are annotated as targets. The median per-drug correlation is not higher with prior-knowledge edges (0.032 with the full graph against 0.037 to 0.052 for the reduced graphs), and every configuration stays below the Random Forest.

**Supplementary Figure S13.**
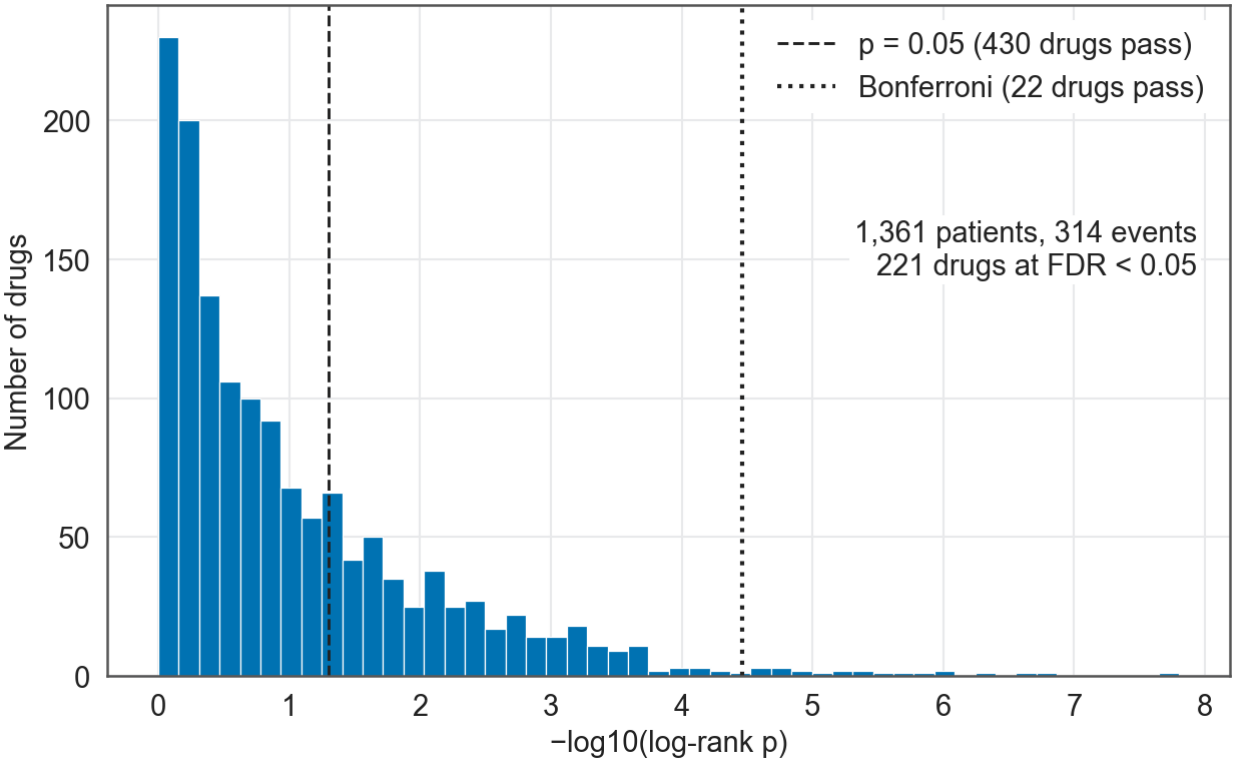
Within-subtype survival screen in SCAN-B, pooled basal-like and HER2-enriched subtypes (n = 1,361 patients, 314 events): distribution of per-drug log-rank p-values (median split of predicted sensitivity), 430 drugs at nominal *p <* 0.05, 221 at FDR *<* 0.05 and 22 after Bonferroni correction. Basal-like alone (n = 686, 167 events) yields 117 drugs. One sample per patient, with the patient rule of the Cox models, restricted to patients with a prediction and overall-survival follow-up.

**Supplementary Figure S14.**
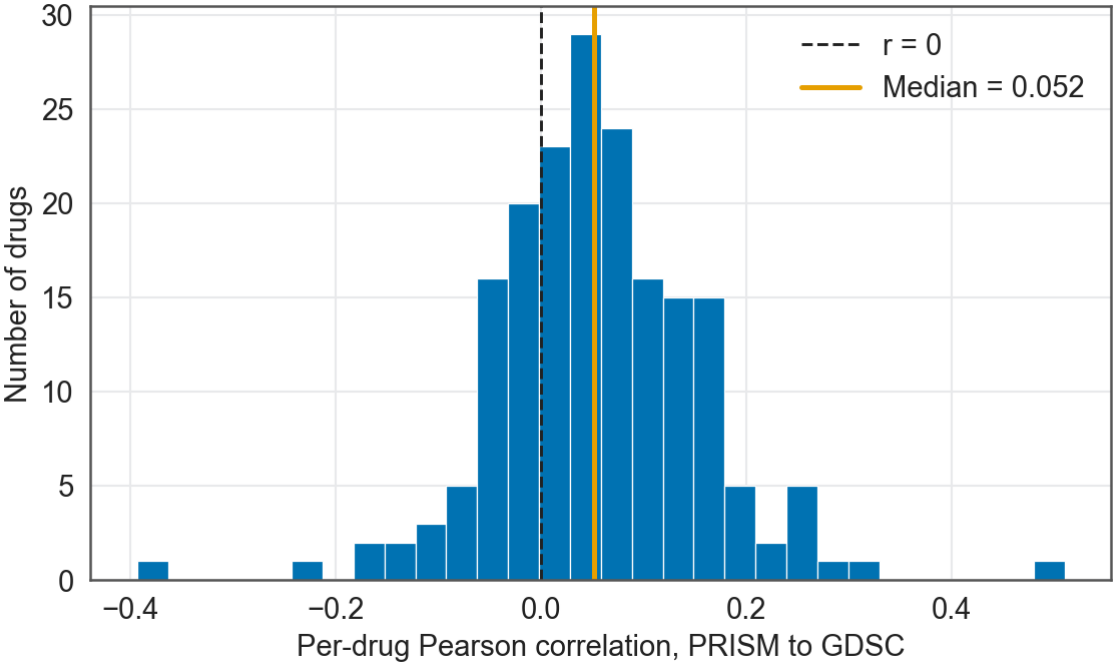
PRISM-to-GDSC cross-screen evaluation of an HGT variant over the 187 drugs shared by both screens, median per-drug Pearson = 0.052. This HGT was trained for 300 epochs (learning rate 10^*−*3^ , no learning-rate schedule, final-epoch weights) on a random 80/20 split of the PRISM response edges (seed 42), and only the 343 GDSC cell lines present in the PRISM graph were evaluated, so every evaluated cell line and drug took part in PRISM training and the correlation is not a cell-line hold-out estimate. The low value may be limited by the agreement between the two screens. The per-drug Ridge models used for clinical transfer were not evaluated across screens.

**Supplementary Table S1.**
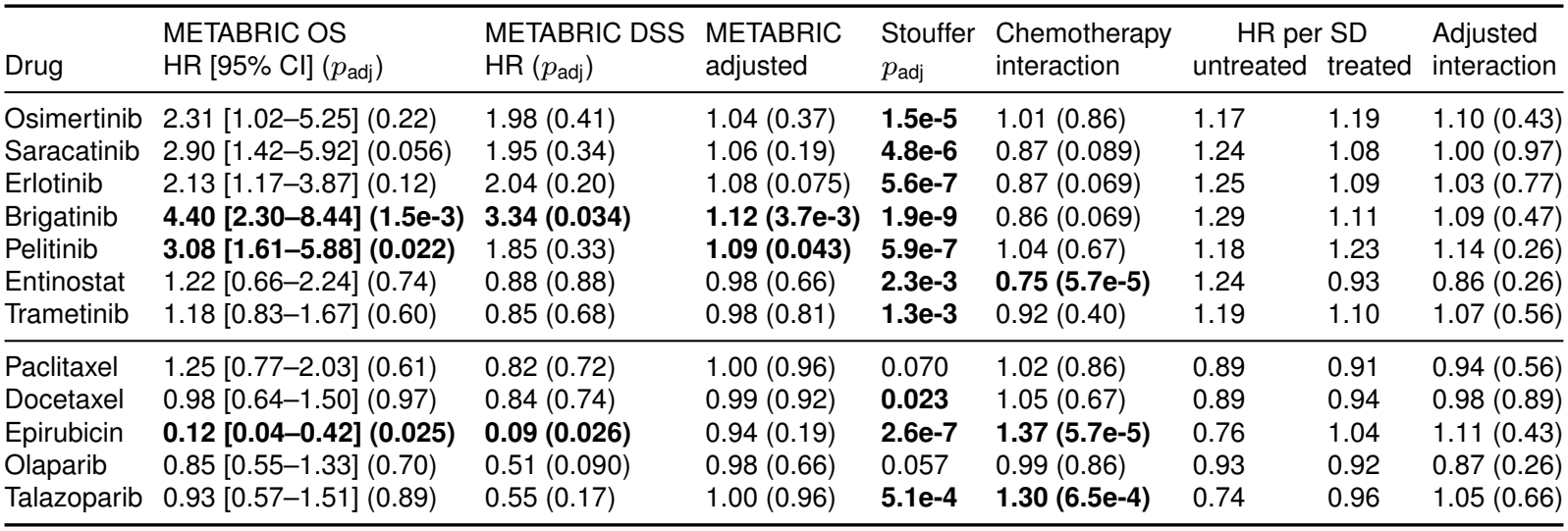
Replication in METABRIC, signed meta-analysis and interaction with chemotherapy for the seven candidates (top) and five positive controls (bottom). METABRIC OS: overall survival (OS), hazard ratio (HR) per unit of predicted AUC in the PAM50-adjusted penalized Cox model of the screen, approximate 95% CI and Benjamini-Hochberg *p*_adj_ over 1,447 drugs. METABRIC DSS: the same model with disease-specific survival (DSS), deaths from other causes censored. METABRIC adjusted: HR per standard deviation (SD) in the unpenalized model with PAM50 as strata and age, positive lymph nodes, ER status, HER2 copy-number gain, the Nottingham Prognostic Index, chemotherapy and hormone therapy as covariates, missing ER or HER2 status as its own category (1,903 patients with complete data, 1,103 events), *p*_adj_ over the 12 drugs. Stouffer: signed meta-analysis of overall survival in the three cohorts, weighted by the square root of the events, *p*_adj_ over 1,447 drugs. Chemotherapy interaction: ratio of the HR per SD of predicted AUC in treated patients to that in untreated patients (exponentiated coefficient of the product of the standardized predicted AUC and chemotherapy receipt) in SCAN-B, with PAM50 as strata (6,843 patients with known chemotherapy status), followed by the HR per SD within untreated and within treated patients. Adjusted interaction: the same model with age, tumor stage, grade, nodal status, ER and PR status and endocrine therapy as main effects (6,173 patients, 946 events). Interaction *p*_adj_ is Benjamini-Hochberg over the 12 drugs. In the METABRIC overall-survival models the score test for proportional hazards (Methods) flagged the prediction terms of six of the 12 drugs (docetaxel, entinostat, epirubicin, erlotinib, pelitinib and talazoparib) and the HER2-enriched, luminal A and luminal B terms of all 12 models (36 of their 48 PAM50 terms). In their refits with PAM50 as strata, it flagged the prediction terms of docetaxel, paclitaxel and talazoparib, all of them controls, in both the penalized and the unpenalized refit. Bold marks *p*_adj_ *<* 0.05.

| Drug | METABRIC OS | METABRIC DSS | METABRIC | Stouffer | Chemotherapy | HR per SD |  | Adjusted |
|---|---|---|---|---|---|---|---|---|
|  | HR [95% CI] (*p*<sub>adj</sub>) | HR (*p*<sub>adj</sub>) | adjusted | *p*<sub>adj</sub> | interaction | untreated | treated | interaction |
| Osimertinib | 2.31 [1.02–5.25] (0.22) | 1.98 (0.41) | 1.04 (0.37) | **1.5e-5** | 1.01 (0.86) | 1.17 | 1.19 | 1.10 (0.43) |
| Saracatinib | 2.90 [1.42–5.92] (0.056) | 1.95 (0.34) | 1.06 (0.19) | **4.8e-6** | 0.87 (0.089) | 1.24 | 1.08 | 1.00 (0.97) |
| Erlotinib | 2.13 [1.17–3.87] (0.12) | 2.04 (0.20) | 1.08 (0.075) | **5.6e-7** | 0.87 (0.069) | 1.25 | 1.09 | 1.03 (0.77) |
| Brigatinib | **4.40 [2.30–8.44] (1.5e-3)** | **3.34 (0.034)** | **1.12 (3.7e-3)** | **1.9e-9** | 0.86 (0.069) | 1.29 | 1.11 | 1.09 (0.47) |
| Pelitinib | **3.08 [1.61–5.88] (0.022)** | 1.85 (0.33) | **1.09 (0.043)** | **5.9e-7** | 1.04 (0.67) | 1.18 | 1.23 | 1.14 (0.26) |
| Entinostat | 1.22 [0.66–2.24] (0.74) | 0.88 (0.88) | 0.98 (0.66) | **2.3e-3** | **0.75 (5.7e-5)** | 1.24 | 0.93 | 0.86 (0.26) |
| Trametinib | 1.18 [0.83–1.67] (0.60) | 0.85 (0.68) | 0.98 (0.81) | **1.3e-3** | 0.92 (0.40) | 1.19 | 1.10 | 1.07 (0.56) |
| Paclitaxel | 1.25 [0.77–2.03] (0.61) | 0.82 (0.72) | 1.00 (0.96) | 0.070 | 1.02 (0.86) | 0.89 | 0.91 | 0.94 (0.56) |
| Docetaxel | 0.98 [0.64–1.50] (0.97) | 0.84 (0.74) | 0.99 (0.92) | **0.023** | 1.05 (0.67) | 0.89 | 0.94 | 0.98 (0.89) |
| Epirubicin | **0.12 [0.04–0.42] (0.025)** | **0.09 (0.026)** | 0.94 (0.19) | **2.6e-7** | **1.37 (5.7e-5)** | 0.76 | 1.04 | 1.11 (0.43) |
| Olaparib | 0.85 [0.55–1.33] (0.70) | 0.51 (0.090) | 0.98 (0.66) | 0.057 | 0.99 (0.86) | 0.93 | 0.92 | 0.87 (0.26) |
| Talazoparib | 0.93 [0.57–1.51] (0.89) | 0.55 (0.17) | 1.00 (0.96) | **5.1e-4** | **1.30 (6.5e-4)** | 0.74 | 0.96 | 1.05 (0.66) |

**Supplementary Table S2.**
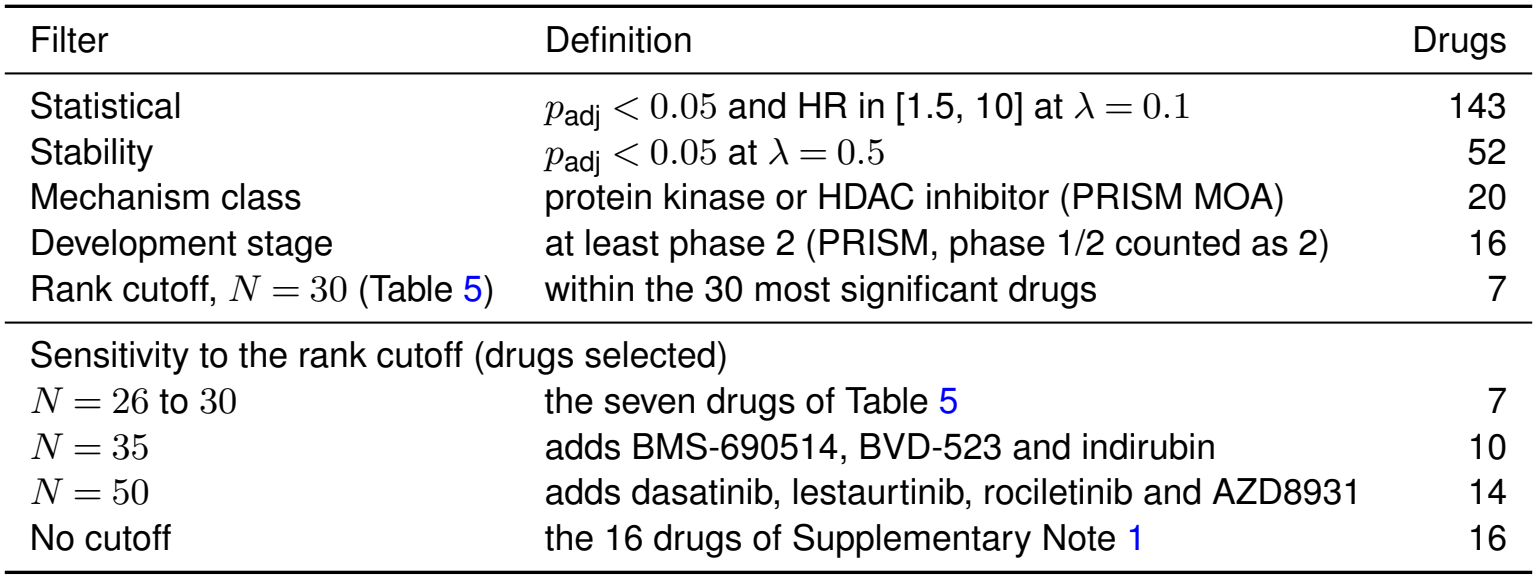
Candidate selection rule (Methods, *Rule-based candidate selection*) applied to the PAM50-adjusted penalized Cox results, and sensitivity of the selected set to the rank cutoff. Filters are applied cumulatively: each row counts the drugs retained after its filter is added to the previous ones. The rank cutoff counts the drugs of the development-stage row that fall within the *N* most significant drugs of the statistical filter. Supplementary Data 1 provides the full list of the 143 drugs passing the statistical filter, with the HR and its 95% confidence interval at *λ* = 0.1, HR and *p*_adj_ at both penalizers, PRISM mechanism of action, molecular target, clinical phase and disease area (release 19Q4, curated by the Broad Drug Repurposing Hub, Corsello et al., 2017), rank, and one indicator per filter.

| Filter | Definition | Drugs |
|---|---|---|
| Statistical | *p*<sub>adj</sub> < 0.05 and HR in [1.5, 10] at *λ* = 0.1 | 143 |
| Stability | *p*<sub>adj</sub> < 0.05 at *λ* = 0.5 | 52 |
| Mechanism class | protein kinase or HDAC inhibitor (PRISM MOA) | 20 |
| Development stage | at least phase 2 (PRISM, phase 1/2 counted as 2) | 16 |
| Rank cutoff, *N* = 30 (Table 5) | within the 30 most significant drugs | 7 |
| Sensitivity to the rank cutoff (drugs selected) |  |  |
| *N* = 26 to 30 | the seven drugs of Table 5 | 7 |
| *N* = 35 | adds BMS-690514, BVD-523 and indirubin | 10 |
| *N* = 50 | adds dasatinib, lestaurtinib, rociletinib and AZD8931 | 14 |
| No cutoff | the 16 drugs of Supplementary Note 1 | 16 |

**Supplementary Table S3.**
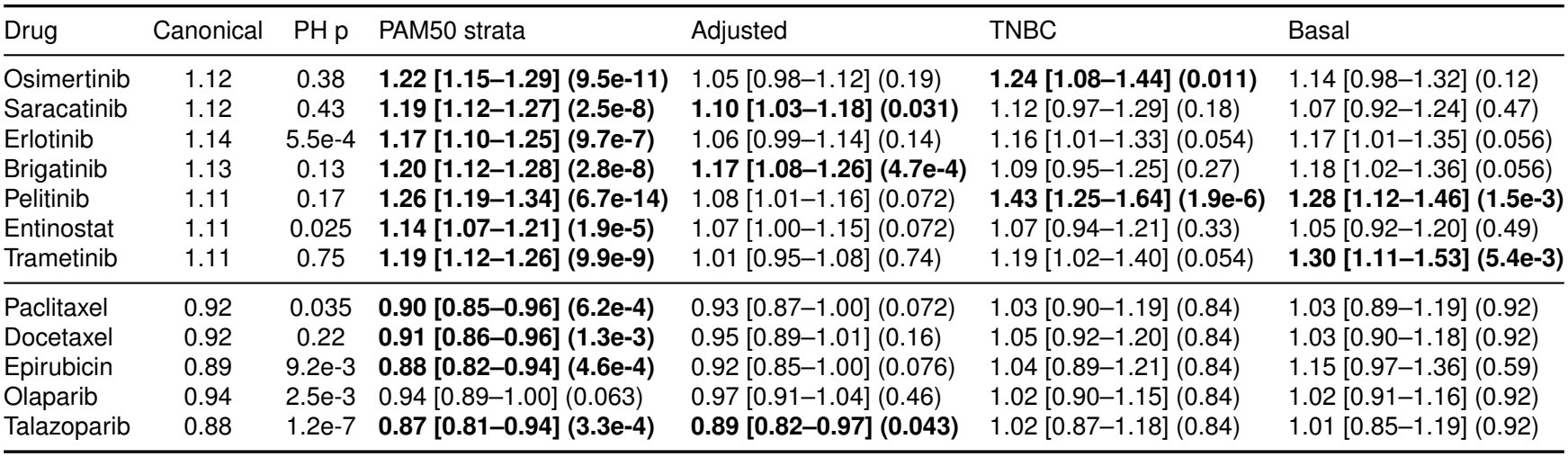
Sensitivity analyses of the survival associations of the seven candidates (top) and five positive controls (bottom) in SCAN-B. All hazard ratios (HR) are per standard deviation of the predicted AUC across patients. Canonical: the PAM50-adjusted penalized Cox model of the screen. PH p: p-value of the score test for proportional hazards (PH) of its prediction term (scaled Schoenfeld residuals), not bolded. PAM50 strata: unpenalized model with PAM50 as strata (7,215 patients, Wald p). Adjusted: unpenalized model with PAM50 as strata and age, tumor stage, grade, nodal status, ER and PR status and the receipt of chemotherapy and endocrine therapy as covariates, missing receptor or treatment status coded as its own category (6,231 patients with complete age, tumor stage, grade and nodal status, 959 events), Benjamini-Hochberg *p*_adj_ over the 12 drugs. TNBC: the 676 patients with ER-, PR- and HER2-negative tumors (183 events). Basal: the 686 PAM50 basal-like patients (167 events). For TNBC and Basal, unpenalized models without PAM50 strata, with Benjamini-Hochberg *p*_adj_ within the candidates and within the controls. Cells give HR [95% CI] (p or *p*_adj_), and bold marks p or *p*_adj_ *<* 0.05 for the association with survival.

| Drug | Canonical | PH p | PAM50 strata | Adjusted | TNBC | Basal |
|---|---|---|---|---|---|---|
| Osimertinib | 1.12 | 0.38 | **1.22 [1.15–1.29] (9.5e-11)** | 1.05 [0.98–1.12] (0.19) | **1.24 [1.08–1.44] (0.011)** | 1.14 [0.98–1.32] (0.12) |
| Saracatinib | 1.12 | 0.43 | **1.19 [1.12–1.27] (2.5e-8)** | **1.10 [1.03–1.18] (0.031)** | 1.12 [0.97–1.29] (0.18) | 1.07 [0.92–1.24] (0.47) |
| Erlotinib | 1.14 | 5.5e-4 | **1.17 [1.10–1.25] (9.7e-7)** | 1.06 [0.99–1.14] (0.14) | 1.16 [1.01–1.33] (0.054) | 1.17 [1.01–1.35] (0.056) |
| Brigatinib | 1.13 | 0.13 | **1.20 [1.12–1.28] (2.8e-8)** | **1.17 [1.08–1.26] (4.7e-4)** | 1.09 [0.95–1.25] (0.27) | 1.18 [1.02–1.36] (0.056) |
| Pelitinib | 1.11 | 0.17 | **1.26 [1.19–1.34] (6.7e-14)** | 1.08 [1.01–1.16] (0.072) | **1.43 [1.25–1.64] (1.9e-6)** | **1.28 [1.12–1.46] (1.5e-3)** |
| Entinostat | 1.11 | 0.025 | **1.14 [1.07–1.21] (1.9e-5)** | 1.07 [1.00–1.15] (0.072) | 1.07 [0.94–1.21] (0.33) | 1.05 [0.92–1.20] (0.49) |
| Trametinib | 1.11 | 0.75 | **1.19 [1.12–1.26] (9.9e-9)** | 1.01 [0.95–1.08] (0.74) | 1.19 [1.02–1.40] (0.054) | **1.30 [1.11–1.53] (5.4e-3)** |
| Paclitaxel | 0.92 | 0.035 | **0.90 [0.85–0.96] (6.2e-4)** | 0.93 [0.87–1.00] (0.072) | 1.03 [0.90–1.19] (0.84) | 1.03 [0.89–1.19] (0.92) |
| Docetaxel | 0.92 | 0.22 | **0.91 [0.86–0.96] (1.3e-3)** | 0.95 [0.89–1.01] (0.16) | 1.05 [0.92–1.20] (0.84) | 1.03 [0.90–1.18] (0.92) |
| Epirubicin | 0.89 | 9.2e-3 | **0.88 [0.82–0.94] (4.6e-4)** | 0.92 [0.85–1.00] (0.076) | 1.04 [0.89–1.21] (0.84) | 1.15 [0.97–1.36] (0.59) |
| Olaparib | 0.94 | 2.5e-3 | 0.94 [0.89–1.00] (0.063) | 0.97 [0.91–1.04] (0.46) | 1.02 [0.90–1.15] (0.84) | 1.02 [0.91–1.16] (0.92) |
| Talazoparib | 0.88 | 1.2e-7 | **0.87 [0.81–0.94] (3.3e-4)** | **0.89 [0.82–0.97] (0.043)** | 1.02 [0.87–1.18] (0.84) | 1.01 [0.85–1.19] (0.92) |

**Supplementary Table S4.**
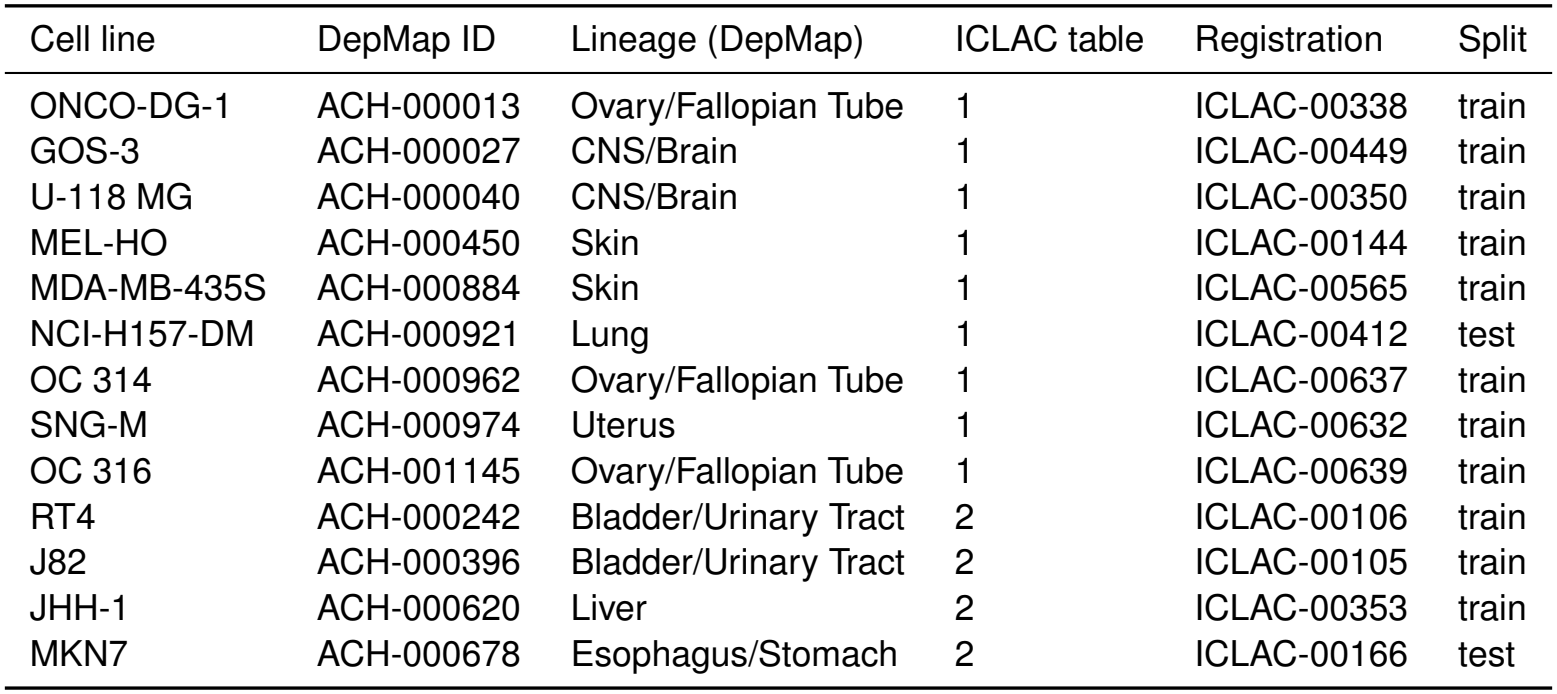
Cell lines of the knowledge graph listed in the ICLAC Register of Misidentified Cell Lines (version 14, 15 February 2026), matched by Cellosaurus accession rather than by name, because name matching flags the parent of a listed derivative. Table 1 of the register lists lines with no known authentic stock, and Table 2 lines for which an authentic stock exists. None of the 12 TNBC lines used for Integrated Gradients is listed. All 13 lines were retained, because the models receive no tissue or lineage label as input. Split refers to the cell-line hold-out drawn with seed 42.

**Supplementary Table S5.**
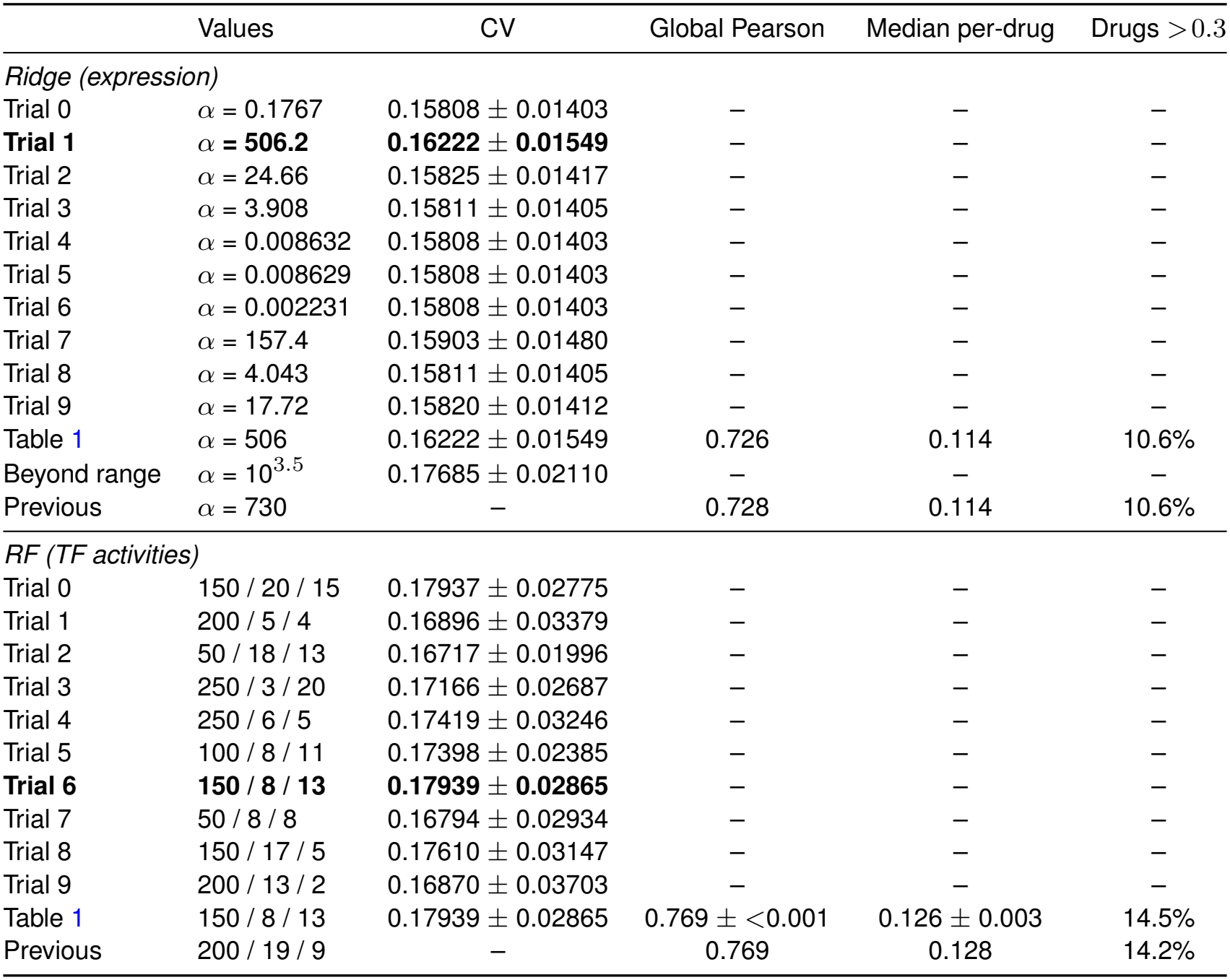
Hyperparameter search of Ridge on expression and the Random Forest on TF activities of Table 1. Ten configurations per model were drawn at random from the search space (Optuna, seed 42) and scored by five-fold cross-validation within the 381 training cell lines, with the lines of each donor in the same fold and the folds stratified by lineage. CV: mean *±* standard deviation over the folds of the median per-drug Pearson correlation over the 200 drugs with most training measurements. Random Forest values are trees / maximum depth / minimum samples per leaf. Bold marks the best configuration of each model. Table 1: the values used in that table, with the *α* of Ridge rounded to three significant figures and its CV recomputed, and their test metrics (Random Forest, mean *±* standard deviation over five seeds). Beyond range: because the best *α* lay near the upper end of the range, one value beyond it was scored as a diagnostic and not evaluated on the test lines, and, as fixed before the search was run, Table 1 keeps the value chosen within the range. Previous: the values of version 1 of the preprint of this work (Methods), chosen on the 95 test lines, with the test metrics they gave. The two best Random Forest configurations scored almost the same.

|  | Values | CV | Global Pearson | Median per-drug | Drugs >0.3 |
|---|---|---|---|---|---|
| *Ridge (expression)* |  |  |  |  |  |
| Trial 0 | α = 0.1767 | 0.15808 ± 0.01403 | – | – | – |
| **Trial 1** | α **= 506.2** | **0.16222** ± **0.01549** | – | – | – |
| Trial 2 | α = 24.66 | 0.15825 ± 0.01417 | – | – | – |
| Trial 3 | α = 3.908 | 0.15811 ± 0.01405 | – | – | – |
| Trial 4 | α = 0.008632 | 0.15808 ± 0.01403 | – | – | – |
| Trial 5 | α = 0.008629 | 0.15808 ± 0.01403 | – | – | – |
| Trial 6 | α = 0.002231 | 0.15808 ± 0.01403 | – | – | – |
| Trial 7 | α = 157.4 | 0.15903 ± 0.01480 | – | – | – |
| Trial 8 | α = 4.043 | 0.15811 ± 0.01405 | – | – | – |
| Trial 9 | α = 17.72 | 0.15820 ± 0.01412 | – | – | – |
| Table 1 | α = 506 | 0.16222 ± 0.01549 | 0.726 | 0.114 | 10.6% |
| Beyond range | α = 10<sup>3.5</sup> | 0.17685 ± 0.02110 | – | – | – |
| Previous | α = 730 | – | 0.728 | 0.114 | 10.6% |
| *RF (TF activities)* |  |  |  |  |  |
| Trial 0 | 150 / 20 / 15 | 0.17937 ± 0.02775 | – | – | – |
| Trial 1 | 200 / 5 / 4 | 0.16896 ± 0.03379 | – | – | – |
| Trial 2 | 50 / 18 / 13 | 0.16717 ± 0.01996 | – | – | – |
| Trial 3 | 250 / 3 / 20 | 0.17166 ± 0.02687 | – | – | – |
| Trial 4 | 250 / 6 / 5 | 0.17419 ± 0.03246 | – | – | – |
| Trial 5 | 100 / 8 / 11 | 0.17398 ± 0.02385 | – | – | – |
| **Trial 6** | **150** / **8** / **13** | **0.17939** ± **0.02865** | – | – | – |
| Trial 7 | 50 / 8 / 8 | 0.16794 ± 0.02934 | – | – | – |
| Trial 8 | 150 / 17 / 5 | 0.17610 ± 0.03147 | – | – | – |
| Trial 9 | 200 / 13 / 2 | 0.16870 ± 0.03703 | – | – | – |
| Table 1 | 150 / 8 / 13 | 0.17939 ± 0.02865 | 0.769 ± <0.001 | 0.126 ± 0.003 | 14.5% |
| Previous | 200 / 19 / 9 | – | 0.769 | 0.128 | 14.2% |

**Supplementary Table S6.** Patients of the survival analyses. Patients, events, sex, age and subtypes refer to the patients of the Cox models, the PAM50-adjusted models of SCAN-B, METABRIC and TCGA-BRCA (Table 2) and the relapse-free survival models of FUSCC-TNBC. Follow-up is the reverse Kaplan-Meier median over the patients with survival data, which in SCAN-B (7,397) and TCGA-BRCA (1,072) include those without a PAM50 subtype. Age is the age at diagnosis in METABRIC, at the index date in TCGA-BRCA and at surgery in FUSCC-TNBC. SCAN-B distributes age in five-year bands, so its values are band values, and has no sex variable. Nine TCGA-BRCA ages are obfuscated by the Genomic Data Commons. FUSCC-TNBC distributes the intrinsic subtype only as basal-like or other. AURORA-US, which enters no survival analysis, is described in Methods. IQR, interquartile range.

|  | SCAN-B | METABRIC | TCGA-BRCA | FUSCC-TNBC |
|---|---|---|---|---|
| Expression platform | RNA-seq | Microarray | RNA-seq | RNA-seq |
| Endpoint | Overall survival | Overall survival | Overall survival | Relapse-free survival |
| Patients | 7,215 | 1,979 | 1,060 | 356 |
| Events | 1,164 | 1,143 | 149 | 49 |
| Median follow-up (years) | 7.0 | 16.3 | 2.7 | 4.0 |
| Women | – | 1,979 | 1,060 | 356 |
| Age (years), median [IQR] | 65 [55–75] | 61.8 [51.4–70.6] | 58 [48–67] | 53.5 [45–61] |
| Age not recorded | 126 | 0 | 0 | 0 |
| *PAM50 subtype* |  |  |  |  |
| Luminal A | 3,569 | 569 | 553 | – |
| Luminal B | 1,749 | 823 | 204 | – |
| HER2-enriched | 675 | 254 | 77 | – |
| Basal-like | 686 | 256 | 187 | 275 |
| Normal-like | 536 | 77 | 39 | – |
| Other | – | – | – | 81 |

## Supplementary Data

**Supplementary Data 1**. Drugs passing the statistical filter of the candidate selection, one row per drug, with the columns described in Supplementary Table S2.

## Supplementary Notes

### Supplementary Note 1: Integrated Gradients on the extended candidate set

To test whether the attributions depend on the rank cut-off of the selection rule, we repeated the Integrated Gradients analysis with the protocol of the main text (seed-42 model for the global ranking, models of seeds 42, 123 and 456 for seed stability, mean-train base-line, 50 steps, the same 12 TNBC cell lines) on an extended set of 20 drugs instead of 11: the 16 kinase and HDAC inhibitors that pass the statistical, stability, mechanism-class and development-stage filters of Supplementary Table S2 without the rank cutoff, plus the four positive controls of the main analysis (paclitaxel, docetaxel, epirubicin and olaparib, without talazoparib). The global TF ranking was essentially unchanged: 19 of the 20 most important TFs were the same in both runs (HIVEP1 only in the main analysis, CTBP1 only in the extended one), and the two global importance profiles over the 771 TFs correlated with Spearman *ρ* = 0.9995. The ten core TFs were identical in both runs, and the AURORA-US primary-versus-metastasis test gave the same five Bonferroni-significant TFs (KLF8, STAT2, AEBP1, SPDEF and SETBP1). The main-text attributions therefore do not depend on the rank cut-off. The near-identity of the global profiles indicates that mean attribution magnitude is driven by TF effects shared across drugs, consistent with the shared sign profile of Figure 3c. The comparison metrics, core sets and extended AURORA-US test are provided with the code.

### Supplementary Note 2: Sensitivity to how the TF activities were computed

The knowledge graph and the PRISM and GDSC TF activities were computed from a CollecTRI snapshot cached on 28 March 2026 (42,990 signed edges, 1,185 TFs). The clinical TF-activity matrices (SCAN-B, METABRIC, TCGA-BRCA) were computed in April 2026 with a CollecTRI network retrieved separately through decoupleR, whose regulons differed for a subset of TFs. Recomputing the three clinical matrices with the snapshot reproduced the activities of most TFs exactly and changed 125 (SCAN-B), 119 (METABRIC) and 137 (TCGA-BRCA) of them. Of these, 26, 30 and 39 had a Pearson correlation below 0.99 against the published values. Rerunning the multi-cohort survival screen on the recomputed matrices gave 604 instead of 623 significant drugs in SCAN-B (574 shared), 87 instead of 74 in METABRIC (71 shared), none in TCGA-BRCA and 558 instead of 551 in the Fisher meta-analysis (525 shared), with 59 drugs significant in both SCAN-B and METABRIC, all with the same hazard-ratio direction. The four positive controls that reach significance in the Fisher meta-analysis remained significant (doc-etaxel at adjusted p = 0.0499), and olaparib remained non-significant. The candidate selection rule yielded 154, 56, 18, 15 and 7 drugs at its five steps, with six of the seven Table 5 candidates unchanged: indirubin (a CDK and GSK3 inhibitor in the PRISM annotation, HR = 8.8) replaced trametinib, which ranked 35th and fell just outside the rank cutoff. The AURORA-US analysis, computed in Python with the same snapshot, was reproduced exactly. The aggregate counts and the positive controls changed little with the network version, but one of the seven candidates did (indirubin replaced trametinib), and 574 of the 623 SCAN-B associations remained significant. The recomputed matrices and every count of this note are produced by the accompanying code (see Code availability).

The same step carries a second dependency, the gene universe each cohort brings. The univariate linear model behind a TF activity grows with the number of genes in the matrix, so cohorts quantified over different gene sets produce activities on different scales, and the quantile transformation of the clinical transfer, fitted on the PRISM activities, clips whatever falls outside the range it saw in training. 12.7% of the SCAN-B activities and 12.3% of the METABRIC ones fall outside that range, against 80.3% of the TCGA-BRCA ones, which were quantified over the much larger gene set of the GDC release. Recomputing the TCGA activities over the 17,170 genes shared with the PRISM expression matrix leaves 11.8% outside, widens the median spread of the predicted AUC from 0.0824 to 0.1315 and moves the genomic-control inflation of the cohort from 0.84 to 1.03, but it does not change what the cohort contributes: TCGA-BRCA still has no drug at FDR *<* 0.05, its share of nominally significant drugs stays at or below chance (34 and 46 of the 1,447 drugs, both below the 5% expected by chance), and the Fisher meta-analysis moves from 551 to 559 drugs, 525 of them shared, with every positive control keeping its status (paclitaxel adjusted p 8.6 *×*10^*−*3^ against 1.2 *×*10^*−*2^, docetaxel 3.1 *×*10^*−*2^ against 3.5*×* 10^*−*2^, and olaparib above 0.05 either way). The published cohort results are therefore kept as they are, and the comparison is reproduced by the accompanying code. No PAM50 term of the TCGA-BRCA models of the candidates and controls departed from proportional hazards with either version of the activities, and two of the twelve drug terms did in each (olaparib and osimertinib with the published activities, brigatinib and talazoparib with those computed over the PRISM gene universe). FUSCC-TNBC leaves 71.4% of its activities outside that range when it is quantified over its full transcriptome and 13.9% over the same PRISM gene universe, which is why it is computed that way from the start.

